# Governing principles of transcriptional logic out of equilibrium

**DOI:** 10.1101/2023.11.20.567770

**Authors:** Smruti Dixit, Teije C. Middelkoop, Sandeep Choubey

## Abstract

Cells face a myriad of environments and signaling cues. In order to survive, adapt, and desvelop, cells respond to external and internal stimuli, by tightly regulating transcription. Transcriptional regulation involves combinatorial binding of a repertoire of transcription factors (TFs) to DNA, which often results in switch-like binary outputs, akin to logic gates. Recent experimental studies have demonstrated that transcription factor binding to DNA often involves energy expenditure, thereby driving the system out of equilibrium. The governing mechanistic principles of transcriptional logic out of equilibrium remain elusive. To this end, we employ a simple two-input model of transcription that explicitly considers the non-equilibrium binding of TFs to DNA. This simple model can accommodate both equilibrium and non-equilibrium mechanisms and allows for a comparative study of logic operations obtained in the two regimes. We find that, out of equilibrium, the regulatory function of two transcription factors gets altered in a mutually exclusive manner. Such behavior allows non-equilibrium regimes to recreate all the logic operations seen in equilibrium and create new logic operations inaccessible in equilibrium. Our findings demonstrate that cells attain a wider range of decision-making abilities by expending energy.

## 1 Introduction

Cells exhibit a wide variety of gene expression outcomes in response to both extrinsic and intrinsic cues. These genetic responses are controlled by the binding of transcription factors (TFs) to the promoter, a region of DNA upstream of a gene, thereby either activating or repressing transcriptional output. By integrating the activity of multiple TFs at the promoter, cells can convert analog TF inputs into binary transcriptional outputs, [1–6]. Unraveling the mechanistic principles of such logical computation at the level of transcriptional is a central challenge in the field of regulatory biology.

The so-called thermodynamic models of transcription have shed light on how cis-regulatory elements integrate different combinations and concentrations of TFs to regulate gene expression. These models have revealed that TF-promoter interaction and cooperativity between TFs allow for implementing different logical computations in bacteria [5, 7–10]. Similar models have been employed to engineer synthetic logic gates in eukaryotes [11]. In these models, binding of transcription factors to the promoter is assumed to happen in thermodynamic equilibrium, where TF binding obeys the condition of detailed balance and the net probability fluxes in the system vanish. Consequently, such a system does not expend any energy. Equilibrium descriptions have been particularly successful in explaining the input-output relationship of genes in bacteria [12–14]. However, an experimental study showed that repression in the widely studied lac operon in bacteria is quantitatively inconsistent with an equilibrium model [15]. On the other hand, various steps in gene regulation in eukaryotes involve energy expenditure and therefore operate out of equilibrium. For example, nucleosome movement, post-translational histone modifications, and the assembly of transcriptional machinery all require ATP consumption [16, 17].

Recent years have seen a growing interest in characterizing gene regulation out of equilibrium and dissecting its physiological implications. For instance, a recent study showed that a thermodynamic model of gene regulation fails to explain the sharp expression of a developmental gene, called hunchback, in Drosophila [18]. The authors proposed a simple non-equilibrium model of TF binding to the promoter to explain the experimental findings, akin to the model of kinetic proofreading, as developed by John Hopfield [19, 20]. In this model, the detailed balance condition was assumed to be broken during TF binding, leading to energy expenditure. Subsequent theoretical studies have shown that non-equilibrium allows for TF binding specificity [21–23], leads to accelerated information transmission during transcription [24], and enhanced sensitivity to TF concentration [25, 26]. Intriguingly, another theoretical study showed that, when operating out of equilibrium, a single transcription factor can show a non-monotonic response curve [26]. Nonetheless, the governing principles of transcriptional logic out of equilibrium remain elusive.

Here we seek to develop a mechanistic understanding of how cis-regulatory elements integrate different TF inputs to carry out logical computation out of equilibrium. To this end, we use a simple model of two-input transcriptional logic. This model can accommodate both equilibrium and non-equilibrium mechanisms of TF binding to the promoter. We show that the same wide range of logical computations, previously ascribed to the equilibrium binding of TFs to the promoter, can also be implemented out of equilibrium. Interestingly, energy expenditure can alter the simplistic notion of activators and repressors; in some parameter regimes, an activator can behave as a repressor and vice versa. Consequently, novel logical computational capabilities emerge out of equilibrium. Our study demonstrates that moving out of equilibrium can significantly expand the logical computational capacity of cells.

## 2 Results

### 2.1 A simple model of two-input transcriptional logic

To decipher the principles of transcriptional logic out of equilibrium, we consider a simple model of gene regulation, wherein two transcription factors regulate the expression of a gene, as shown in Fig.1. This model has been employed widely to gain mechanistic insights into the regulation of gene expression [11, 27–29]. In this model, the promoter consists of two binding sites to which the two transcription factors, A and B can bind. The promoter can exist in four different states-empty (state 1), TF A bound (state 2), TF A and B bound (state 3), and TF B bound (state 4). The promoter can switch between different states, either through the binding or the unbinding of a protein, with the rates detailed in Fig.1. The rate of mRNA production from promoter state *i* is given by *r*_*i*_. An mRNA molecule degrades at a constant rate *γ*. The inputs to the model are the concentrations of TF A and TF B, and the output is the average mRNA level. We consider two scenarios: first, when TF A and TF B act as activators, we assume *r*_1_ = 0 while *r*_2_ = *r*_3_ = *r*_4_ = *r* that is, transcription commences at the same rate when A or B or both are bound to the promoter. Second, when TF A and TF B act as repressors, we assume that *r*_2_ = *r*_3_ = *r*_4_ = 0 and *r*_1_ = *r* [27], where the binding of either protein to the promoter blocks transcription. The binding rates (*k*_1_, *k*_2_, *k*_−3_, *k*_−4_) have units of *µM* ^−1^*s*^−1^, the unbinding rates (*k*_−1_, *k*_−2_, *k*_3_, *k*_4_) have units of *s*^−1^ while the concentrations of A ([*A*]) and B ([*B*])have units of *µM* [7, 11, 30].

**Figure 1.**
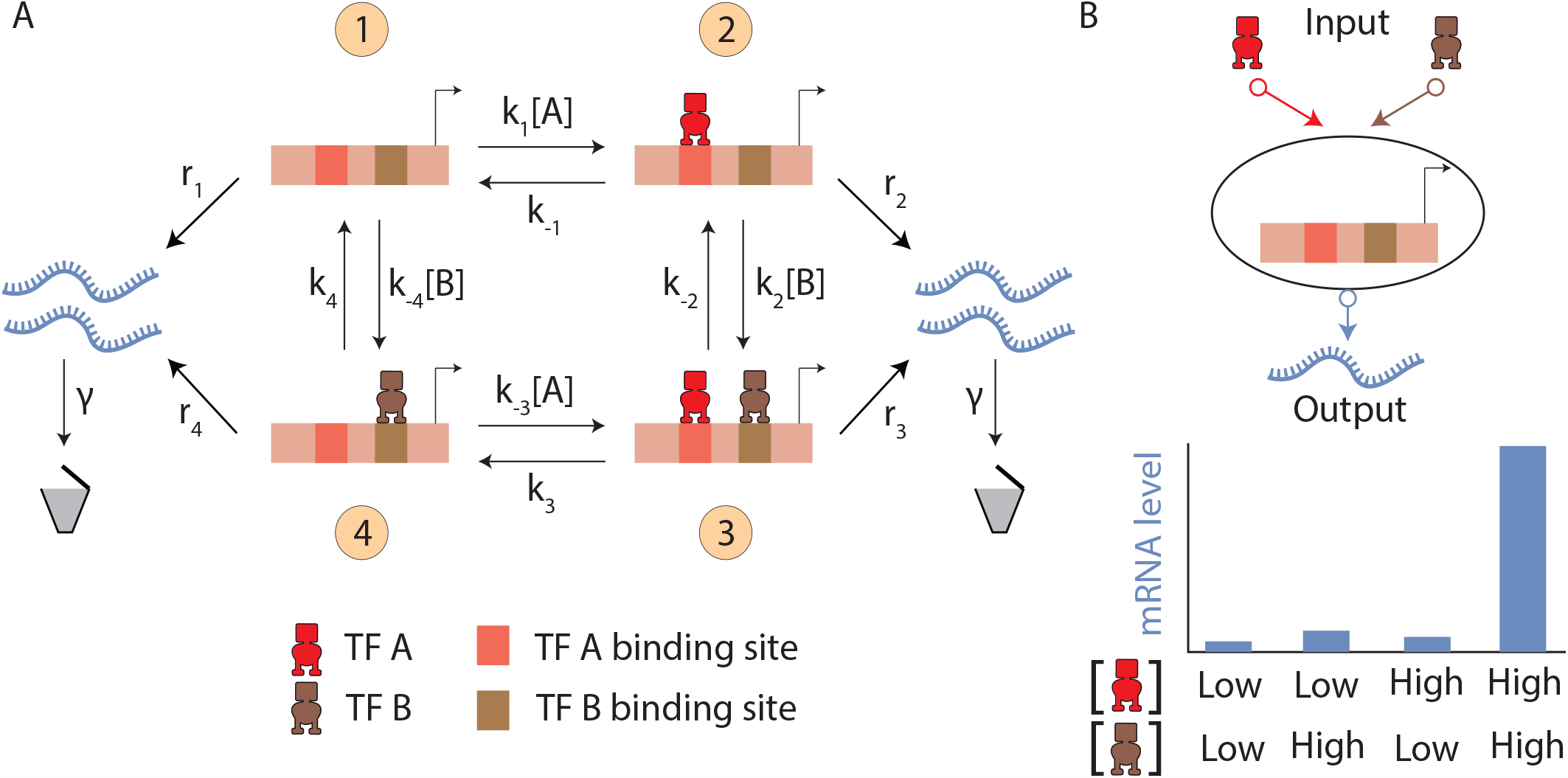
A simple model of 2 input transcriptional logic: A) The promoter can bind two transcription factors A (red) and B (brown). There are four possible states of the promoter, depicted as states 1,2,3 and 4. The promoter can switch between these states, described by the four binding rates (*k*_1_, *k*_2_, *k*_*−*3_, *k*_*−*4_) and unbinding rates (*k*_*−*1_, *k*_*−*2_, *k*_3_, *k*_4_) of the two transcription factors. Rate of transcription from each state *i* is denoted by *r*_*i*_, rate of degradation of an mRNA transcript is *γ*. B) A depiction of the transcriptional logic gate. The concentrations of the transcription factors act as inputs and the steady-state mRNA levels are the output, shown below is an example of a representative logical computation, mimicking an AND gate.

The steady-state probability for the promoter to occupy the *i*-th state is denoted by *P*_*i*_. *P*_*i*_s are a function of the concentrations of the two input TFs and the rate of switching between different promoter states. In line with previous studies, we assume that the average mRNA level at steady state is proportional to promoter occupancies in transcriptionally active states [18, 27]. Hence, the average mRNA level, ⟨*m*⟩ is given by.

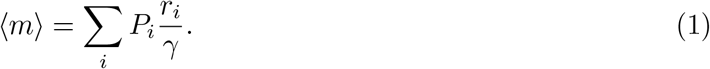

It is important to note that the average mRNA level is governed primarily by the promoter state occupancies. Hence the key to unraveling the principles of building logic gates lies in understanding how the promoter state occupancies are tuned as functions of input TF concentrations and the rates of promoter switching.

We compute promoter state occupancies using the Matrix Tree Theorem [31] (see SI 1). First, the weights of each of the four promoter states are given by

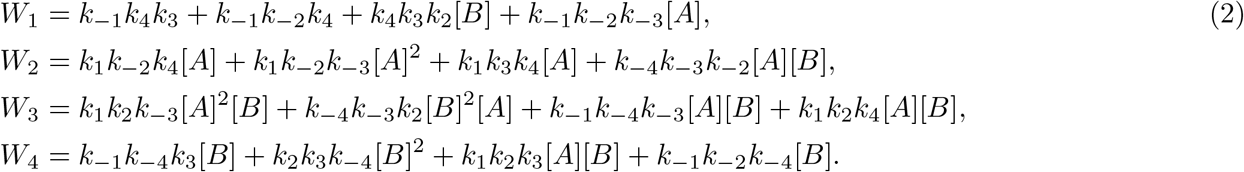

Next, we find the steady-state occupancies of the different promoter states by normalizing the weights, as computed from Equation 2, given by

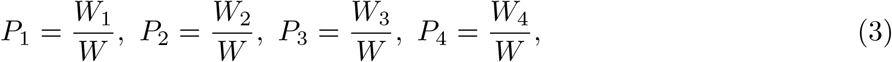

where *P*_*i*_ is the steady-state occupancy of the *i*-th promoter state, and

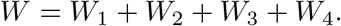

Different limits of our model exhibit equilibrium and non-equilibrium behavior. In equilibrium, the steady state of the system obeys the condition of detailed balance. The detailed balance condition states that for any two states *i* and *j* in the cycle shown in Fig.1, the probability flux from state *i* to *j* is balanced by an equal flux from state *j* to *i*. That is, *P*_*i*_*k*_*ij*_ = *P*_*j*_*k*_*ji*_, where *P*_*i*_ and *P*_*j*_ are the steady-state occupancies of states *i* and *j* respectively, *k*_*ij*_ is the rate of transition from state *i* to *j* and *k*_*ji*_ is the rate of transition from state *j* to *i*. For a cyclic system, such as Fig.1, the condition of detailed balance is captured by

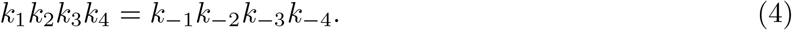

This condition is referred to as the Kolmogorov criterion for a cycle. The system is in equilibrium if the promoter switching rates satisfy this condition [31, 32]. Hence in equilibrium, the weights of the promoter states are,

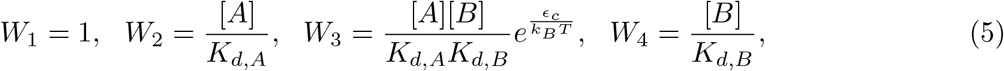

where *K*_*d,A*_ and *K*_*d,B*_ are the dissociation constants for TF A and TF B in *µM*, and *ϵ*_*c*_ is the cooperative interaction energy between the two proteins in *k*_*B*_*T* (see SI 1). These parameters are related to the promoter switching rates in the following manner:

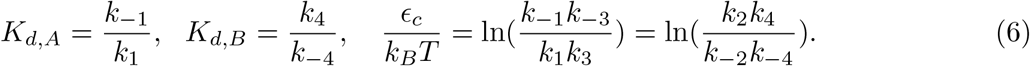

When the Kolmogorov criterion does not hold, the steady-state occupancies no longer obey the detailed balance condition and as a result, the system operates out of equilibrium. In this case, the steady-state occupancies are maintained by a steady-state probability flux along one of the directions of the cycle [32]. Unlike the case of equilibrium circuits, maintaining a non-equilibrium steady state requires an external energy source, such as a chemical potential gradient, and such a system generates heat in the process. The dissipated energy per cycle in the anticlockwise direction is given by [32–34].

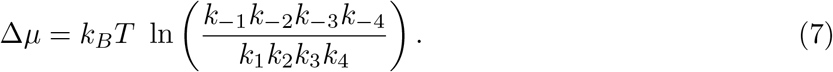

The sign of Δ*µ* informs about the direction of the steady state probability flux. A positive value indicates that the net flux around the cycle is anticlockwise whereas a negative value indicates that the net flux around the cycle is clockwise. It must be noted that this quantity is independent of the concentration of TFs A and B.

For a given set of promoter switching rates and input TF concentrations, we find the promoter state occupancies using Equations 2 and 3. We then compute the average mRNA level using Equation 1. We compute the average mRNA level for a set of four inputs; when [*A*] and [*B*] are low ([*A*]_*Low*_, [*B*]_*Low*_), when [*A*] is low and [*B*] is high ([*A*]_*Low*_, [*B*]_*High*_), when [*A*] is high and [*B*] is low ([*A*]_*High*_, [*B*]_*Low*_) and when both [*A*] and [*B*] are high ([*A*]_*High*_, [*B*]_*High*_). Afterward, we compare the mRNA output of the model to that of an ideal logic gate.

### 2.2 A measure of goodness of a logic gate

To unravel the principles behind building logic gates, we quantify how closely the output of our model resembles that of the target logic gate. We employ the *l*_1_ norm to compare the output of the model to that of an ideal gate. First, we normalize the average mRNA over all the inputs and then compare it to the normalized output of an ideal gate. The *l*_1_ norm is then given by

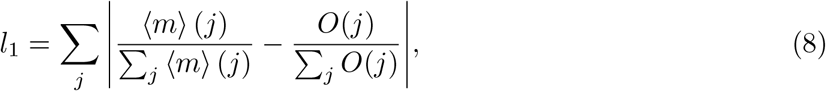

where *j* = {([*A*]_*Low*_, [*B*]_*Low*_), ([*A*]_*Low*_, [*B*]_*High*_), ([*A*]_*High*_, [*B*]_*Low*_), ([*A*]_*High*_, [*B*]_*High*_)} is the set of inputs and *O*(*j*) is the output of an ideal gate for each of these inputs which takes the values 0 and 1.

The *l*_1_ norm is bounded between 0 and 2, with a lower value indicating a better match to the target gate. It has the intuitive interpretation of adding the errors, where each term in the sum measures the extent to which the system’s output for a given input differs from the ideal gate. We define an ‘optimum gate’ as a circuit with an *l*_1_ norm below 0.1. We also take into account the average mRNA level when the transcriptional output is in the ‘high’ state. This is because cells need a certain number of mRNA transcripts to carry out their functions [30, 35, 36]. We reject all optimum gates with a low mRNA count (see Materials and Methods for details). This constraint imposed on the average mRNA level along with the cutoff imposed on the *l*_1_ norm does not change the qualitative results presented in this manuscript (see SI 6 and 7).

Having established all the necessary ingredients to carry out our analysis, we begin with a systematic dissection of the promoter occupancies in and out of equilibrium, as described in the ensuing section.

### 2.3 Promoter state occupancies exhibit non-monotonic behavior out of equilibrium

Building optimal logic gates entails a detailed understanding of how average mRNA level changes as a function of input TF concentrations. Since promoter state occupancies dictate the average mRNA level, it is imperative that we first characterize promoter state occupancies in and out of equilibrium. For convenience, we define a reference value of [*A*]_0_ = [*B*]_0_ = 1*µM* to logarithmically sample the dimensionless concentrations of TF A ([*A*]*/*[*A*]_0_) and TF B ([*B*]*/*[*B*]_0_). For reference, we first take an equilibrium circuit where both A and B have the same dissociation constants (i.e. *K*_*d,A*_ = *K*_*d,B*_). Furthermore, we assume that there is no cooperative interaction between the TFs (*ϵ*_*c*_ = 0). In equilibrium, promoter state occupancies behave monotonically with one of the TF concentrations when the concentration of the second protein remains constant as shown in Fig.2 A, B (SI 1). The introduction of cooperativity (*ϵ*_*c*_ *>* 0) leads to an increase in the area of the interface between *P*_1_ and *P*_3_, and increases *P*_3_ in regions of lower [*A*] and [*B*] as shown in Fig.2 B. This allows for a sharp transition from state 1 into state 3 with an increase in [*A*] or [*B*].

**Figure 2.**
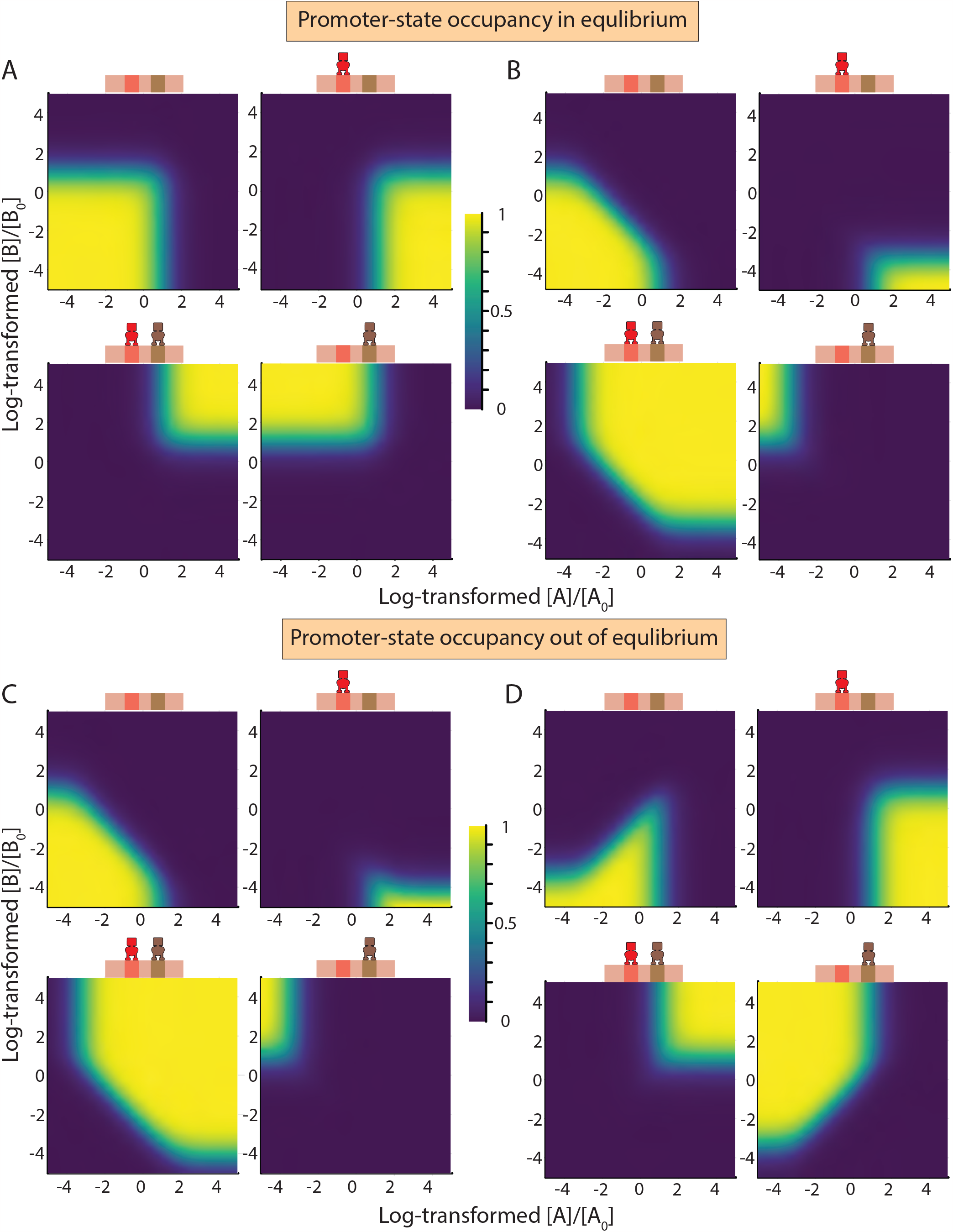
Promoter occupancies in and out of equilibrium: Promoter state occupancies as a function of log_10_ ([*A*]*/*[*A*]_0_) (x-axis) and log_10_ ([*B*]*/*[*B*]_0_)(y-axis), shown as a heatmap with the accompanying color bars ranging from 0 to 1. A) Promoter occupancies at equilibrium, no cooperativity, B) Promoter occupancies at equilibrium, 10*k*_*B*_ *T* of coopera-tivity, C) Promoter occupancies out of equilibrium, 2 *k*_*B*_ *T* of energy dissipation, D) Promoter occupancies out of equilibrium, 10 *k*_*B*_ *T* of energy dissipation. For parameter values see Materials and Methods Table 2.

Next, we explore occupancy profiles of different promoter states out of equilibrium using the Kolmogorov criterion. This criterion divides the eight-dimensional space spanned by promoter state switching rates into two mutually exclusive and complementary subsets. One consists of the set of rates satisfying the Kolmogorov criterion and describing an equilibrium circuit, the second set violates the criterion and describes circuits operating out of equilibrium. Such a clear distinction between equilibrium and non-equilibrium circuits does not arise at the level of the promoter state occupancies (see Fig.2 B, C). This is because each of the *P*_*i*_s is a well-defined and continuous function of the various rates. This continuity ensures that a small change in one or more of the promoter switching rates changes the occupancy of the different promoter states by only a small magnitude. While such a change drives the system out of equilibrium, the promoter state occupancy profiles do not get altered appreciably. Hence, non-equilibrium circuits can mimic equilibrium circuits, with small amounts of energy dissipation, as shown in Fig.2 B, C.

**Table 1:**
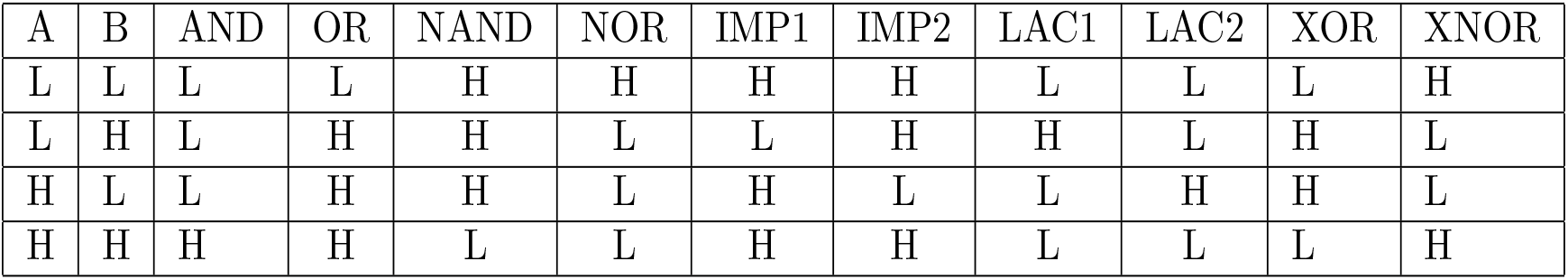
Truth Table.

| A | B | AND | OR | NAND | NOR | IMP1 | IMP2 | LAC1 | LAC2 | XOR | XNOR |
| --- | --- | --- | --- | --- | --- | --- | --- | --- | --- | --- | --- |
| L | L | L | L | H | H | H | H | L | L | L | H |
| L | H | L | H | H | L | L | H | H | L | H | L |
| H | L | L | H | H | L | H | L | L | H | H | L |
| H | H | H | H | L | L | H | H | L | L | L | H |

**Table 2:**
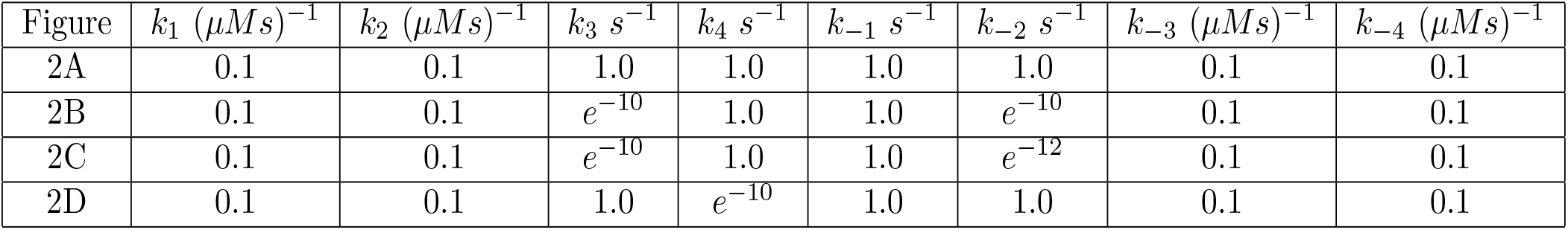
Parameters for Fig.2.

| Figure | $k_1 (\mu Ms)^{-1}$ | $k_2 (\mu Ms)^{-1}$ | $k_3 s^{-1}$ | $k_4 s^{-1}$ | $k_{-1} s^{-1}$ | $k_{-2} s^{-1}$ | $k_{-3} (\mu Ms)^{-1}$ | $k_{-4} (\mu Ms)^{-1}$ |
| --- | --- | --- | --- | --- | --- | --- | --- | --- |
| 2A | 0.1 | 0.1 | 1.0 | 1.0 | 1.0 | 1.0 | 0.1 | 0.1 |
| 2B | 0.1 | 0.1 | $e^{-10}$ | 1.0 | 1.0 | $e^{-10}$ | 0.1 | 0.1 |
| 2C | 0.1 | 0.1 | $e^{-10}$ | 1.0 | 1.0 | $e^{-12}$ | 0.1 | 0.1 |
| 2D | 0.1 | 0.1 | 1.0 | $e^{-10}$ | 1.0 | 1.0 | 0.1 | 0.1 |

A key feature that stands out in non-equilibrium is the emergence of non-monotonic behavior of promoter state occupancies as a function of input TF concentration. For example, in Fig.2 D, *P*_1_ behaves non-monotonically as a function of [*A*] for a finite range of [*A*] and [*B*]. At first, an increase in [*A*] increases the occupancy of state 1 while a further increase in [*A*] decreases the occupancy of state 1. In addition to *P*_1_, all the other promoter states can show non-monotonic behavior as well, as shown in (SI 3.1). Non-monotonicity in the occupancies of the different promoter states arises owing to their quadratic dependency on input TF concentrations (see S1 2).

Next, we ask if both the TFs can simultaneously exhibit non-monotonicity in promoter state occupancies. Interestingly, we find that at a given instance, non-monotonicity in promoter state occupancy profiles arises only for one of the two TFs. When one TF shows a non-monotonic behavior, only a monotonic profile can emerge for the other TF (see Fig.3); for formal mathematical proof, see the SI. This ties back to the direction of the non-equilibrium steady state flux which is captured by the sign of Δ*µ*. For instance, we find that for *P*_1_ or *P*_3_ to be non-monotonic with [*A*], we require Δ*µ >* 0, i.e., the cycle has to operate anticlockwise. For *P*_1_ or *P*_3_ to be non-monotonic with [*B*], we require Δ*µ <* 0 which indicates that the cycle is biased clockwise. As these two conditions cannot be met simultaneously, we observe non-monotonicity for only one of the TFs. We see a similar contradiction in the case of *P*_2_ and *P*_4_. For *P*_2_ to be non-monotonic with respect to [*A*], we need Δ*µ <* 0 while for non-monotonicity with respect to [*B*], we need Δ*µ >* 0. *P*_4_ may be non-monotonic with [*A*] only if Δ*µ >* 0 while non-monotonicity with [*B*] may be seen only when Δ*µ <* 0. The necessary conditions for non-monotonicity are discussed in greater detail in SI 2.

**Figure 3.**
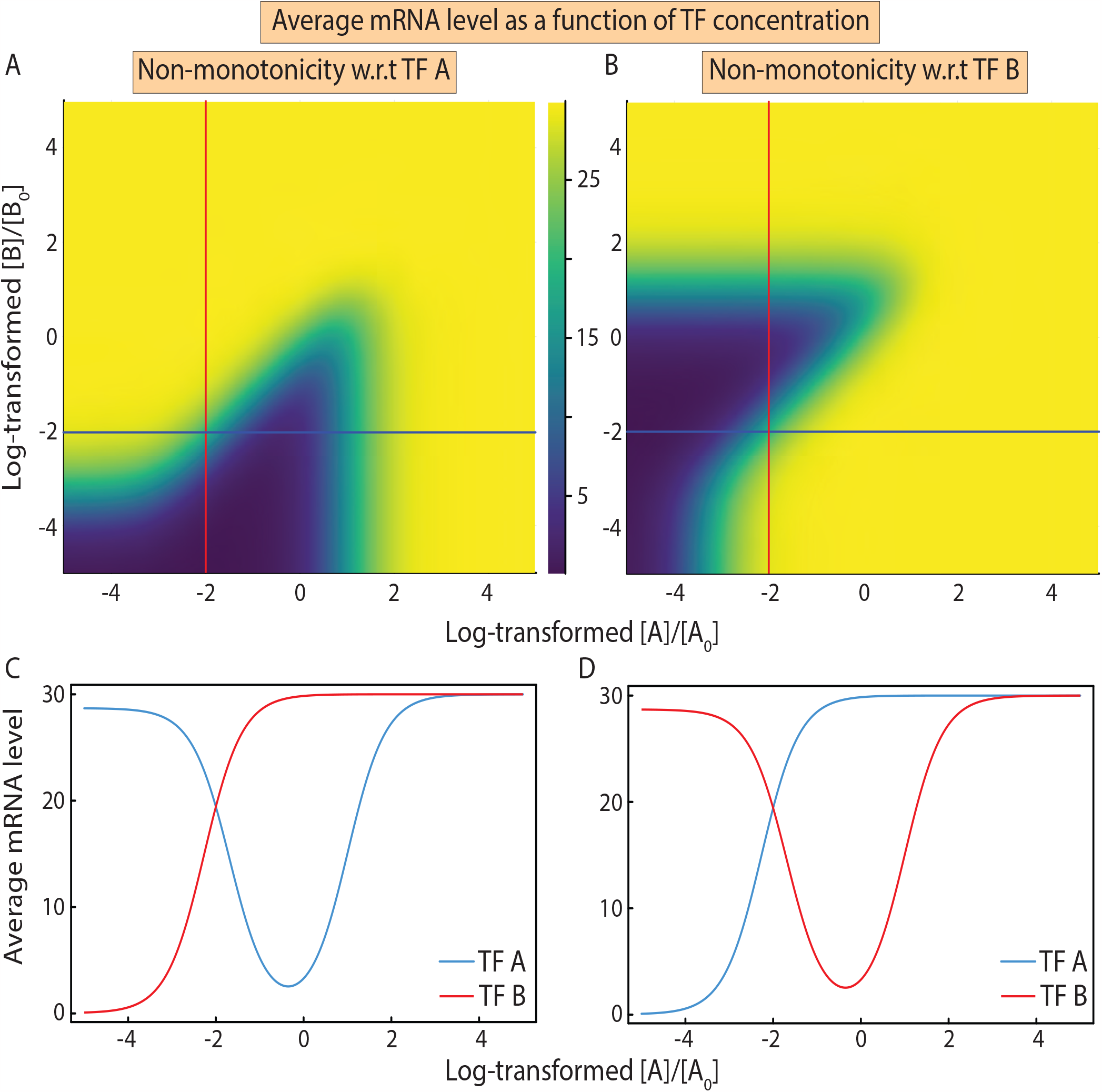
Non-monotonic promoter state occupancies alter regulatory roles of one of two TFs. A) and B) Average mRNA output as a function of log_10_ ([*A*]*/*[*A*]_0_) (x-axis) and log_10_ ([*B*]*/*[*B*]_0_)(y-axis), shown as a heatmap when A and B are activators. Colorbar ranges from 0 to 30 copies. C) and D) The average mRNA level along the blue and red lines in A) and B) respectively. Red corresponds to the line [*A*] = 0.01*µM*, Blue corresponds to the line [*B*] = 0.01*µM*. Non-monotonicity in *P*_1_ leads to an alteration of regulatory function. Non-monotonicity with respect to one of the TFs enforces monotonicity for the other. For the parameter values used, see Materials and Methods Table 3.

A systematic exploration of promoter occupancy profiles allows us to characterize average mRNA levels as a function of activator and repressor concentrations.

**Table 3:** Parameters for Fig.3.

| Figure | $k_1 (\mu Ms)^{-1}$ | $k_2 (\mu Ms)^{-1}$ | $k_3 s^{-1}$ | $k_4 s^{-1}$ | $k_{-1} s^{-1}$ | $k_{-2} s^{-1}$ | $k_{-3} (\mu Ms)^{-1}$ | $k_{-4} (\mu Ms)^{-1}$ |
| --- | --- | --- | --- | --- | --- | --- | --- | --- |
| A,C | 0.1 | 0.1 | 1.0 | $e^{-10}$ | 1.0 | 1.0 | 0.1 | 0.1 |
| B,D | 0.1 | 0.1 | 1.0 | 1.0 | $e^{-10}$ | 1.0 | 0.1 | 0.1 |
| $r_1 = 0, r_2 = r_3 = r_4 = 0.3s^{-1}, \gamma = 0.01s^{-1}$ [27] | | | | | | | | |

**Table 4:** Parameters for Fig.4.

| Gate | $k_1 (\mu Ms)^{-1}$ | $k_2 (\mu Ms)^{-1}$ | $k_3 s^{-1}$ | $k_4 s^{-1}$ | $k_{-1} s^{-1}$ | $k_{-2} s^{-1}$ | $k_{-3} (\mu Ms)^{-1}$ | $k_{-4} (\mu Ms)^{-1}$ |
| --- | --- | --- | --- | --- | --- | --- | --- | --- |
| AND | 0.1 | 0.1 | $e^{-12}$ | 1.0 | 1.0 | $e^{-14}$ | 0.1 | 0.1 |
| OR | 0.1 | 0.1 | $e^{-7}$ | $e^{-9}$ | $e^{-7}$ | $e^{-7}$ | 0.1 | 0.1 |
| NAND | 0.1 | 0.1 | $e^{-16}$ | 1.0 | 1.0 | $e^{-14}$ | 0.1 | 0.1 |
| NOR | 0.1 | 0.1 | $e^{-9}$ | $e^{-12}$ | $e^{-9}$ | $e^{-9}$ | 0.1 | 0.1 |
| For AND, OR: $r_1 = 0, r_2 = r_3 = r_4 = 0.3s^{-1}, \gamma = 0.01s^{-1}$ | | | | | | | | |
| For NAND, NOR: $r_2 = r_3 = r_4 = 0, r_1 = 0.3s^{-1}, \gamma = 0.01s^{-1}$ | | | | | | | | |

**Table 5:**
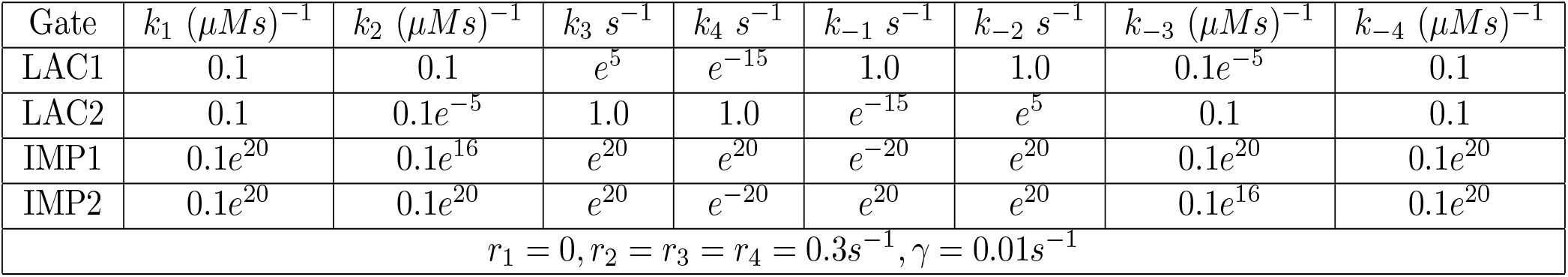
Parameters for Fig.6.

| Gate | $k_1 (\mu Ms)^{-1}$ | $k_2 (\mu Ms)^{-1}$ | $k_3 s^{-1}$ | $k_4 s^{-1}$ | $k_{-1} s^{-1}$ | $k_{-2} s^{-1}$ | $k_{-3} (\mu Ms)^{-1}$ | $k_{-4} (\mu Ms)^{-1}$ |
| --- | --- | --- | --- | --- | --- | --- | --- | --- |
| LAC1 | 0.1 | 0.1 | $e^5$ | $e^{-15}$ | 1.0 | 1.0 | $0.1e^{-5}$ | 0.1 |
| LAC2 | 0.1 | $0.1e^{-5}$ | 1.0 | 1.0 | $e^{-15}$ | $e^5$ | 0.1 | 0.1 |
| IMP1 | $0.1e^{20}$ | $0.1e^{16}$ | $e^{20}$ | $e^{20}$ | $e^{-20}$ | $e^{20}$ | $0.1e^{20}$ | $0.1e^{20}$ |
| IMP2 | $0.1e^{20}$ | $0.1e^{20}$ | $e^{20}$ | $e^{-20}$ | $e^{20}$ | $e^{20}$ | $0.1e^{16}$ | $0.1e^{20}$ |
| $r_1 = 0, r_2 = r_3 = r_4 = 0.3s^{-1}, \gamma = 0.01s^{-1}$ | | | | | | | | |

#### 2.3.1 Alteration of the regulatory function of two TFs out of equilibrium is mutually exclusive

The promoter state occupancies depend solely on the concentration of the TFs and the promoter switching rates. Whether these TFs act as repressors or activators is determined by the rate of transcription (*r*_*i*_) from the different promoter states. We find that the non-monotonicity of promoter state occupancies leads to one of the TFs switching its regulatory behavior for a range of [*A*] and [*B*]. For instance, as shown in Fig.3, when TF A and TF B are activators with the appropriate *r*_*i*_ as detailed in section 2.1, we find that at first an increase in [*A*] increases the occupancy of state 1, a state of no transcription. A further increase in [*A*] decreases the occupancy, allowing the system to switch to a transcriptionally active state. This allows activator A to effectively behave as a repressor at low concentrations. Similarly, non-monotonic promoter occupancy profiles can alter the regulatory function of repressors, allowing them to behave as activators for a given concentration range. It must be noted that switching of regulatory roles is independent of our choice of *r*_*i*_ (SI 3.2). However, the only constraint is that TF A and B cannot switch regulatory functions simultaneously since promoter occupancies exhibit non-monotonicity with one of the two TFs only.

Notably, Mahdavi et al. [26] showed theoretically that gene expression, out of equilibrium, can vary non-monotonically with TF concentration for a gene that is regulated by a single activator. In contrast, for a gene that is regulated by two TFs, the regulatory function of only one of the TFs can be altered. This mutual exclusivity of the non-monotonic behavior of gene expression of TFs imposes important constraints on the logic operations accessible in non-equilibrium regimes, as we discuss in the ensuing sections.

### 2.4 AND, NAND, NOR, and OR logic operations can be attained in and out of equilibrium

First, we study AND, OR, NAND, and NOR gates as these operations are more common and have also been synthetically implemented in living systems [11, 37–41]. To compare the design principles of building AND, OR, NAND, and NOR gates in and out of equilibrium, we first summarize the principles governing equilibrium logic. Since these gates have been studied in great detail using an equilibrium framework, they serve as a reference to study transcriptional logic out of equilibrium. A visual examination of the promoter occupancy profiles seen in Fig.2 reveals that AND and OR gates can be built when TF A and B act as activators whereas NAND and NOR logic can be achieved when A and B act as repressors.

To explore how these gates can built optimally, we sweep the parameter space of equilibrium circuits spanned by *K*_*d,A*_, *K*_*d,B*_ and *ϵ*_*c*_ and select circuits with an *l*_1_ norm below 0.1 (see SI 4). We find that AND, NOR, and NAND gates require weak TF-promoter binding and strong cooperativity between the TFs. This finding is consistent with previous studies that have demonstrated that cooperativity is essential for the design of AND and NAND gates [7, 11], due to the sharp, switch-like behavior it grants to the mRNA level as we move from the ([*A*]_*High*_, [*B*]_*Low*_) and ([*A*]_*Low*_, [*B*]_*High*_) inputs to the ([*A*]_*High*_, [*B*]_*High*_) input. In contrast, OR gates occupy two distinct regimes in the parameter space; one of the regimes is similar to AND, NAND, and NOR gates, while the other regime prefers *K*_*d*_ *<* 1, and *ϵ*_*c*_ *<* 0. In the second regime, TFs strongly bind the promoter but together they bind weakly. Such binding of TFs could be a result of repulsion or steric hindrance [42].

Next, we seek to explore the principles governing transcriptional logic out of equilibrium. To this end, we sweep the eight-dimensional parameter space spanned by promoter switching rates. In particular, we select a set of default rates 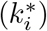 detailed in Table 7, corresponding to an equilibrium circuit with *K*_*d,A*_ = *K*_*d,B*_ and *ϵ*_*c*_ = 0, and drive the system out of equilibrium by allowing any combination of rates to be increased or decreased from their default values. We select all optimum gates with an *l*_1_ norm below 0.1 and use this set for further analysis. A key finding is that AND, NAND, NOR, and OR gates can be built out of equilibrium as well. This result is not surprising since promoter occupancy profiles out of equilibrium can mimic the corresponding equilibrium profiles. However, building these gates out of equilibrium involves energy expenditure. To assess the energetic costs, we plot the distributions of Δ*µ* (see Fig.4 A, B, C, D). The distributions peak near Δ*µ* = 0, indicating that a small dissipation of energy is sufficient to create these gates. Moreover, these distributions are symmetric about Δ*µ* = 0, as that optimum AND, NAND, OR, and NOR gates do not distinguish between the TFs A and B, i.e, they do not distinguish between the ([*A*]_*Low*_, [*B*]_*High*_) and ([*A*]_*High*_, [*B*]_*Low*_) inputs. We can exchange the binding/unbinding rates of TF A with those of TF B and obtain an optimum gate with the net steady-state flux operating in the opposite direction.

**Table 6:** Parameters for sweeping equilibrium spaces.

| Parameter | Symbol | Values |
| --- | --- | --- |
| Rate of transcription | $r$ | $0.3 \text{ s}^{-1}$ |
| Rate of degradation of mRNA | $\gamma$ | $0.01 \text{ s}^{-1}$ |
| Low input concentration | $[A]_{Low}, [B]_{Low}$ | $0.01 \text{ }\mu\text{M}$ |
| High input concentration | $[A]_{High}, [B]_{High}$ | $1.0 \text{ }\mu\text{M}$ |
| $\langle m \rangle$ cutoff | - | 5 |
| $l_1$ norm cutoff | - | 0.1 |
| For AND, OR: $r_1 = 0, r_2 = r_3 = r_4 = r$ | | |
| For NAND, NOR: $r_2 = r_3 = r_4 = 0, r_1 = r$ | | |

**Table 7:** Parameter values for sweeping the non-equilibrium space.

| Parameter | Symbol | Values |
| --- | --- | --- |
| Rate of transcription | $r$ | $0.3 \text{ s}^{-1}$ |
| Rate of degradation of mRNA | $\gamma$ | $0.01 \text{ s}^{-1}$ |
| Low input concentration | $[A]_{Low}, [B]_{Low}$ | $0.01 \text{ }\mu\text{M}$ |
| High input concentration | $[A]_{High}, [B]_{High}$ | $1.0 \text{ }\mu\text{M}$ |
| Default binding rates | $k_1^*, k_2^*, k_{-3}^*, k_{-4}^*$ | $0.1 (\text{ }\mu\text{Ms})^{-1}$ |
| Default unbinding rates | $k_{-1}^*, k_{-2}^*, k_3^*, k_4^*$ | $1.0 \text{ s}^{-1}$ |
| $\langle m \rangle$ cutoff | - | 5 |
| $l_1$ norm cutoff | - | 0.1 |
| For LAC1, LAC2, IMP1, IMP2, AND, OR: $r_1 = 0, r_2 = r_3 = r_4 = r$ | | |
| For NAND, NOR: $r_2 = r_3 = r_4 = 0, r_1 = r$ | | |

**Figure 4.**
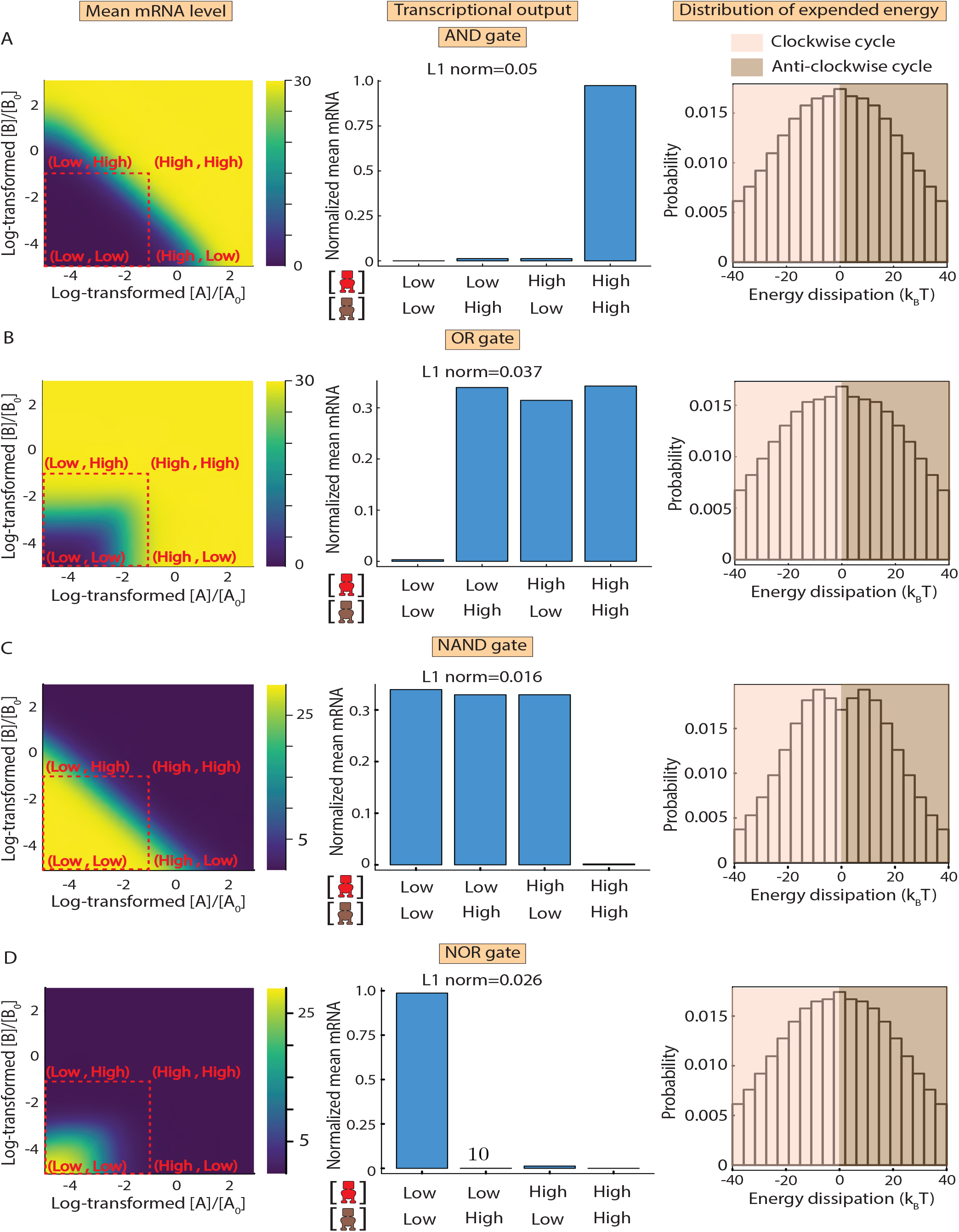
AND, OR, NAND and NOR out of equilibrium: Examples of non-equilibrium AND, OR, NAND and NOR gates, parameter values can be found in Materials and Methods Table 4. The first column is a representation of the mRNA level as a function of log_10_ ([*A*]*/*[*A*]_0_) (x-axis) and log_10_ ([*B*]*/*[*B*]_0_)(y-axis), with the points marked in red acting as the input concentrations. In the second column, we show the normalized transcriptional output of the circuit for each input concentration, along with its *l*_1_ norm. In the third column, we show the distribution of the dissipated energy (Δ*µ*) for all optimum gates, lighter shade indicates the cycle is biased clockwise, darker shade indicates the cycle is biased anticlockwise A) AND, B) OR, C) NAND, D) NOR.

To further unravel the underlying design principles of the best-performing gates, we choose a lower cutoff on the *l*_1_ norm (*<* 0.1) such that we have sufficient data for analyses and look at the relationship between the corresponding promoter switching rates (Table 7). In particular, we look at the fold change 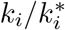 of the binding and unbinding rates at each of the four edges of the cycle.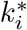 is the default rate for the transition under consideration. These relations provide crucial insights into the propensity at each of the four edges and reveal if binding or unbinding is favored overall. As there is symmetry in the clockwise and anticlockwise cycles, we only consider those operating in the anticlockwise direction. In the case of optimum AND, NOR, and NAND gates, there is only one parameter regime (Fig.5 A, SI 5.1). The fold change in the unbinding rate is greater than that of the binding rate when the TFs interact with bare DNA. This relation is reversed and binding is favored when one of the TFs is already bound to DNA in an effect reminiscent of cooperativity (see Fig.5 B). These relations do not change for optimum gates with the cycle biased clockwise. For OR gates, while one regime is identical to the other gates under consideration (Fig.5 D), in the second parameter regime, binding is preferred to unbinding when the TFs interact with bare DNA. Binding is unevenly deterred for B when A is already bound (see Fig.5 E) in an effect reminiscent of repulsion. For clockwise cycles, we find that binding is unevenly deterred for A when B is already bound.

**Figure 5.**
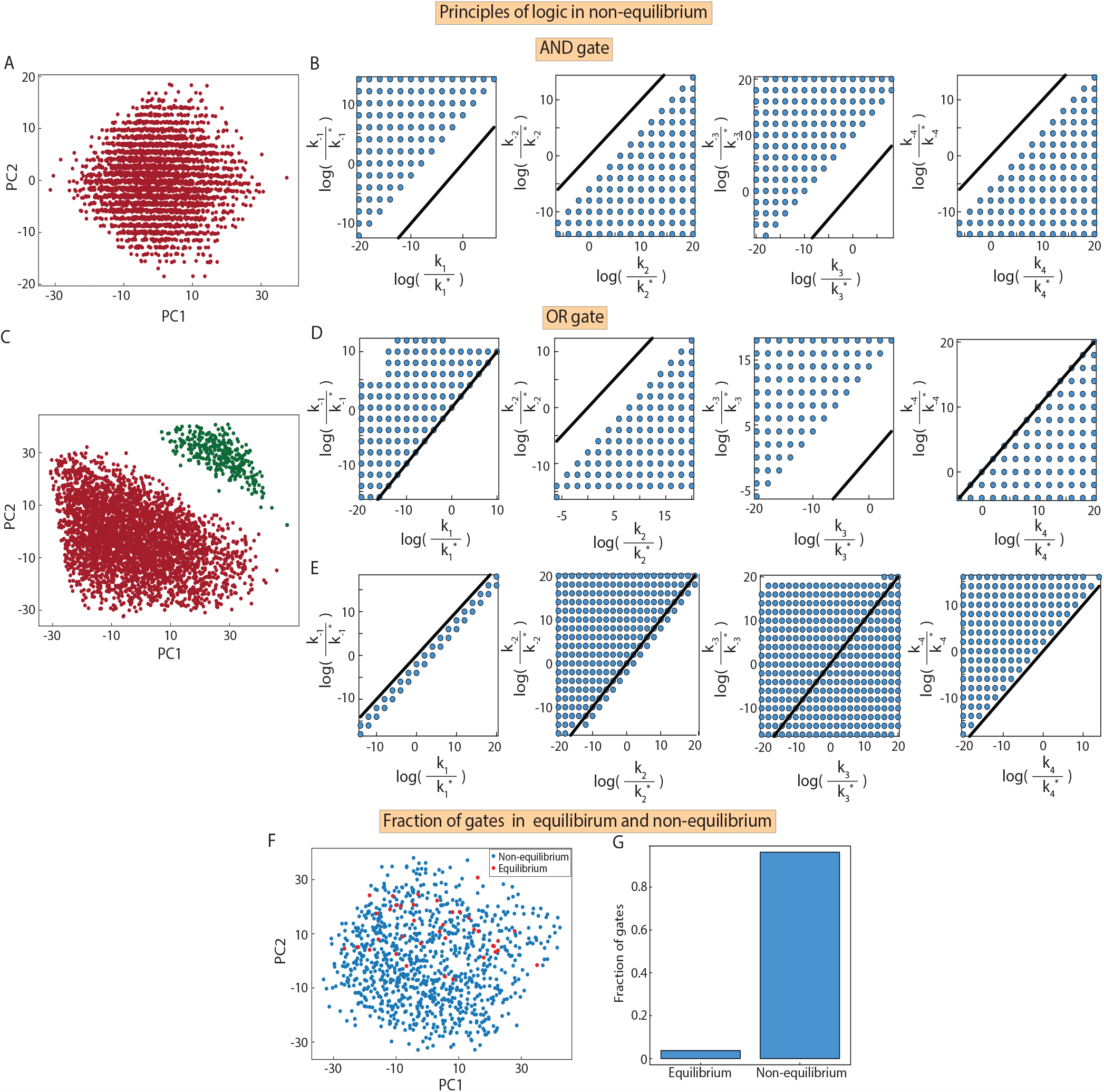
Design principles of optimum AND and OR gates out of equilibrium. A) PCA of optimum non-equilibrium AND gates with Δ*µ >* 0. The eight-dimensional space spanned by the fold change in the promoter switching rates, 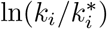, is projected onto a 2-dimensional space, spanned by the first two principal components. B) Associated rate relations between binding and unbinding rates at each edge for non-equilibrium AND gates, the black line serves as a reference with a unit slope and no intercept. Along this line, the unbinding and binding rates at each edge change by the same magnitude. C) PCA of optimum non-equilibrium OR rates with Δ*µ >* 0. D) Associated rate relations between the binding and unbinding rates at each edge for C) (green points) are shown. E) Associated rate relations between the binding and unbinding rates at each edge for C) (red points) are shown. F) A visual depiction of optimum equilibrium and non-equilibrium AND gates encountered in the eight-dimensional space on a PCA plot. G) A bar graph depicting the fraction of optimum AND gates encountered in and out of equilibrium. The ratio of their heights is 4:96.

Having explored the design principles of AND, NAND, NOR, and OR gates in both in and out of equilibrium, we are in a position to carry out a systematic comparison between these regimes. In the case of the AND, NAND, and NOR operations, we observe similar principles across both regimes, that binding to TF-bound DNA is favored over binding to bare DNA. In the case of OR gates, two possible parameter regimes exist both in and out of equilibrium; one of these two is similar to the regime found for AND, NAND, and NOR gates. Thus, despite having a higher number of free parameters out of equilibrium, we do not find any new parameter regimes for these gates. One key difference between the two regimes, however, lies in the number of optimum gates that can be attained; the number of optimal gates in non-equilibrium far exceeds the number in equilibrium, as shown in Fig.5 G, SI 5.2). By uniformly sampling a collection of optimum gates emanating in and out of equilibrium, we find that equilibrium gates are scattered in a pool of non-equilibrium gates (Fig.5 F,G).

### 2.5 The emergence of novel logic operations out of equilibrium

Next, we ask if novel logic operations emerge in transcription when TF-promoter binding happens out of equilibrium. Our analysis showed that promoter occupancies out of equilibrium can exhibit non-monotonicity as a function of TF concentration (Fig.2 D), thereby allowing the TFs to take on different regulatory roles in different concentration intervals (Fig.3). This switching of regulatory function gives rise to four new logic operations out of equilibrium; the two implication logics, A implies B (IMP1) and B implies A (IMP2), as well as their negations which we refer to as LAC1 and LAC2 respectively (Table 1), due to its similarity to the logic operation created by CRP and LacR at the lac operon [7, 43]. The emergence of these operations holds for different schemes of transcription and is not restricted to our choice of transcription rate (*r*_*i*_) from different promoter states (SI 8).

We sweep the non-equilibrium parameter space to find a collection of optimum gates and look at the distribution of Δ*µ* (Fig.6). Unlike in the case of AND, OR, NOR, and NAND gates, the distributions for these four gates are peaked at non-zero values. A large number of gates, however, are still well within a reasonable energy budget of 40*k*_*B*_*T*, the free energy released by the hydrolysis of two molecules of ATP [30]. We also note that these distributions are not symmetric around Δ*µ* = 0. This is because, for all these logic operations, the optimum gates do distinguish between A and B. The ([*A*]_*High*_, [*B*]_*Low*_) input elicits a different response than the ([*A*]_*Low*_, [*B*]_*High*_) input; the binding and unbinding rates associated with TF A cannot be exchanged with the binding and unbinding rates associated with TF B, i.e., the probability flux has to be biased in one particular direction.

**Figure 6.**
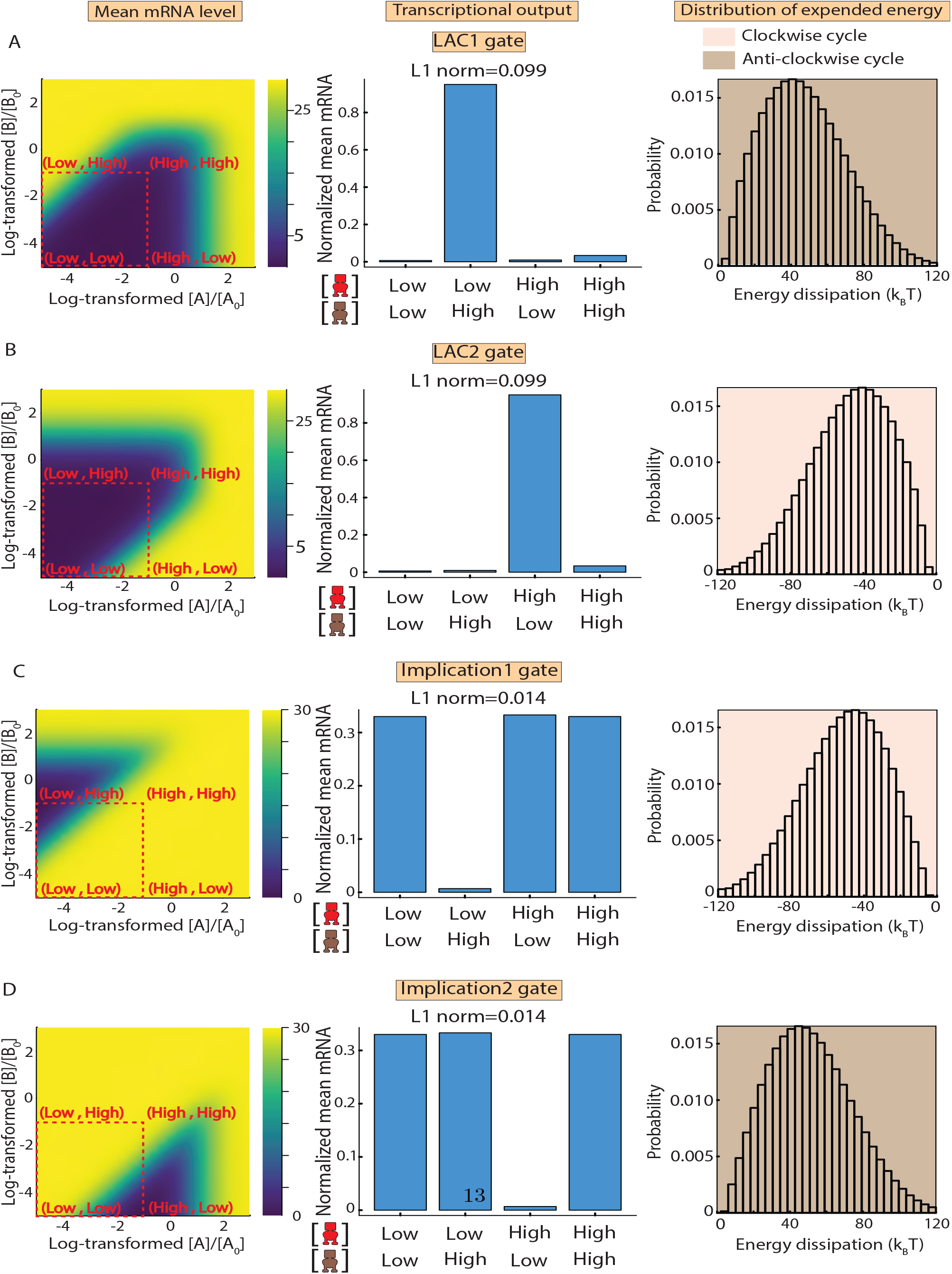
LAC1, LAC2, IMP1, IMP2 are found exclusively out of equilibrium: A) LAC1, B) LAC2, C) IMP1, D) IMP2. Examples of gates found exclusively out of equilibrium, parameter values can be found in Materials and Methods Table 5. The first column is a representation of the average mRNA level as a function of log_10_([*A*]*/*[*A*]_0_) (x-axis) and log_10_([*B*]*/*[*B*]_0_)(y-axis), shown on a heatmap, with the points marked in red depicting the input concentrations. In the second column, we show the normalized transcriptional output of the circuit for each input concentration, along with its *l*_1_ norm. In the third column, we show the distribution of the dissipated energy for all optimum gates. A lighter shade indicates that the circuits are biased clockwise, darker shade indicates the circuits are biased anticlockwise.

To make IMP and LAC gates in equilibrium, we would require one of the TFs to be a repressor and the other an activator, as there is no switching of regulatory roles in equilibrium. For instance, creating LAC1 or IMP2 gates entails that A act as a repressor and B act as an activator. Hence, the non-monotonicity in promoter state occupancies, resulting from non-equilibrium binding kinetics, expands the regulatory potential of transcription factors without affecting their molecular nature.

Taken together, our results show that, by moving out of equilibrium, various logic gates can be generated using two transcription factors that have the same molecular function, i.e., activator or repressor. Each of these logic gates is characterized by a defined set of promoter switching rates. We next sought to systematically analyze the promoter switching rates that underlie the various non-equilibrium gates. To this end, we projected all the hitherto explored optimum logic gates (i.e. AND, NAND, OR, NOR, LAC1, LAC2, IMP1, IMP2) that span the space of promoter switching rates on a 2D-UMAP plot (see Fig.7). Non-equilibrium logic gates that can also arise in equilibrium (AND, NAND, OR, NOR) form a well-separated cluster that does not display directional bias in

**Figure 7.**
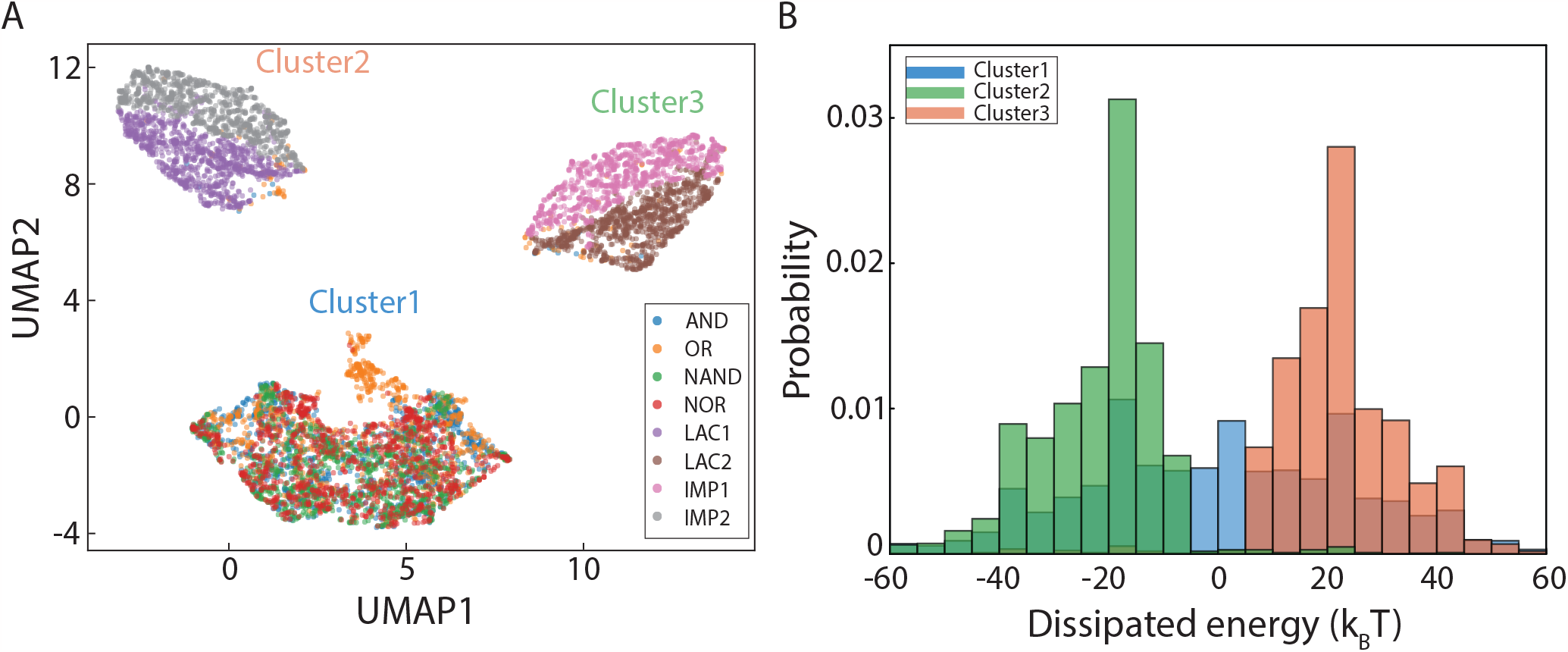
UMAP projection of the optimal gates. A) Optimum AND, NAND, OR, NOR, LAC1, LAC2, IMP1, and IMP2 gates in the space spanned by the promoter switching rates are visualized on a UMAP. Cluster 1 contains AND, OR, NAND, and NOR. Cluster 2 contains IMP2, LAC1, AND and OR. Cluster 3 contains IMP1, LAC2, AND and OR. B) The histograms of Δ*µ* for each of the three clusters are shown.

Δ*µ*. The logic gates that are unique to non-equilibrium (LAC1, LAC2, IMP1, IMP2) emerge as two distinct clusters, which display opposite directional bias in Δ*µ*. Notably, there is some spillover from AND and OR gates into these clusters. These results suggest that the non-equilibrium logic gates operate in three distinct non-overlapping parameter regimes, with two of these regimes giving rise to logic operations that solely arise out of equilibrium.

## 3 Discussion

Cells carry out complex logic computations at the level of transcription by integrating different combinations and concentrations of transcription factors. While decades of work have shed light on the principles of transcriptional logic, [5, 7, 44] none of them have accounted explicitly for non-equilibrium mechanisms at the level of the promoter. In this manuscript, we attempt to understand the mechanisms underlying simple transcriptional logic by studying a gene regulated by two transcription factors.

Our study showed that AND, NAND, OR, and NOR gates can be made both in and out of equilibrium and have similar design principles in both regimes. For the design of AND and NAND in particular, non-equilibrium circuits can mimic equilibrium circuits by substituting high values of cooperativity with a small amount of energy dissipation. This may confer an advantage to non-equilibrium circuits as cooperativity is usually non-specific and weak [7, 22, 45]. Interestingly, there are far more optimum gates to be found in the non-equilibrium regime than in equilibrium. This important feature may confer robustness to gene regulatory networks. Since transcriptional decision-making is a fundamental cellular function, a stimulating plausibility is that evolutionary pressures could shape the transcriptional machinery to move out of equilibrium to maximize the number of ways cells can achieve different computational capabilities.

### 3.1 Non-monotonic genetic responses result in new logic operations but also constraints the accessible computations

Previous studies have demonstrated that non-monotonic transcriptional output curves could emerge for genes regulated by one transcription factor, through mechanisms such as energy expenditure [26] or feed-forward loops [46]. Non-monotonicity alters the regulatory behavior of the TF. However, in both prokaryotes and eukaryotes, genes are often regulated by multiple transcription factors. For a gene regulated by two TFs, we find that a non-monotonic response is possible for only one of the TFs. Such non-monoticity allows us to access new logic operations out of equilibrium, such as the LAC and IMP logics, but also imposes a constraint that forbids the emergence of other logic operations, such as the XOR and EQ (or XNOR) gates.

The XOR and EQ gates are difficult to create and often require complex architectures [7, 47–49]. In our model, they remain inaccessible in both equilibrium and non-equilibrium regimes due to the nature of the promoter state occupancies. To create an XOR or EQ gate, we would require both TFs to play the dual role of an activator and a repressor [7] but this is not possible due to the constraint in the non-monotonicity of the promoter state occupancies, as discussed in section 2.3.1.

While the prospect of finding experimental signatures of transcriptional logic out of equilibrium is an exciting one, our findings can also potentially aid the process of building synthetic transcriptional circuits out of equilibrium. In particular, the larger repertoire of possibilities that can be attained out of equilibrium should pave the way for building novel synthetic circuits.

## Acknowledgments

S.C. is supported by the Ramalingaswami Re-entry Fellowship (BT/HRD/35/02/2006), a reentry scheme of the Department of Biotechnology, Ministry of Science and Technology, Government of India. T.C.M. is supported by the Lumina quaeruntur grant (LQ200522301) of the Czech Academy of Sciences.

## 4 Materials and Methods

### 4.1 Parameter sweeps

#### 4.1.1 Equilibrium Spaces

Data used for section 2.4 was generated by sweeping the parameter space spanned by log_10_(*K*_*d,A*_*/*[*A*]_0_), log_10_(*K*_*d,B*_*/*[*B*]_0_) and *ϵ*_*c*_ such that −10 ≤ log_10_(*K*_*d,A*_*/*[*A*]_0_), log_10_(*K*_*d,B*_*/*[*B*]_0_) ≤ 10 with increments of 0.1 and −50≤ *ϵ*_*c*_(*k*_*B*_*T*) ≤ 50 with increments of 0.5. Other parameters and cutoffs are documented in 6.

#### 4.1.2 Non-equilibrium spaces

Data used for sections 2.5 and 2.4 were generated by sweeping the parameter space spanned by 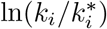 such that 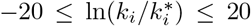 with increments of 2 for each promoter switching rate *k*_*i*_. All parameters and cutoffs are documented in Table 7. Note that this method also selects equilibrium gates. All non-equilibrium gates were selected and the histograms for Δ*µ* were generated. From these optimum gates, the top 10^6^ gates with the lowest *l*_1_ norms were selected for the analysis described in Fig.5. The corresponding *l*_1_ norm cutoffs may be found in Table. 8. For our analysis and the symmetry between the rates, only gates with Δ*µ >* 0 were considered. For Fig.5 A, B, randomly chosen points in the eight-dimensional space spanned by 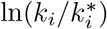 were projected onto a PCA for better visualization. Cluster identities were assigned using Agglomerative clustering. All analyses were performed using Python’s Sci-kit learn library.

**Table 8:** *l*_1_ norm cutoffs *<* 0.1.

| Gate | AND | OR | NOR | NAND |
| --- | --- | --- | --- | --- |
| $l_1$ norm cutoff | 0.028 | 0.065 | 0.028 | 0.07 |

### 4.2 UMAP

8000 optimum gates were randomly selected from the pool of optimum AND, OR, NAND, NOR, LAC1, LAC2, IMP1, and IMP2 gates (1000 each), which were obtained by sweeping the parameter space as described in section 4.1.2. The eight-dimensional space spanned by 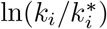 were projected on a 2D-plane using a UMAP (n neighbors = 20) [50]. Cluster identities were assigned using Agglomerative Clustering from Python’s Sci-kit learn library.

## Supplementary Information

### 1 General solution for the steady-state promoter state occupancies

We compute the steady-state occupancies for each promoter state using the Matrix Tree Theorem (MTT) [1]. MTT provides a diagrammatic method to compute steady-state occupancies of occupying the various vertices on a graph. Briefly, MTT states that in steady-state, the probability of occupying any state (a vertex on a graph) is proportional to the sum of products of rate constants over all spanning trees rooted in that state. A spanning tree rooted at a vertex *i*, is a directed subset of edges on the graph where every vertex is visited only once, and there are no outgoing edges from the root.

The weights of the various promoter states in the steady state as a function of the eight promoter switching rates and the concentrations of TF A and TF B are given by

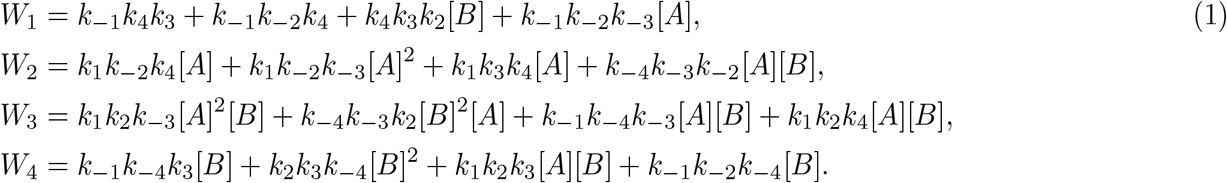

And we get the corresponding promoter state occupancies by normalizing the weights as

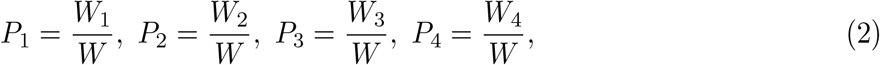

where *P*_*i*_ is the steady-state occupancy of the *i*-th promoter state, and *W* =∑_*i*_ *W*_*i*_. In thermodynamic equilibrium, the Kolmogorov criterion is given by

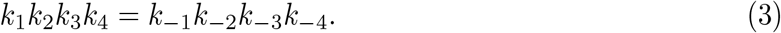

The Kolmogorov criterion states that for a cyclic graph to be reversible, the product of the rates of all clockwise edges is equal to the product of the rates on all the anticlockwise edges [1]. To draw a comparison to the equilibrium description of the system, we divide the weights of the various promoter states in Eqn. 1 by *W*_1_,

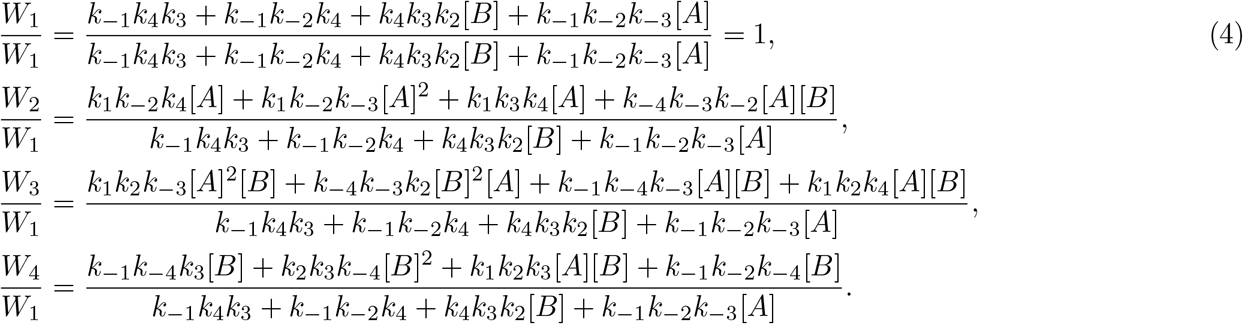

By factoring out some of the rates, we get

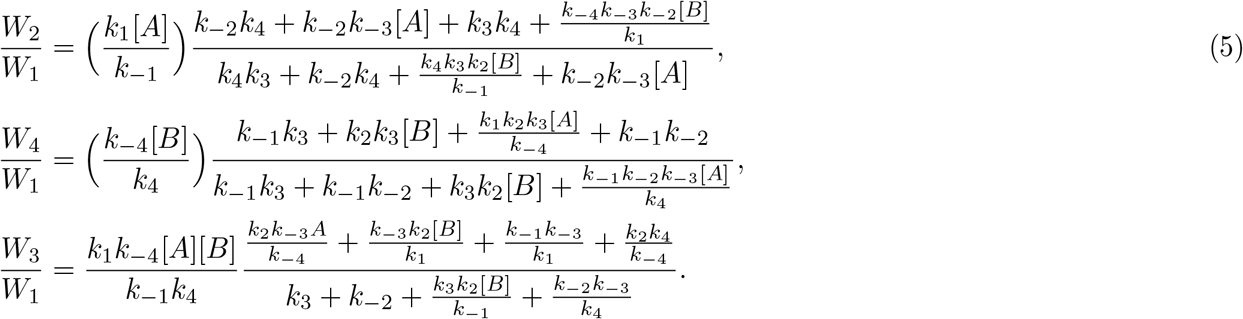

On substituting the cooperativity 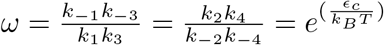 in the numerator of *W*_3_, we get

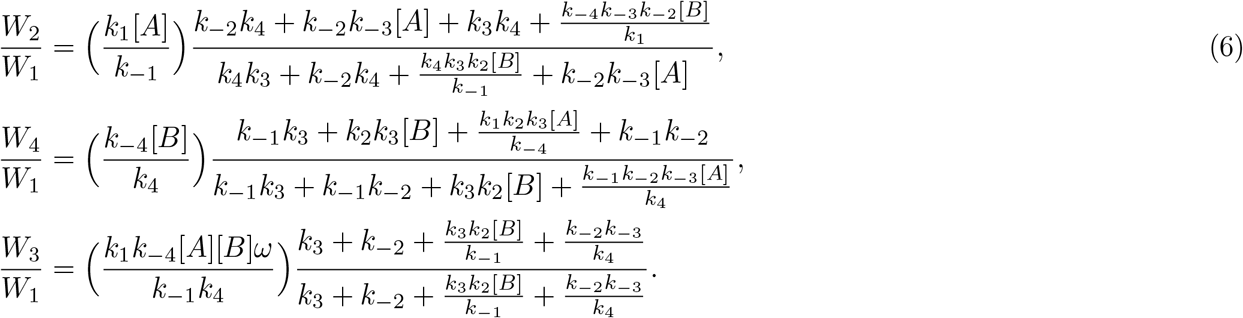

Using the Kolmogorov criterion, we obtain the weights of different promoter states in thermodynamic equilibrium,

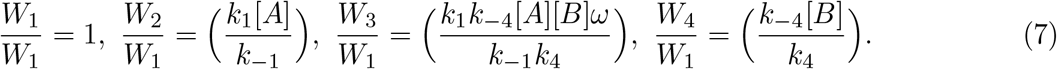

Substituting 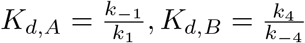, we get

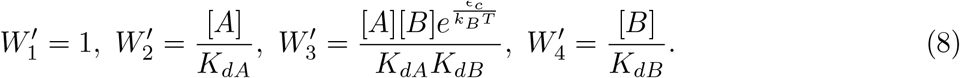

This is exactly what we would get using a grand canonical description of the system in thermodynamic equilibrium. The corresponding partition function is [2]

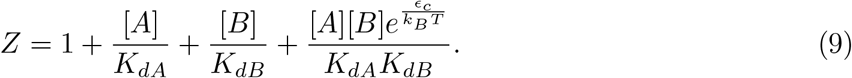

Note that none of the equilibrium *P*_*i*_ can show non-monotonic behavior with any of the TFs. If we write down the *P*_*i*_ in equilibrium as a function of one of the TFs while keeping the concentration of the other constant, they can have one of two forms,

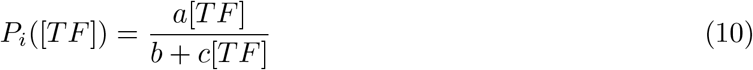

Or

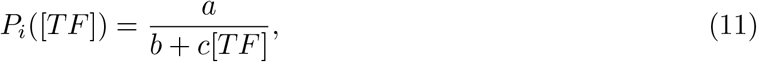

where *a, b, c >* 0.

The first form monotonically increases with TF concentration while the second form monotonically decreases with TF concentration. Thus we find that *P*_1_ monotonically decreases with [*A*] and [*B*], *P*_3_ monotonically increases with [*A*] and [*B*], *P*_2_ monotonically increases with [*A*] and decreases with [*B*] while *P*_4_ monotonically decreases with [*A*] and increases with [*B*].

### 2 Promoter state occupancy can exhibit non-monotonicity with respect to one of the TFs

#### 2.1 Promoter occupancy of state 1

For each promoter occupancy *P*_*i*_, non-monotonicity is possible with respect to either [*A*] or [*B*] but not both. To prove this, we find necessary conditions for non-monotonicity for each *P*_*i*_ with respect to [*A*] and [*B*]. We first consider the case of *P*_1_. On taking the partial derivative with[*A*], we get a form

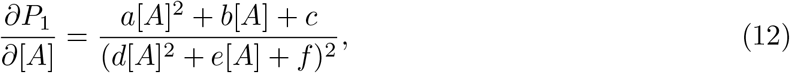

where *a, b, c, d, e, f* are functions of the promoter switching rates and [*B*]. For non-monotonicity, this derivative must change sign at some point [*A*] *>* 0. The denominator is always positive, so we focus our attention on the numerator which is a quadratic in [*A*]. We find the sufficient and necessary conditions for a sign change by looking for a positive root. From our calculations shown in the accompanying *Mathematica* notebook, ‘*Proofs for non-monotonicity of the promoter state occupancies as a function of TF concentration*,’ we observe that *a, b <* 0. The parabola defined by *a*[*A*]^2^ + *b*[*A*] + *c* opens downward and has a maximum at 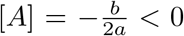. We have two options for *c*. Either *c >* 0 or *c*≤0.

If *c <* 0, then the parabola either has no real roots or two roots at some negative value of [*A*]. For non-monotonicity, we need *c >* 0 as the derivative has to change the sign.

This is a sufficient and necessary condition that guarantees non-monotonicity of *P*_1_ with[*A*] for some values of [*B*], but the conditions are difficult to interpret so we look at the necessary condition of Δ = *b*^2^ − 4*ac >* 0 given *c >* 0. We find the necessary conditions for non-monotonicity, given by

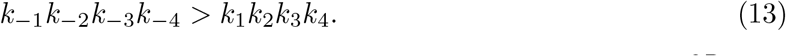

That is, *P*_1_ can be non monotonic with [*A*] only if Δ*µ >* 0. We do a similar exploration for 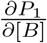 and using the same argument, find the necessary conditions for non-monotonicity with respect to [*B*]. We find that in his case we need

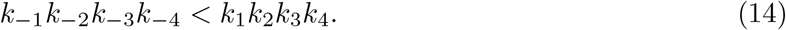

That is, Δ*µ <* 0. As these two conditions necessary for the non-monotonicity of *P*_1_ with respect to A and B are contradictory, we can conclude that *P*_1_ can behave non-monotonically with respect to either [*A*] or [*B*] but not both.

#### 2.2 Promoter occupancy of state 3

As in the case of *P*_1_, we look for the conditions for a sign change in the partial derivatives of *P*_3_ with [*A*] and [*B*] and find the same contradiction. We take the case of the partial derivative with [*A*], which has a similar form as above,

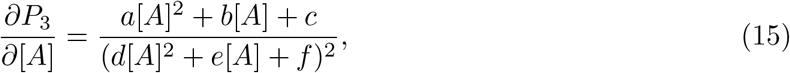

and consider the numerator, as the denominator is always positive. Here, we find that *b, c >* 0, leaving two options for *a*. If *a >* 0, the parabola defined by *a*[*A*]^2^ + *b*[*A*] + *c* curves upwards and if it has real roots, then both of them must be at some [*A*] *<* 0. If *a <* 0, then the parabola curves downwards and it is possible to find a real root at some [*A*] *>* 0.

As before, imposing this condition on *a* gives us sufficient and necessary conditions for the nonmonotonicity of *P*_3_ with respect to [*A*]. These sufficient and necessary conditions are difficult to interpret so we look for necessary conditions by mandating Δ = *b*^2^ − 4*ac >* 0 given *a <* 0.

We find that, as in the case of *P*_1_, a positive value of Δ*µ* is required. That is,

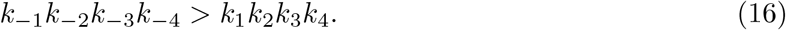

On performing the same exercise with 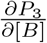, we find that the necessary condition for non-monotonicity with respect to [*B*] is given by Δ*µ <* 0. That is,

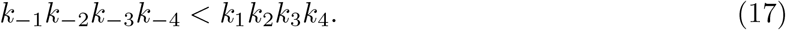

Again, *P*_3_ cannot be non-monotonic with [*A*] and [*B*] simultaneously. Moreover, these are the same necessary conditions for non-monotonicity as seen for *P*_1_.

The solutions for *P*_2_ and *P*_4_ are provided in the Mathematica notebook, as attached below.

### 3 Supplementary Figures

#### 3.1 Effect of promoter switching rates on the promoter state occupancies out of equilibrium

In the following figures, we take a set of default promoter switching rates and increase or decrease one of them at a time by a factor of *e*^±10^. We plot the promoter occupancies as a heatmap as a function of log_10_([*A*]*/*[*A*]_0_) and log_10_([*B*]*/*[*B*]_0_) where [*A*]_0_ = [*B*]_0_ = 1*µM*. The default rates are

- 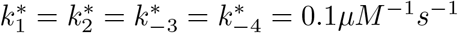
- 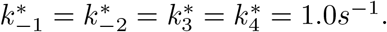

Note that these default rates correspond to an equilibrium circuit with *K*_*d,A*_ = *K*_*d,B*_, *ϵ*_*c*_ = 0. Non-monotonicity can arise for each of the four promoter state occupancies.

**Figure 1.**
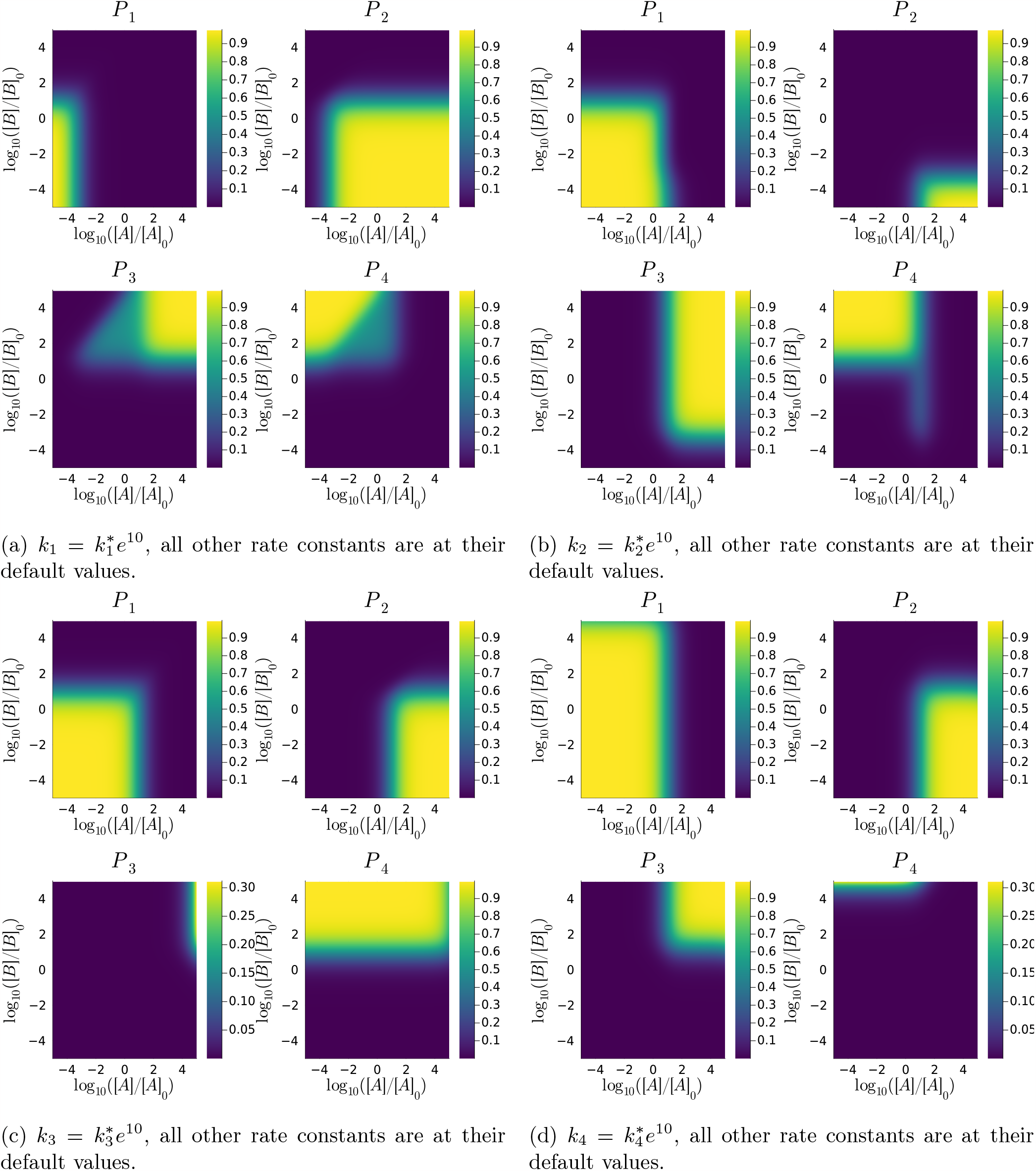
Effect of speeding up each clockwise rate: *P*_*i*_ as a function of log_10_([*A*]*/*[*A*]_0_) (x-axis) and log10([*B*]*/*[*B*]_0_) (y-axis).

**Figure 2.**
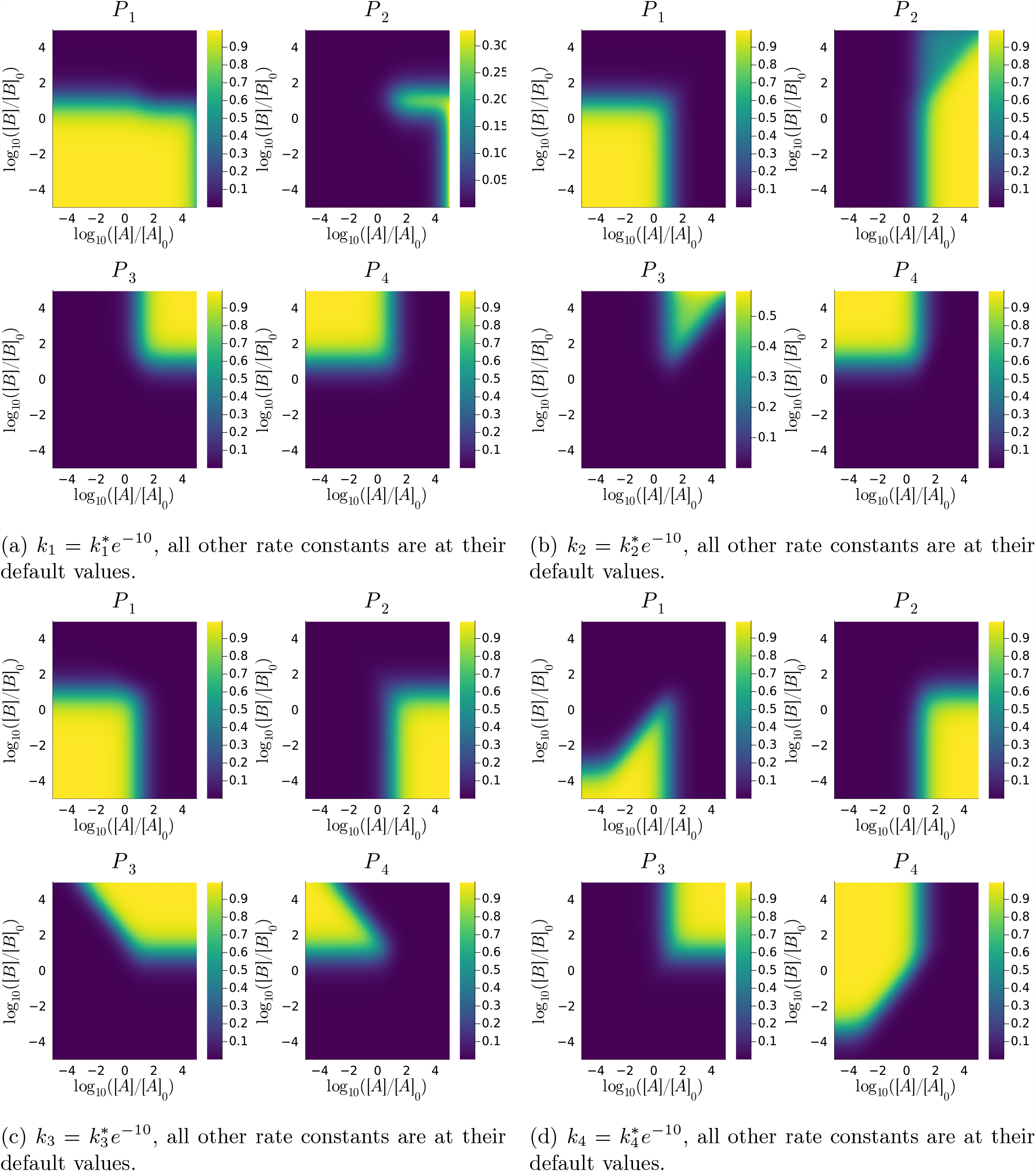
Effect of slowing down each clockwise rate: *P*_*i*_ as a function of log_10_([*A*]*/*[*A*]_0_) (x-axis) and log_10_([*B*]*/*[*B*]_0_) (y-axis).

**Figure 3.**
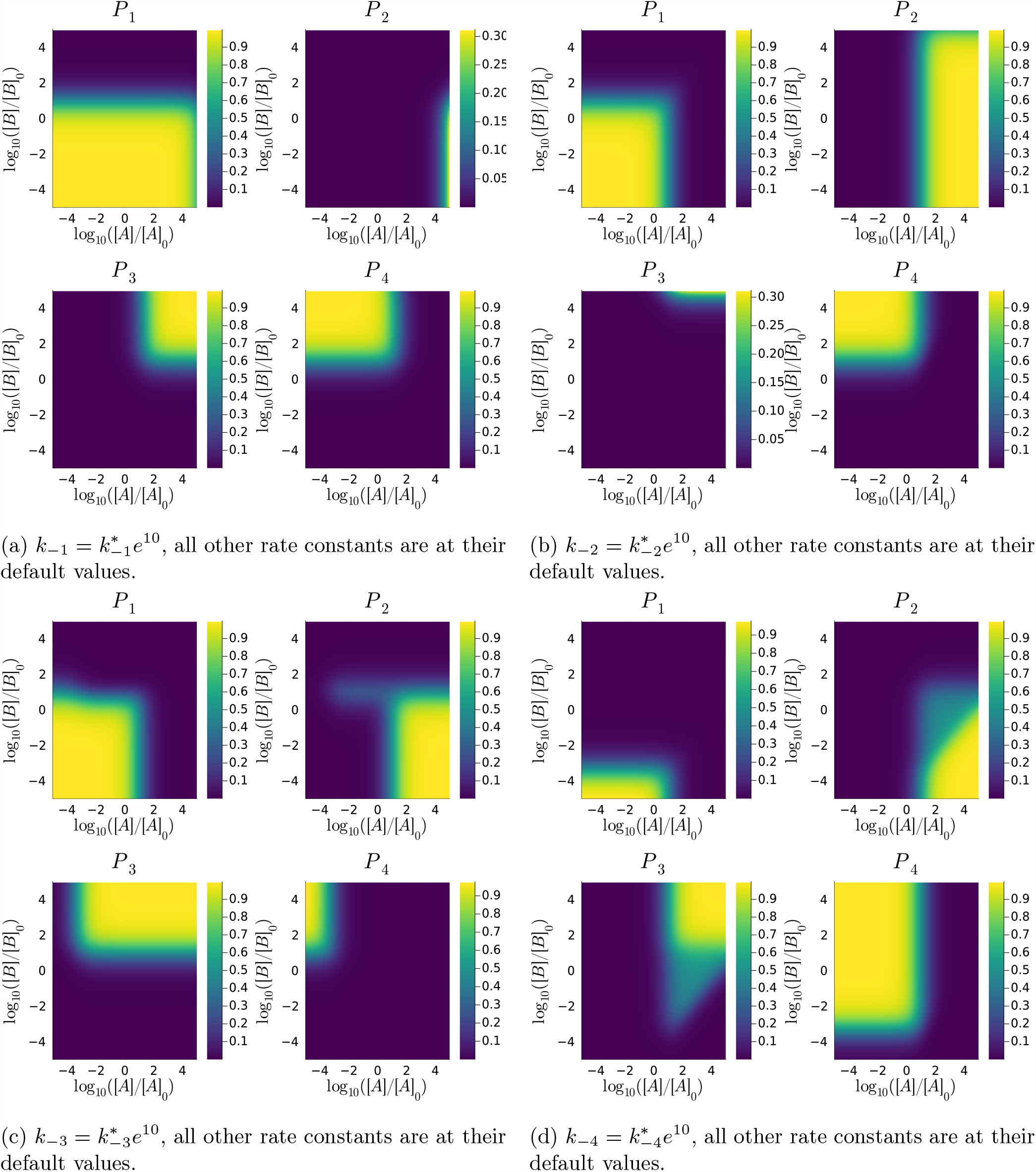
Effect of speeding up each anticlockwise rate: *P*_*i*_ as a function of log_10_([*A*]*/*[*A*]_0_) (x-axis) and log_10_([*B*]*/*[*B*]_0_) (y-axis).

**Figure 4.**
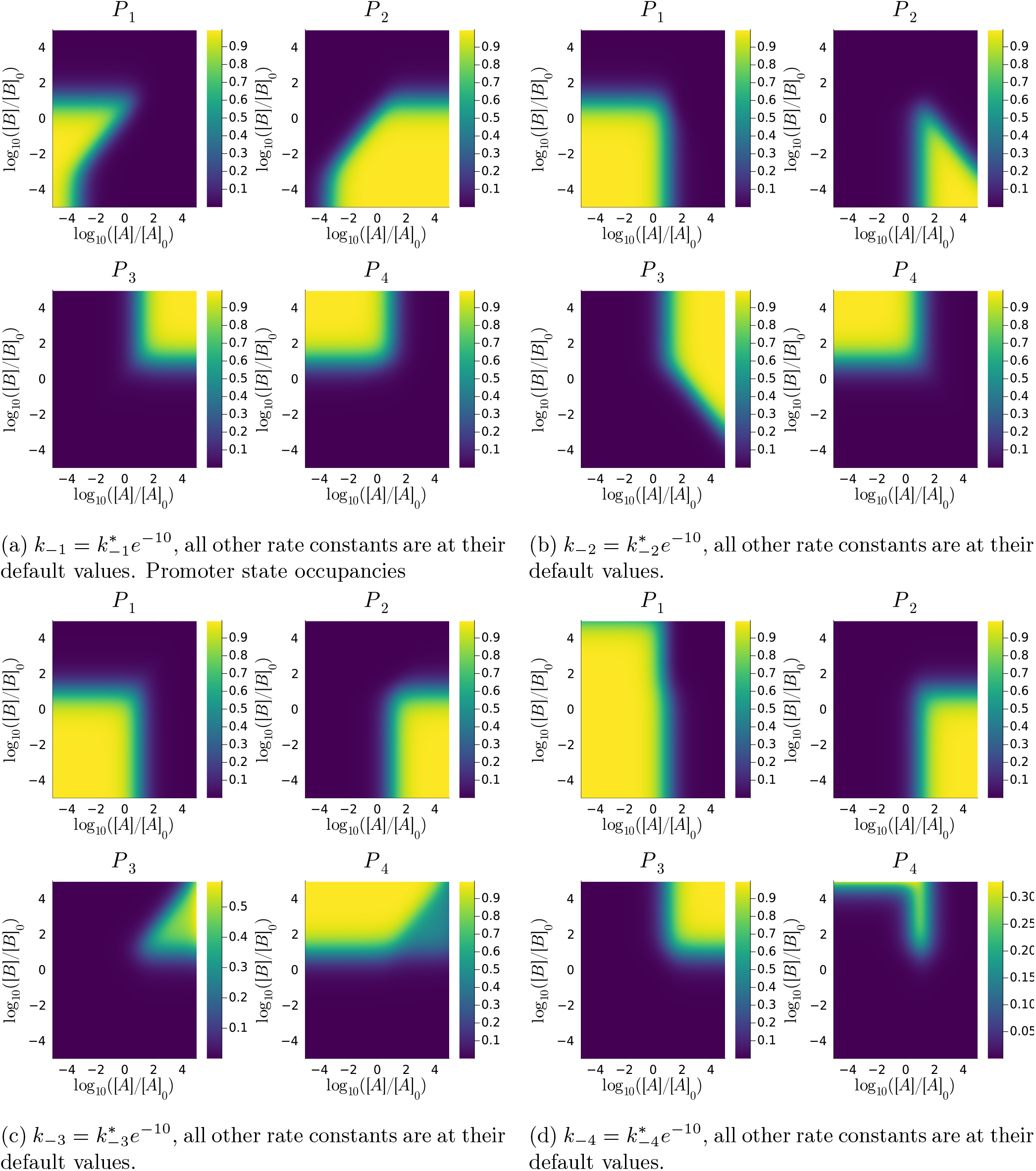
Effect of slowing down each anticlockwise rate: *P*_*i*_ as a function of log_10_([*A*]*/*[*A*]_0_) (x-axis) and log_10_([*B*]*/*[*B*]_0_) (y-axis).

#### 3.2 Alteration of regulatory behavior of TFs is possible for many gene expression strategies

Here we look at three commonly used gene expression strategies when A and B are activators [3]:

- One or more: *r*_1_ = 0.*r*_2_ = *r*_3_ = *r*_4_ = *r* (fig 3 in Main Text)
- All bound: *r*_1_ = *r*_2_ = *r*_4_ = 0, *r*_3_ = *r*
- Partial Activation: 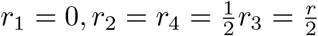

Note that with these strategies, the TFs act as repressors at higher concentrations as the alteration of regulatory roles depend on the non-monotonicity of *P*_3_.

##### 3.2.1 All bound

**Figure 5.**
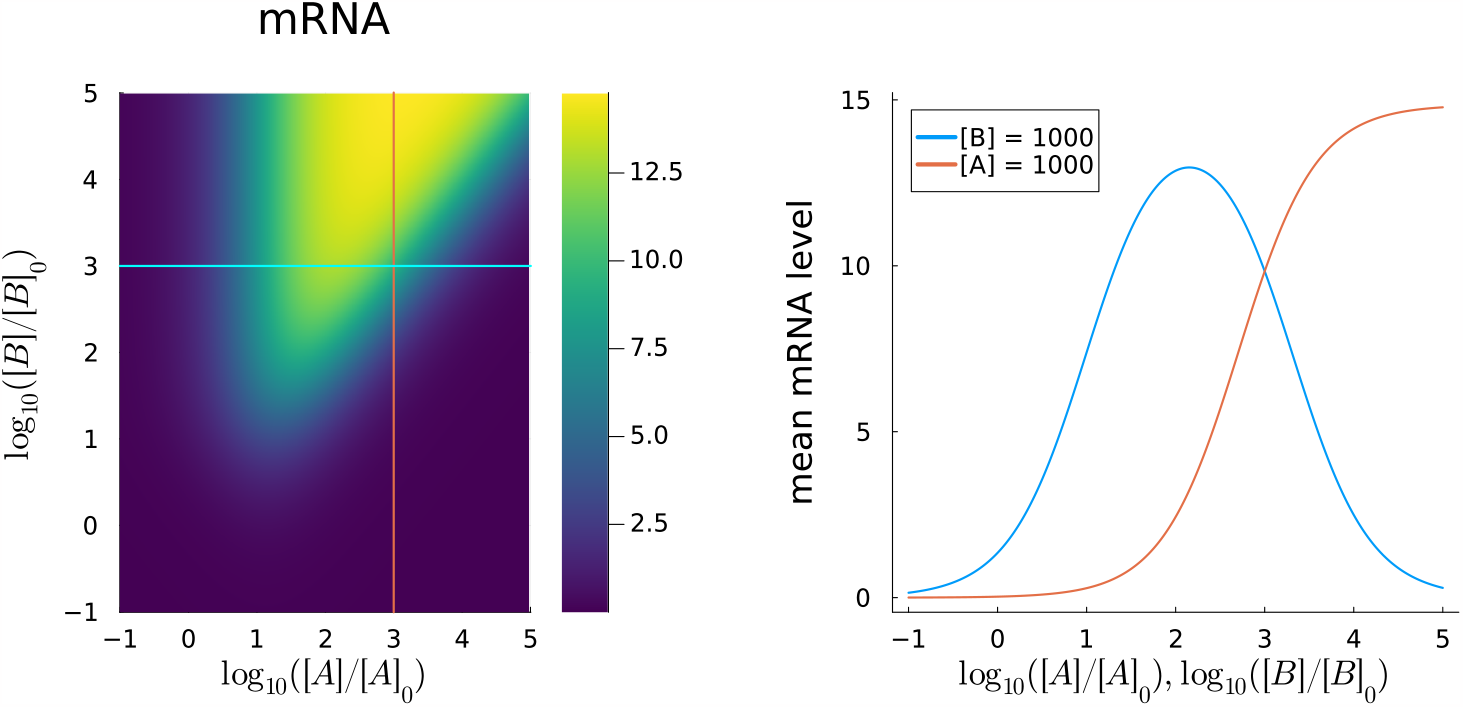
All or nothing strategy: A acts as a repressor at higher concentrations. *r* = 0.3*s*^−1^, *γ* = 0.01*s*^−1^. *k*_1_ = *k*_−3_ = *k*_−4_ = 0.1*µM* ^−1^*s*^−1^.*k*_−1_ = *k*_−2_ = *k*_3_ = *k*_4_ = 1.0*s*^−1^.*k*_2_ = 0.1*e*^−20^*µM* ^−1^*s*^−1^. Left: Heatmap of mean mRNA level as a function of log_10_([*A*]*/*[*A*]_0_) (x-axis) and log_10_([*B*]*/*[*B*]_0_) (y-axis). Blue line is at [*B*] = 10^3^*µM*, red line is at [*A*] = 10^3^*µM*. Right: Mean mRNA level along the blue and red lines.

**Figure 6.**
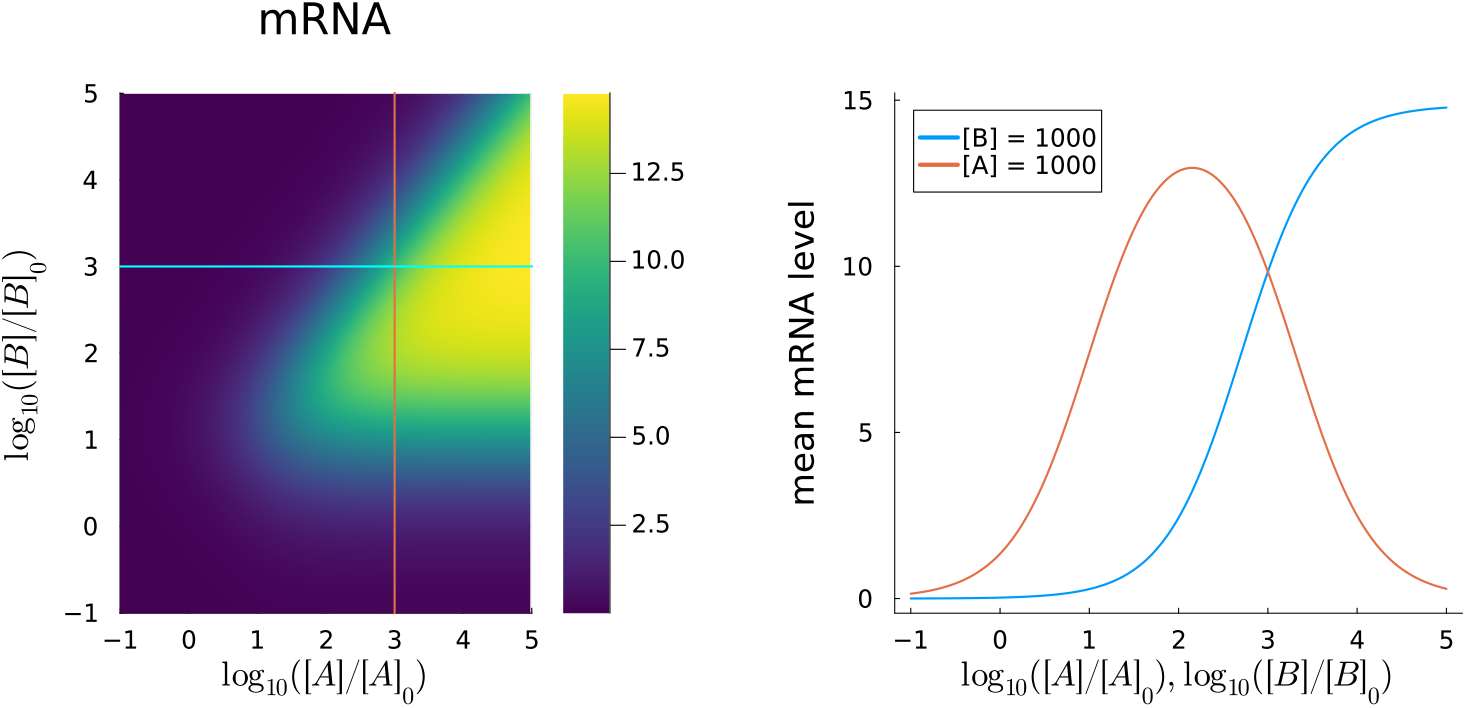
All or nothing strategy: B acts as a repressor at higher concentrations. *r* = 0.3*s*^−1^, *γ* = 0.01*s*^−1^. *k*_1_ = *k*_2_ = *k*_−4_ = 0.1*µM* ^−1^*s*^−1^.*k*_−1_ = *k*_−2_ = *k*_3_ = *k*_4_ = 1.0*s*^−1^.*k*_−3_ = 0.1*e*^−20^*µM* ^−1^*s*^−1^. Left: Heatmap of mean mRNA level as a function of log_10_([*A*]*/*[*A*]_0_) (x-axis) and log_10_([*B*]*/*[*B*]_0_) (y-axis). Blue line is at [*B*] = 10^3^*µM*, red line is at [*A*] = 10^3^*µM*. Right: Mean mRNA level along the blue and red lines.

##### 3.2.2 Partial Activation

**Figure 7.**
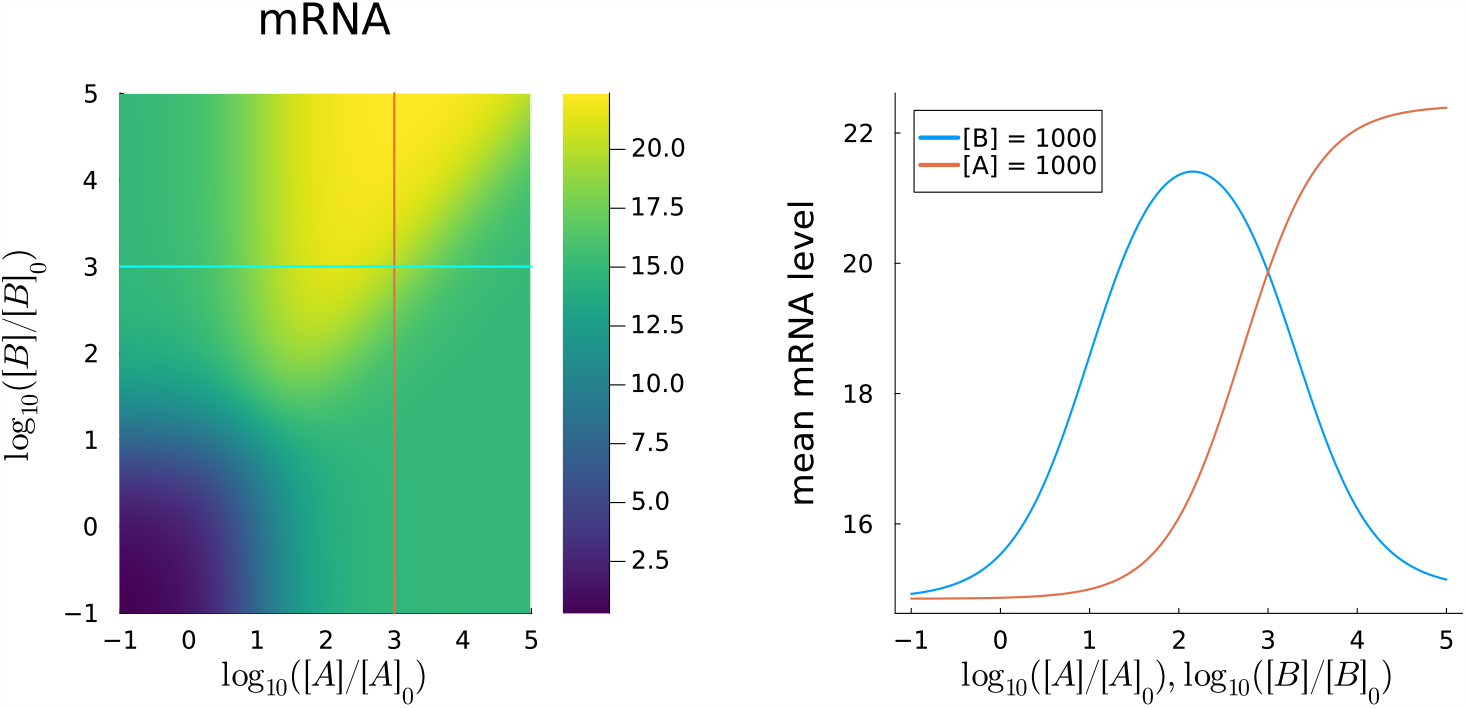
Partial activation strategy: A acts as a repressor at higher concentrations. *r*_3_ = 0.3*s*^−1^, *r*_2_ = *r*_4_ = 0.15*s*^−1^, *γ* = 0.01*s*^−1^. *k*_1_ = *k*_−3_ = *k*_−4_ = 0.1*µM* ^−1^*s*^−1^.*k*_−1_ = *k*_−2_ = *k*_3_ = *k*_4_ = 1.0*s*^−1^.*k*_2_ = 0.1*e*^−20^*µM* ^−1^*s*^−1^. Left: Heatmap of mean mRNA level as a function of log_10_([*A*]*/*[*A*]_0_) (x-axis) and log_10_([*B*]*/*[*B*]_0_) (y-axis). Blue line is at [*B*] = 10^3^*µM*, red line is at [*A*] = 10^3^*µM*. Right: Mean mRNA level along the blue and red lines.

**Figure 8.**
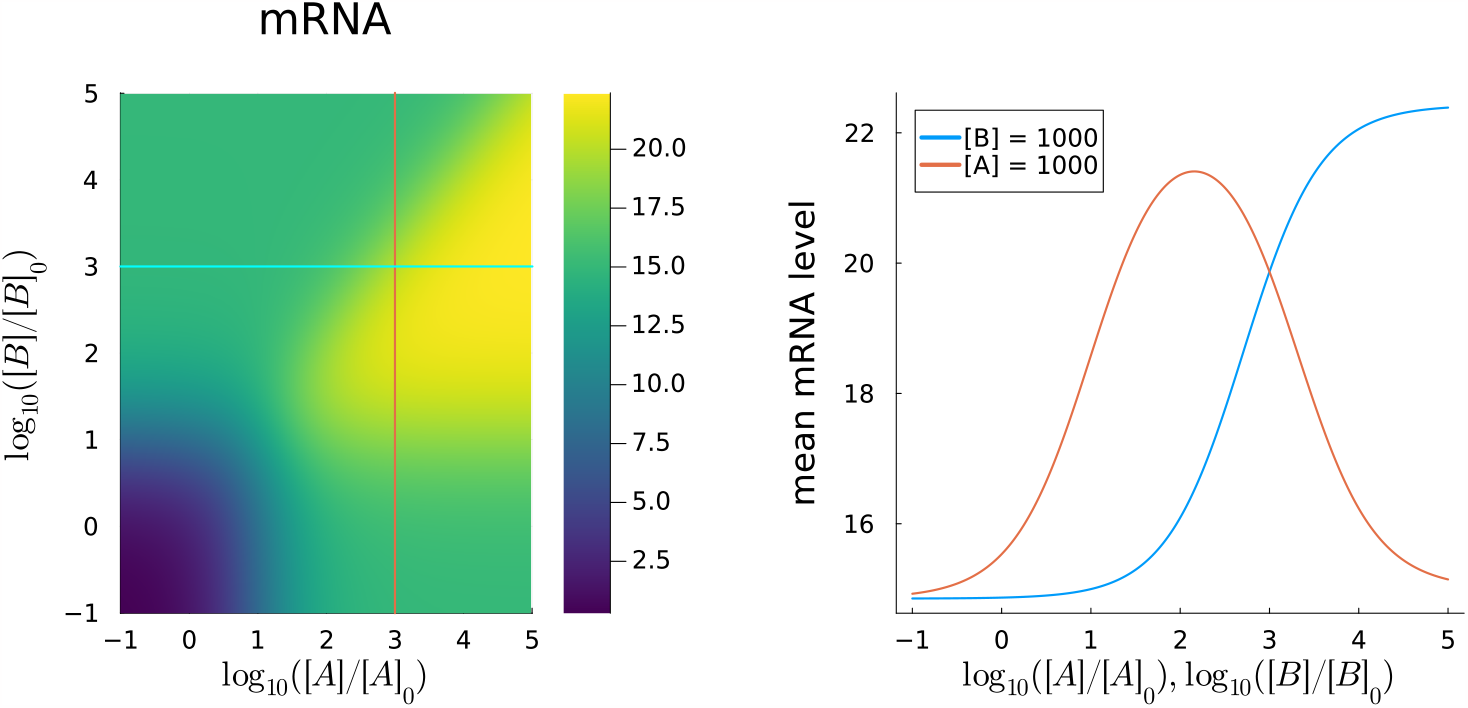
Partial activation strategy: B acts as a repressor at higher concentrations. *r*_3_ = 0.3*s*^−1^, *r*_2_ = *r*_4_ = 0.15*s*^−1^, *γ* = 0.01*s*^−1^. *k*_1_ = *k*_2_ = *k*_−4_ = 0.1*µM* ^−1^*s*^−1^.*k*_−1_ = *k*_−2_ = *k*_3_ = *k*_4_ = 1.0*s*^−1^.*k*_−3_ = 0.1*e*^−20^*µM* ^−1^*s*^−1^. Left: Heatmap of mean mRNA level as a function of log_10_([*A*]*/*[*A*]_0_) (x-axis) and log_10_([*B*]*/*[*B*]_0_) (y-axis). Blue line is at [*B*] = 10^3^*µM*, red line is at [*A*] = 10^3^*µM*. Right: Mean mRNA level along the blue and red lines.

#### 4 Design Principles of gates at Equilibrium

**Figure 9.**
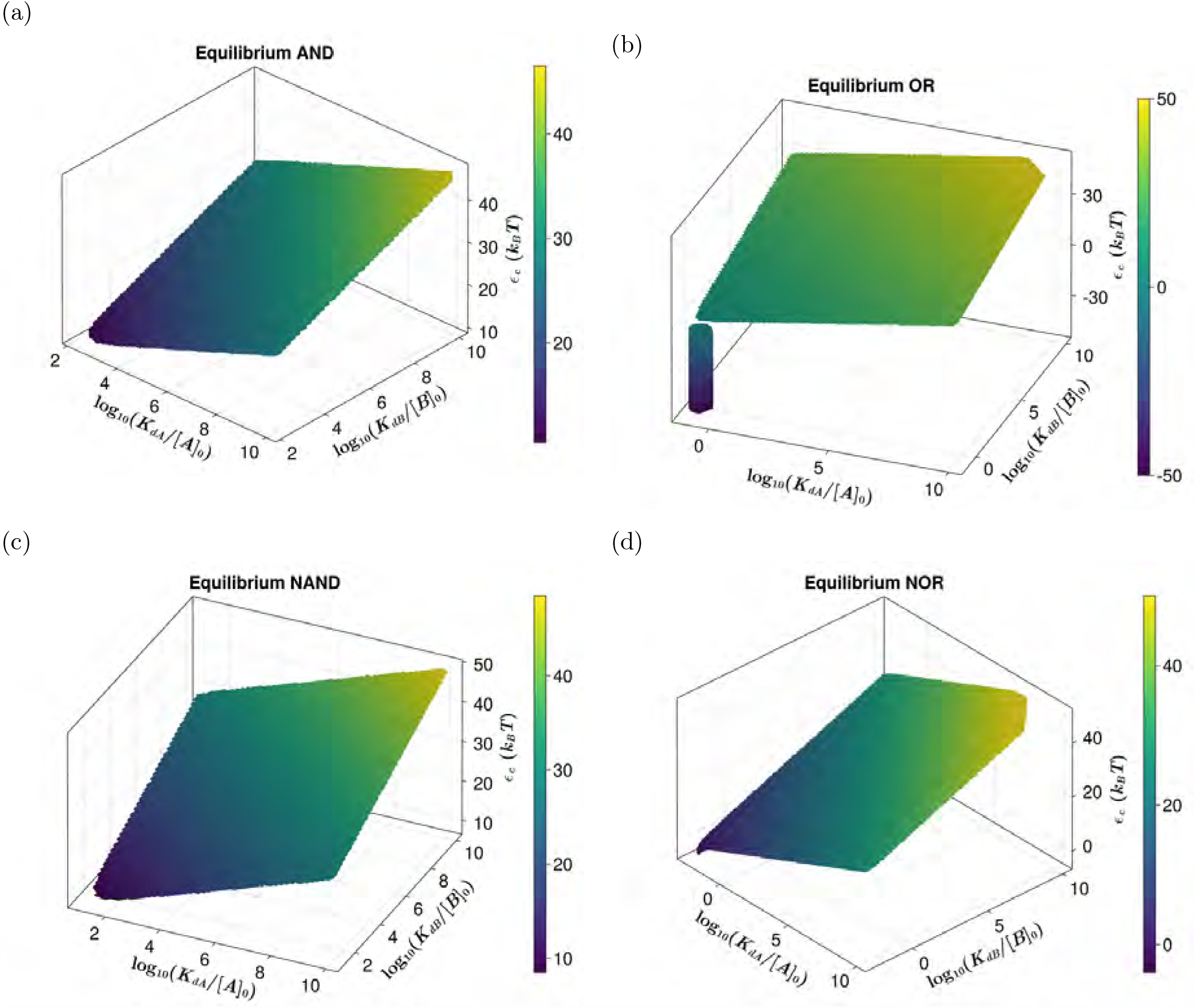
Design principles of gates in equilibrium: Parameter values for optimum equilibrium NOR gates on a 3D plot, log_10_(*K*_*dA*_*/*[*A*]_0_) (x-axis), log_10_(*K*_*dB*_*/*[*B*]_0_) (y-axis) and *ϵ*_*c*_(*k*_*B*_*T*) (z - axis). Data was generated using the parameters and methods documented in Materials and Methods 4.1.1. (a) AND, (b) OR, (c) NAND, (d) NOR.

#### 5 Design Principles of Gates out of equilibrium

##### 5.1 Complete rate Relations for AND, OR, NAND, NOR

**Figure 10.**
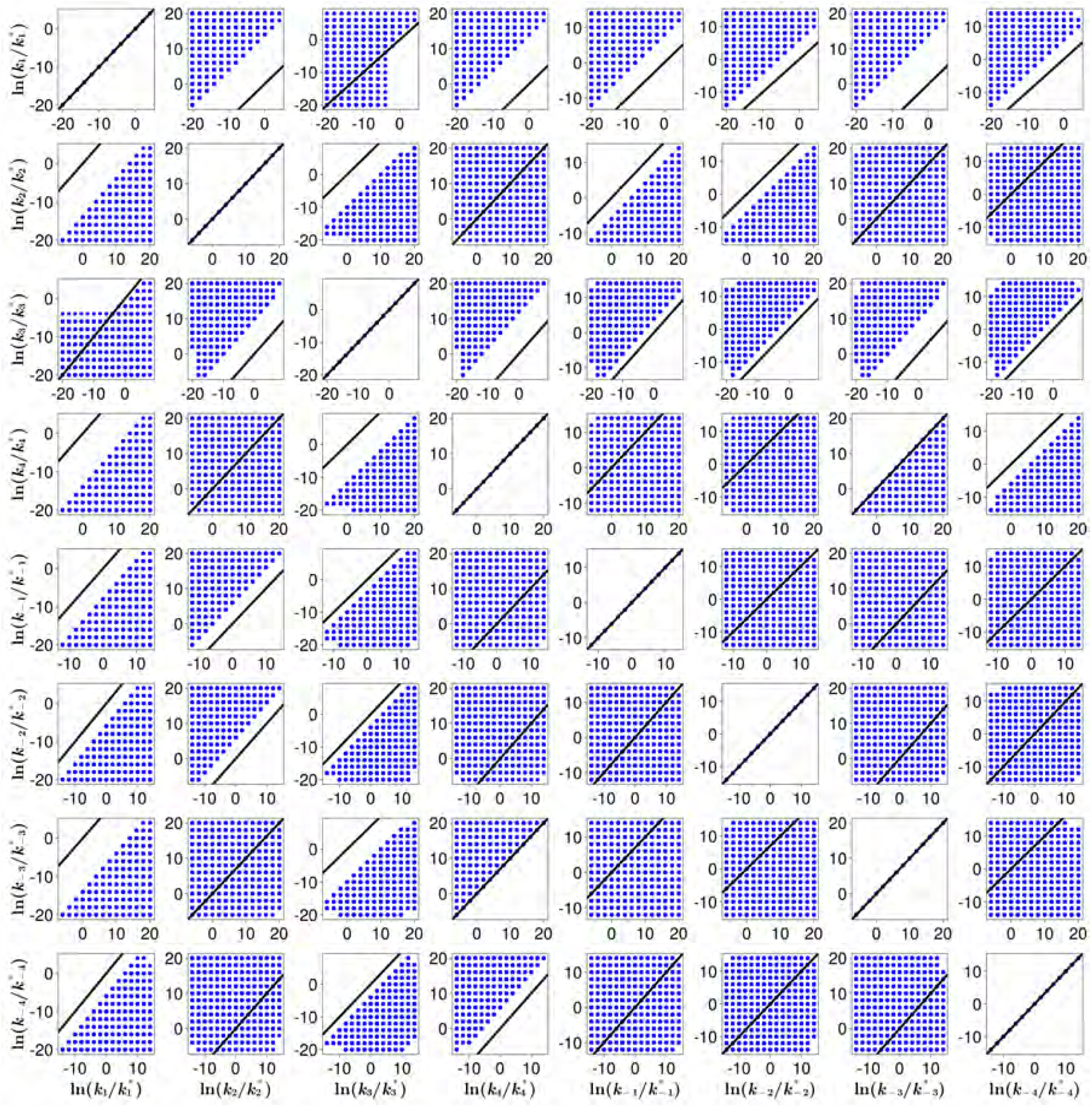
AND: Rate relations between the fold change 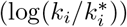 in each pair of the eight promoter switching rates for optimum gates operating anticlockwise. Column labels (bottom) indicate the rate on the x-axis. Row labels (right) indicate the rates on the y-axis. Black line has slope = 1 and intercept = 0. Data was generated using the parameters and methods documented in Materials and Methods 4.1.2.

**Figure 11.**
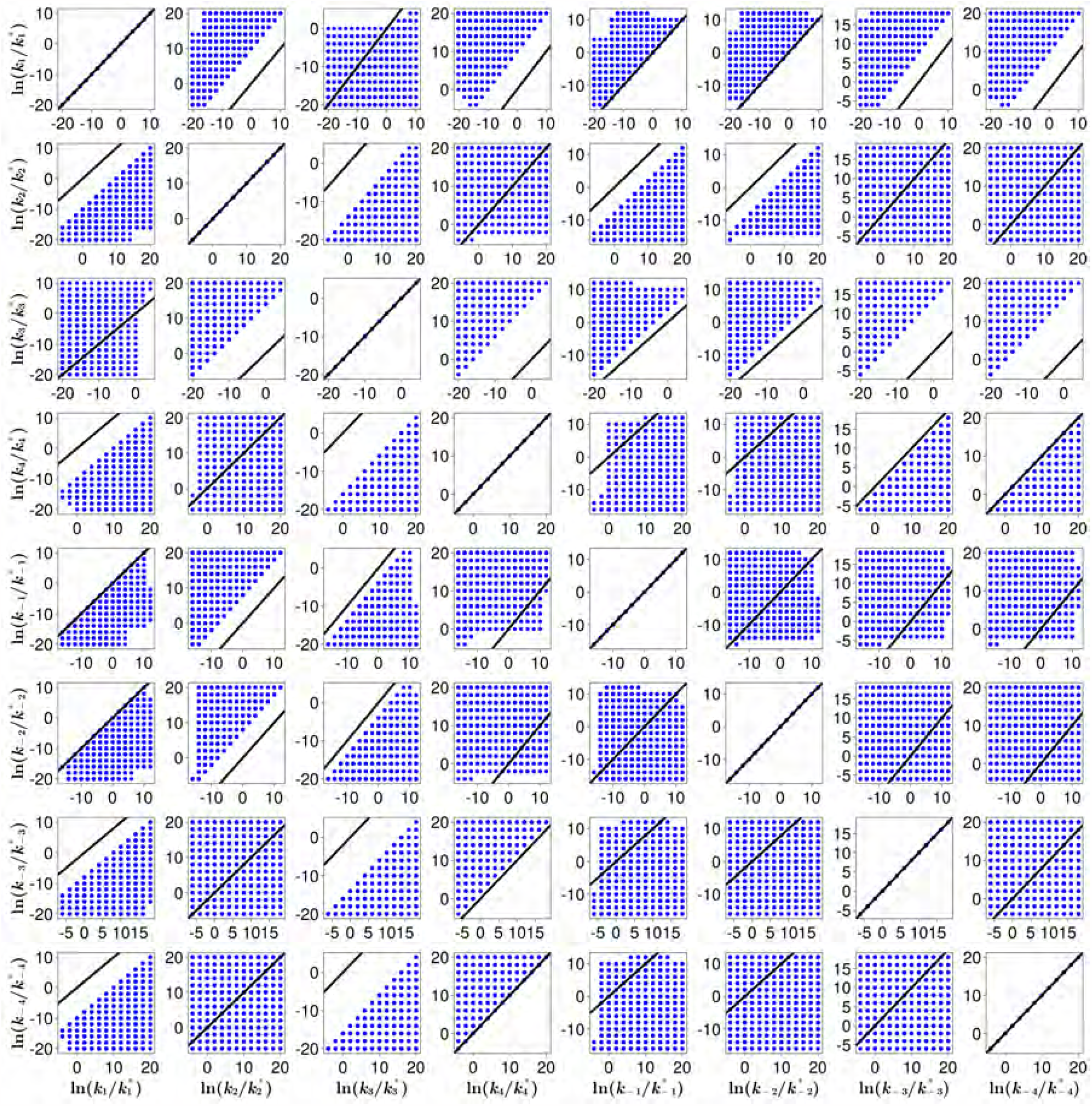
OR cluster 1: Rate relations between the fold change 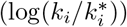 in each pair of the eight promoter switching rates for optimum gates operating anticlockwise for one of the clusters. Column labels (bottom) indicate the rate on the x-axis. Row labels (right) indicate the rates on the y-axis. Black line has slope = 1 and intercept = 0. Data was generated using the parameters and methods documented in Materials and Methods 4.1.2.

**Figure 12.**
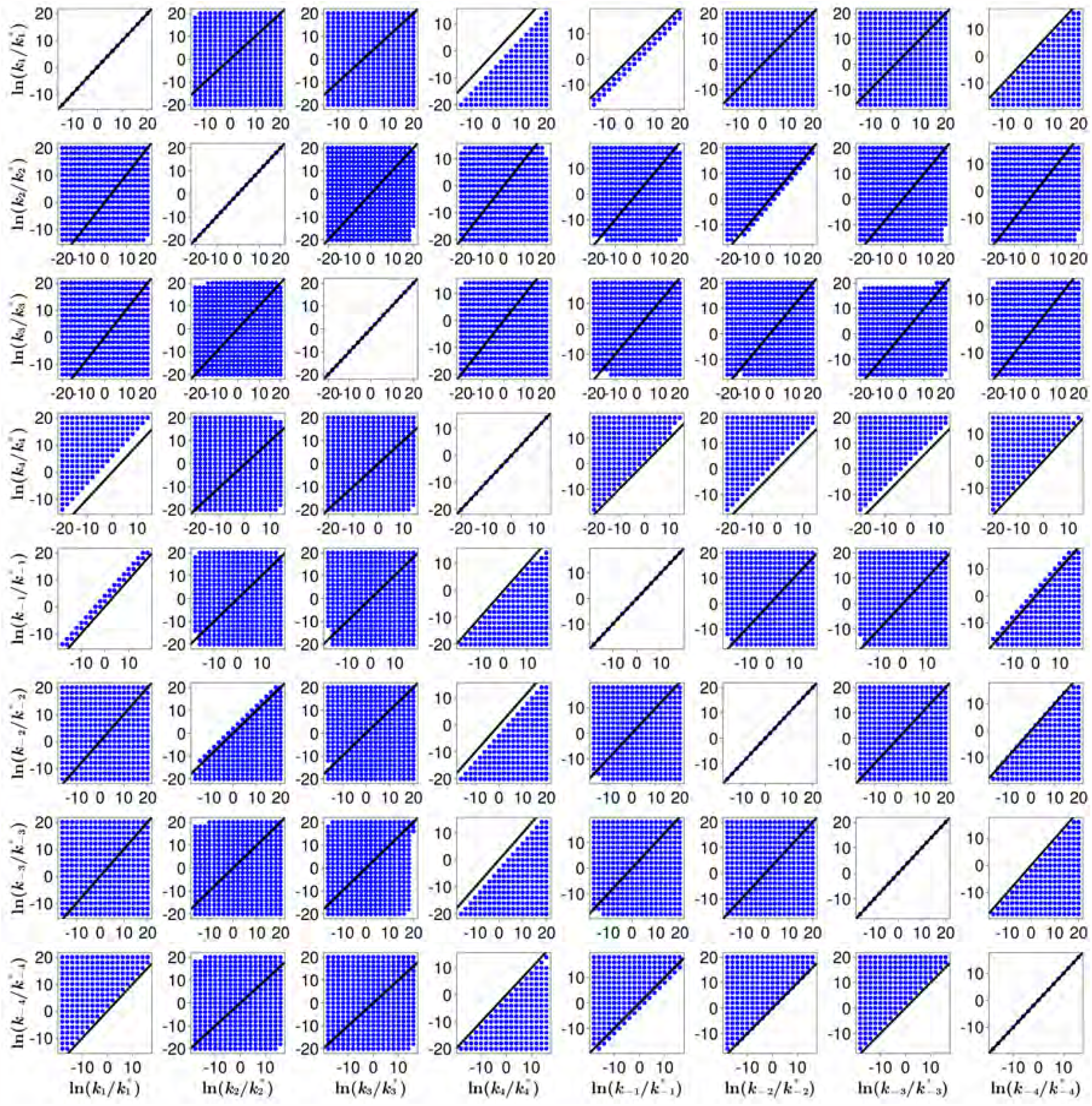
OR cluster 2: Rate relations between the fold change 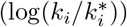 in each pair of the eight promoter switching rates for optimum gates operating anticlockwise for the second cluster. Column labels (bottom) indicate the rate on the x-axis. Row labels (right) indicate the rates on the y-axis. Black line has slope = 1 and intercept = 0. Data was generated using the parameters and methods documented in Materials and Methods 4.1.2.

**Figure 13.**
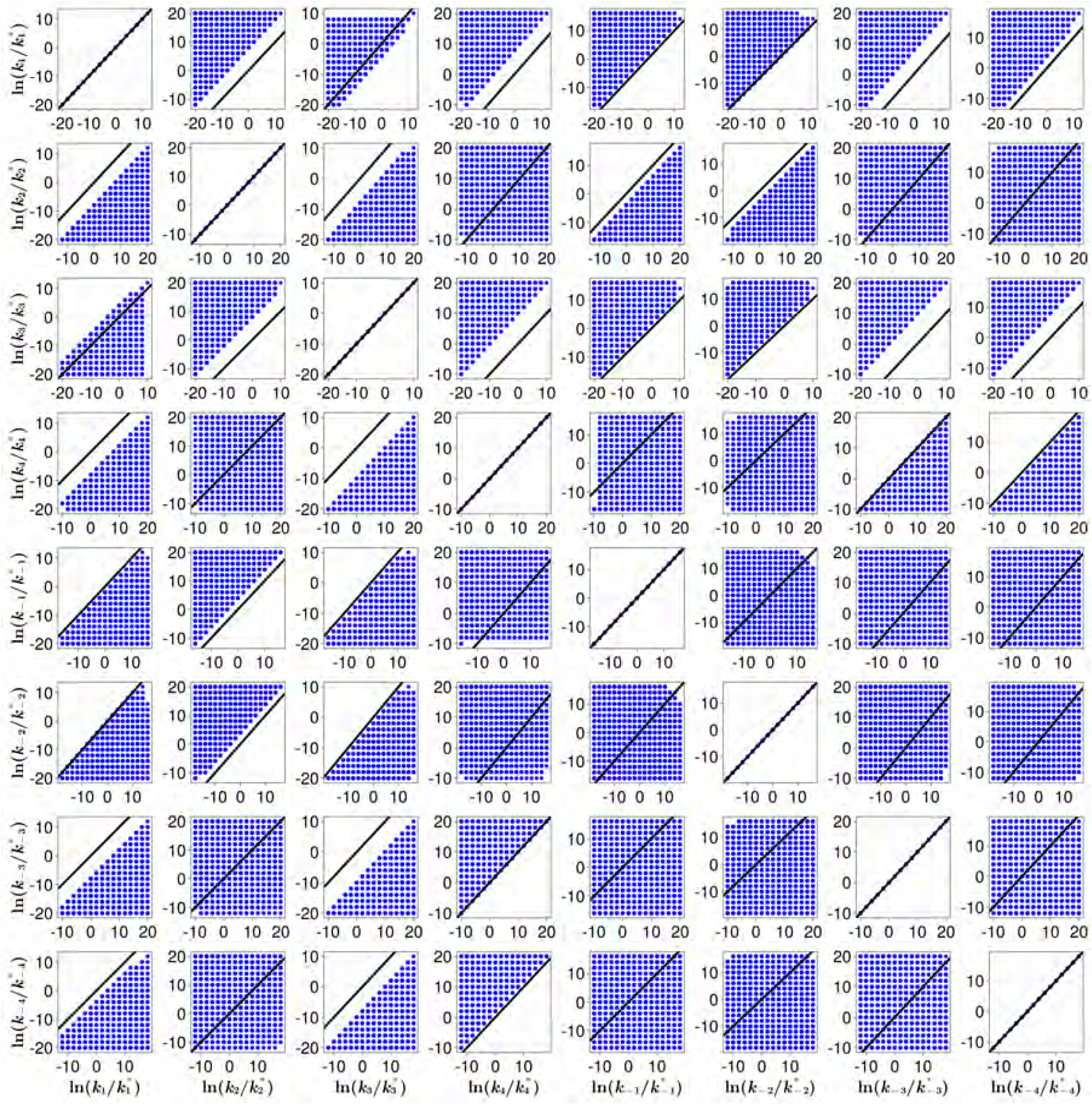
NAND: Rate relations between the fold change 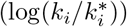 in each pair of the eight promoter switching rates for optimum gates operating anticlockwise. Column labels (bottom) indicate the rate on the x-axis. Row labels (right) indicate the rates on the y-axis. Black line has slope = 1 and intercept = 0. Data was generated using the parameters and methods documented in Materials and Methods 4.1.2.

**Figure 14.**
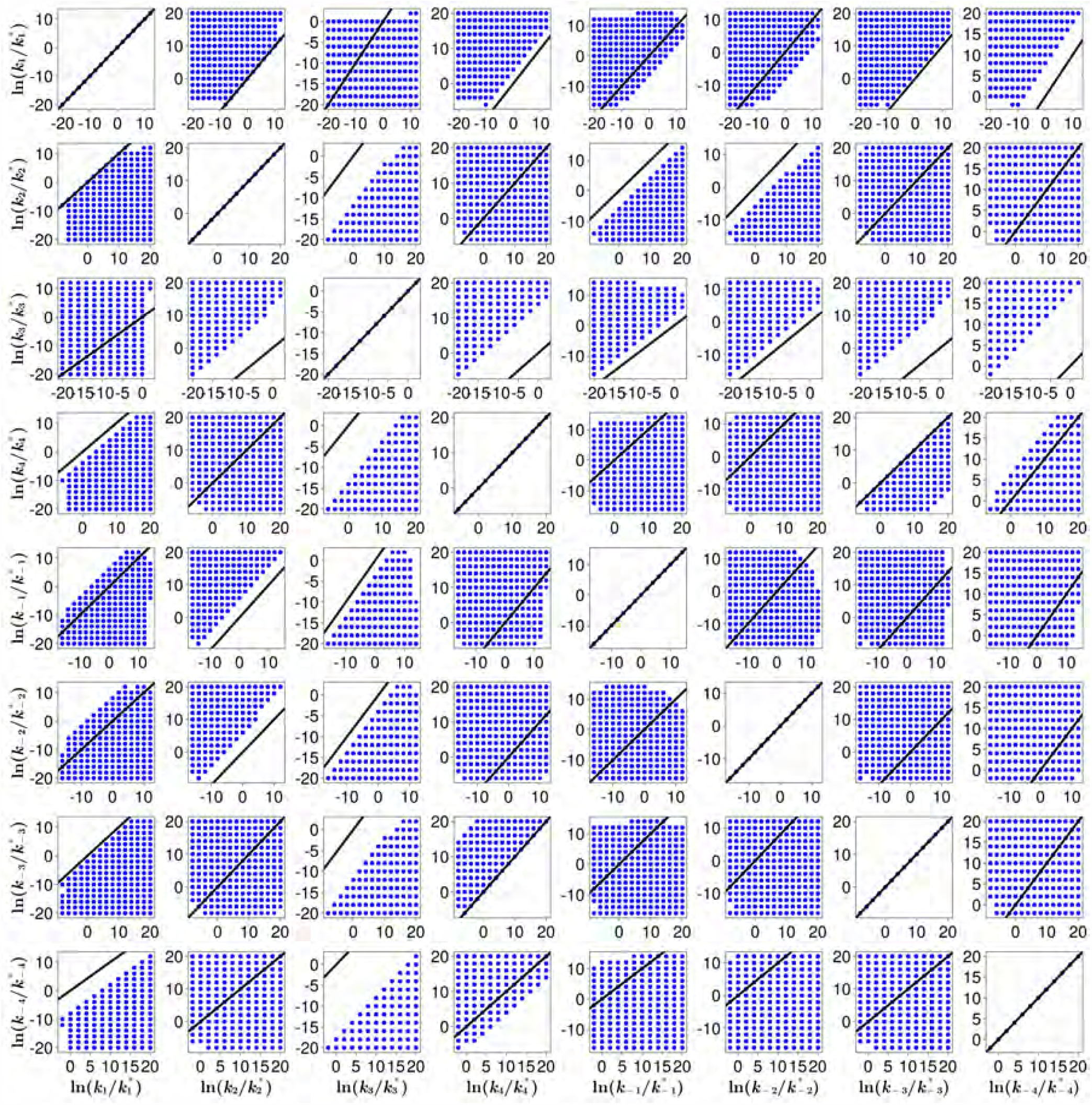
NOR: Rate relations between the fold change 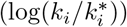 in each pair of the eight promoter switching rates for optimum gates operating anticlockwise. Column labels (bottom) indicate the rate on the x-axis. Row labels (right) indicate the rates on the y-axis. Black line has slope = 1 and intercept = 0. Data was generated using the parameters and methods documented in Materials and Methods 4.1.2.

##### 5.2 Fraction of gates in each regime

**Figure 15.**
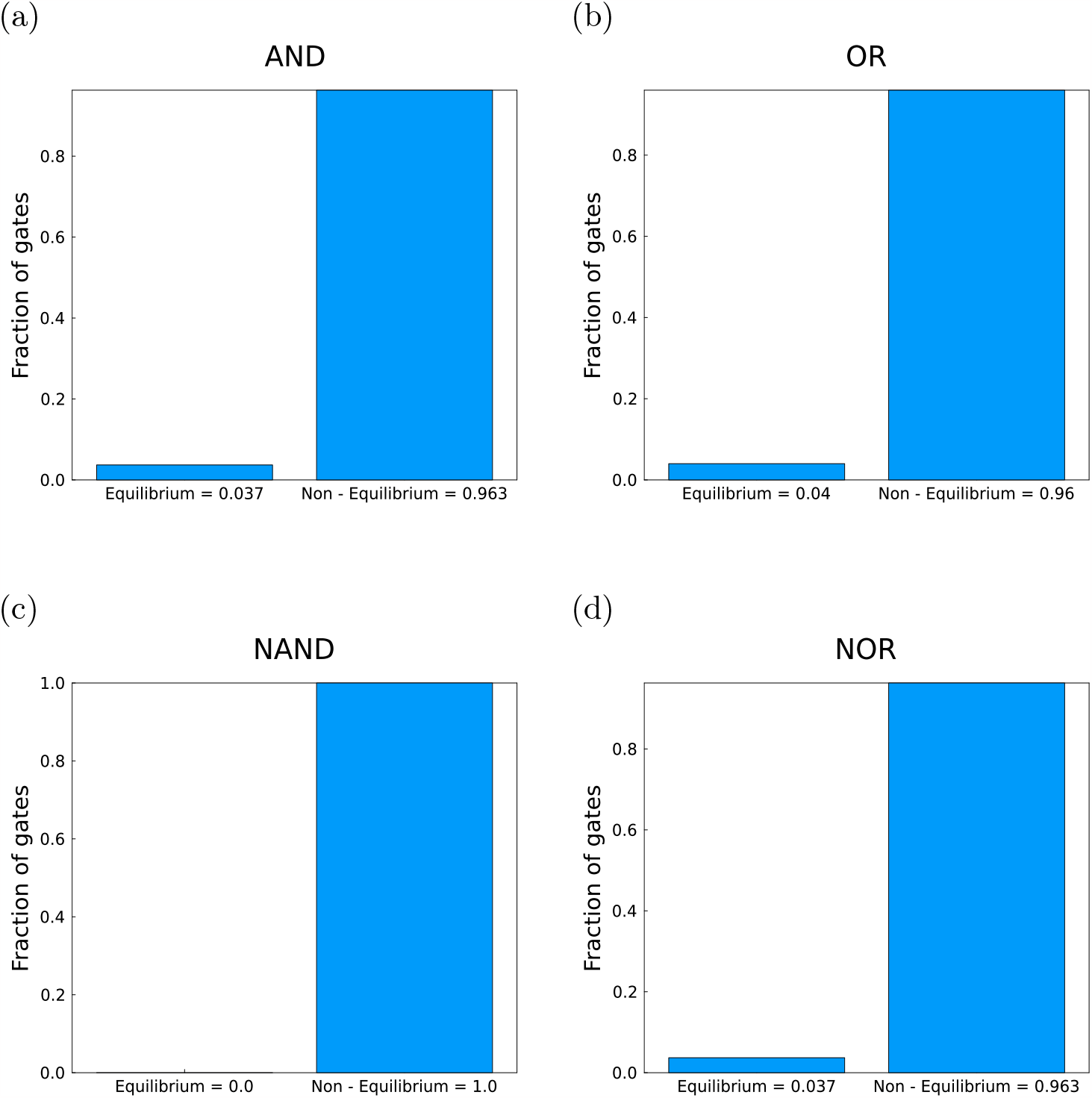
Fraction of the gates encountered in each regime. Data was generated using the methods and parameters described in Materials and Methods 4.1.2. Heights of the bars are provided in the x-axis labels. (a) AND, (b) OR, (c) NAND, (d) NOR.

#### 6 The *l*_1_ norm cutoff does not qualitatively affect the results for AND, NAND, OR and NOR gates

Data used for the following plots was generated by sweeping the parameter space spanned by 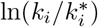 such that 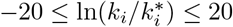 with increments of 2 for each promoter switching rate *k*_*i*_. The mRNA cutoff, ⟨*m*⟩, was taken as 5. The *l*_1_ norm cutoff was taken as 0.07. All other parameter values are the same as Table 7 of the Materials and Methods section. Non equilibrium gates were filtered out to plot the histograms in the following figures.

##### 6.1 Histograms of dissipated energy for gates with *l*_1_ norm lower than 0.07

**Figure 16.**
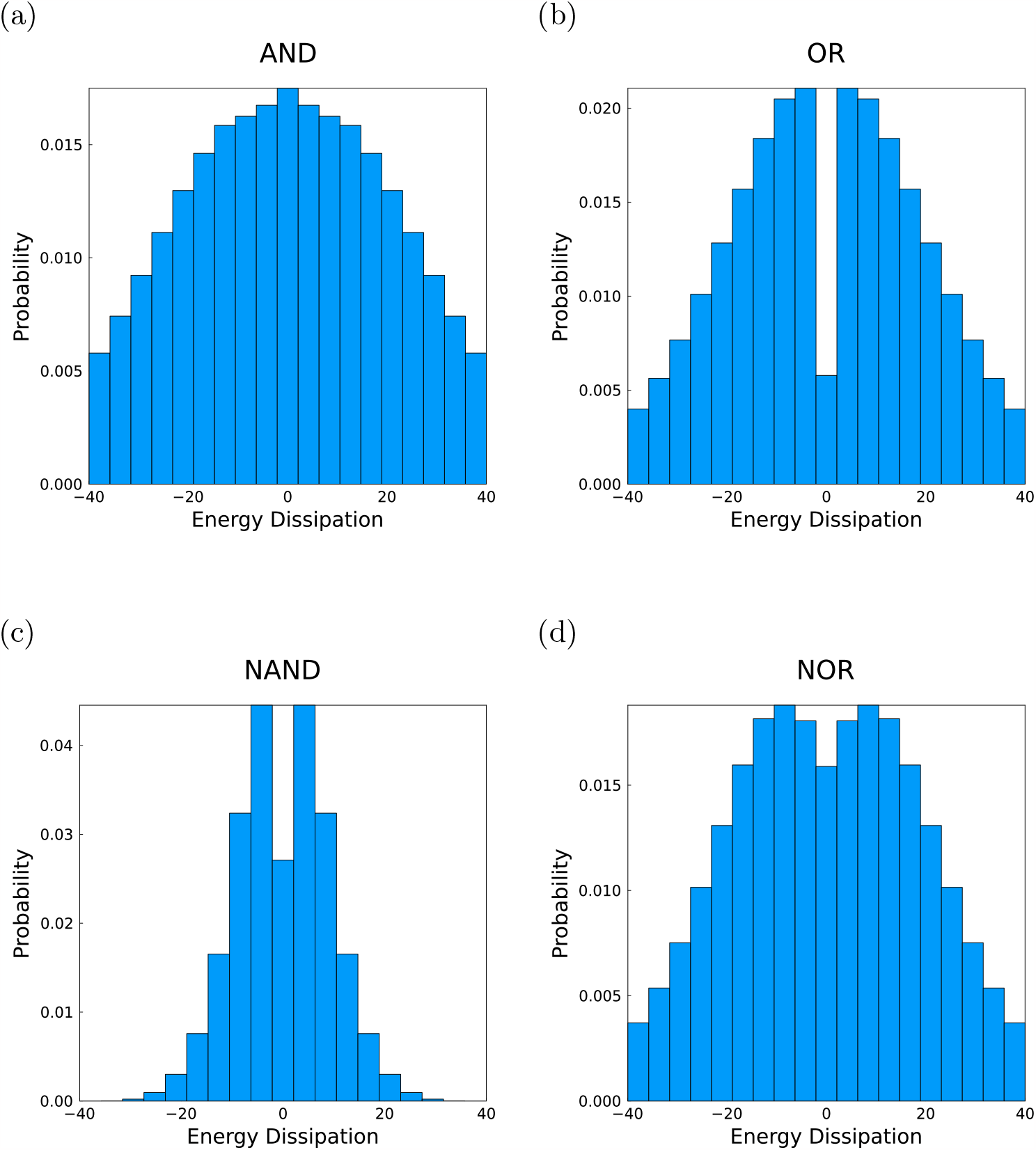
Energy dissipation in *k*_*B*_*T* for non-equilibrium gates with *l*_1_ norms below 0.07. (a) AND, OR, (c) NAND, (d) NOR.

##### 6.2 Fraction of gates in each regime with *l*_1_ cutoff = 0.07

**Figure 17.**
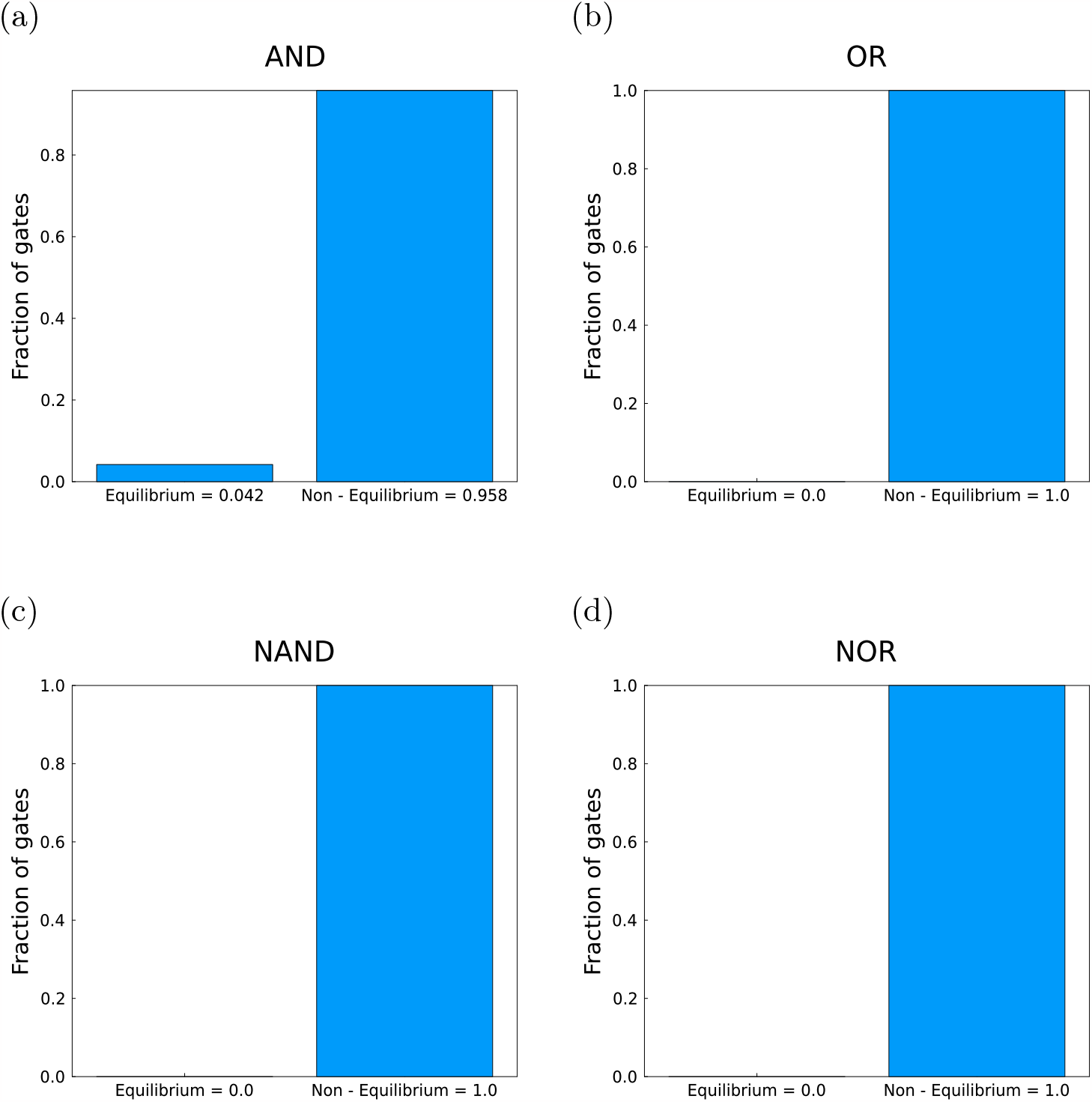
Fraction of the gates encountered in each regime with *l*_1_ norms below 0.07. Heights of the bars are provided in the x-axis labels. (a) AND, (b) OR, (c) NAND, (d) NOR.

#### 7 The mRNA cutoff does not affect the qualitative results of AND, NAND, OR and NOR

Data used for the following plots was generated by sweeping the parameter space spanned by 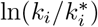 such that 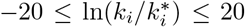 with increments of 4 for each promoter switching rate *k*_*i*_ in the case of AND,OR and NOR and increments of 2 in the case of NAND. The mRNA cutoff, ⟨*m*⟩, was taken as 1. The *l*_1_ norm cutoff was taken as 0.1. All other parameter values are the same as Table 7 of the Materials and Methods section. Non equilibrium gates were filtered out to plot the histograms in the following figures. For the rate relations, all anticlockwise gates with an *l*_1_ norm below the values documented in Table 8 were taken. In the case of OR gate, the rate relations were generated taking a slightly higher *l*_1_ norm cutoff of 0.075 to allow sufficient data for analysis. Clusters were assigned using Agglomerative Clustering from Python’s Sci-kit learn library.

##### 7.1 Histograms with mRNA cutoff = 1

**Figure 18.**
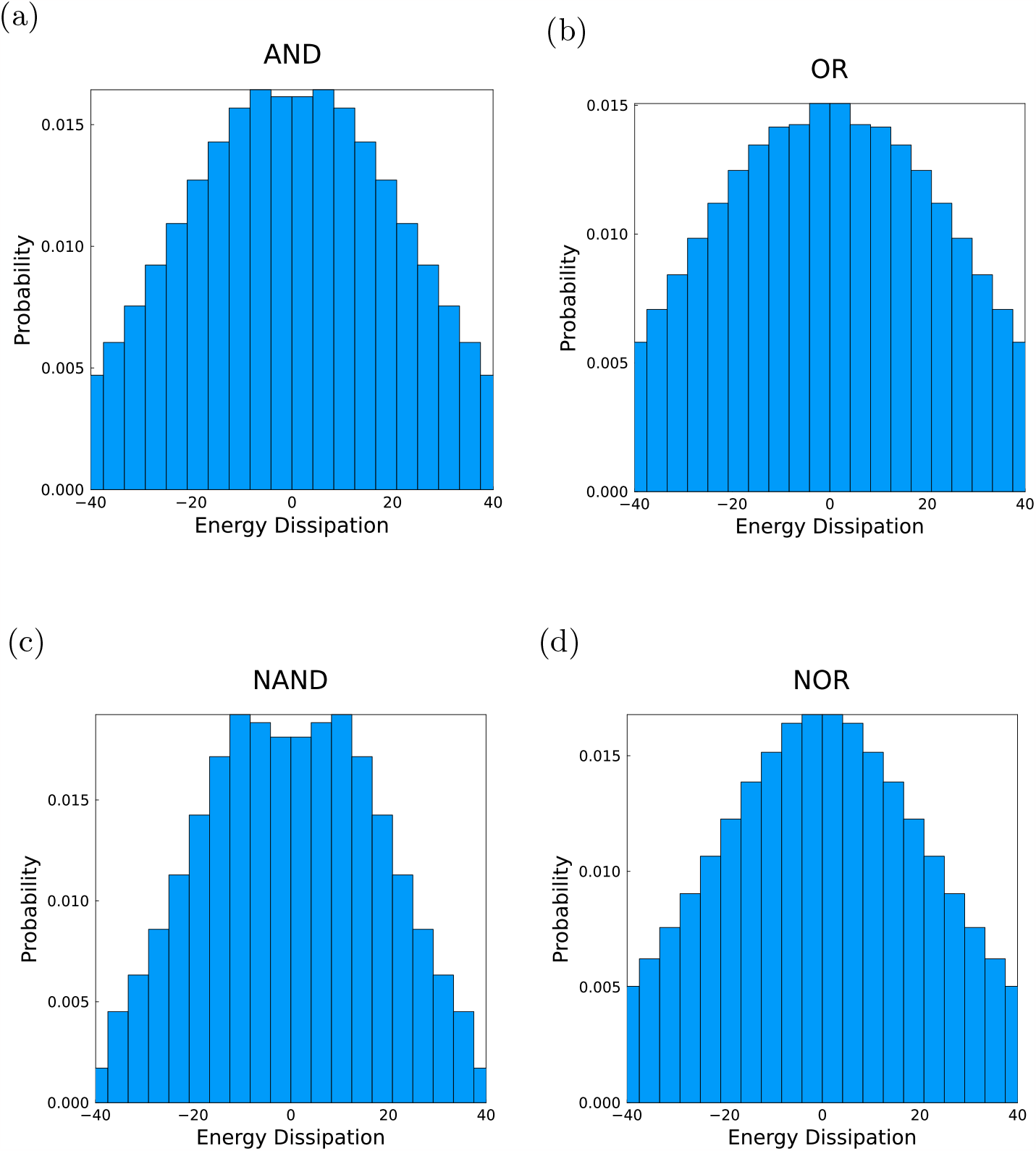
Energy dissipation in *k*_*B*_*T* for non-equilibrium gates with an mRNA cutoff at 1. (a) AND, (b) OR, (c) NAND, (d) NOR.

##### 7.2 Rate relations with mRNA cutoff = 1

**Figure 19.**
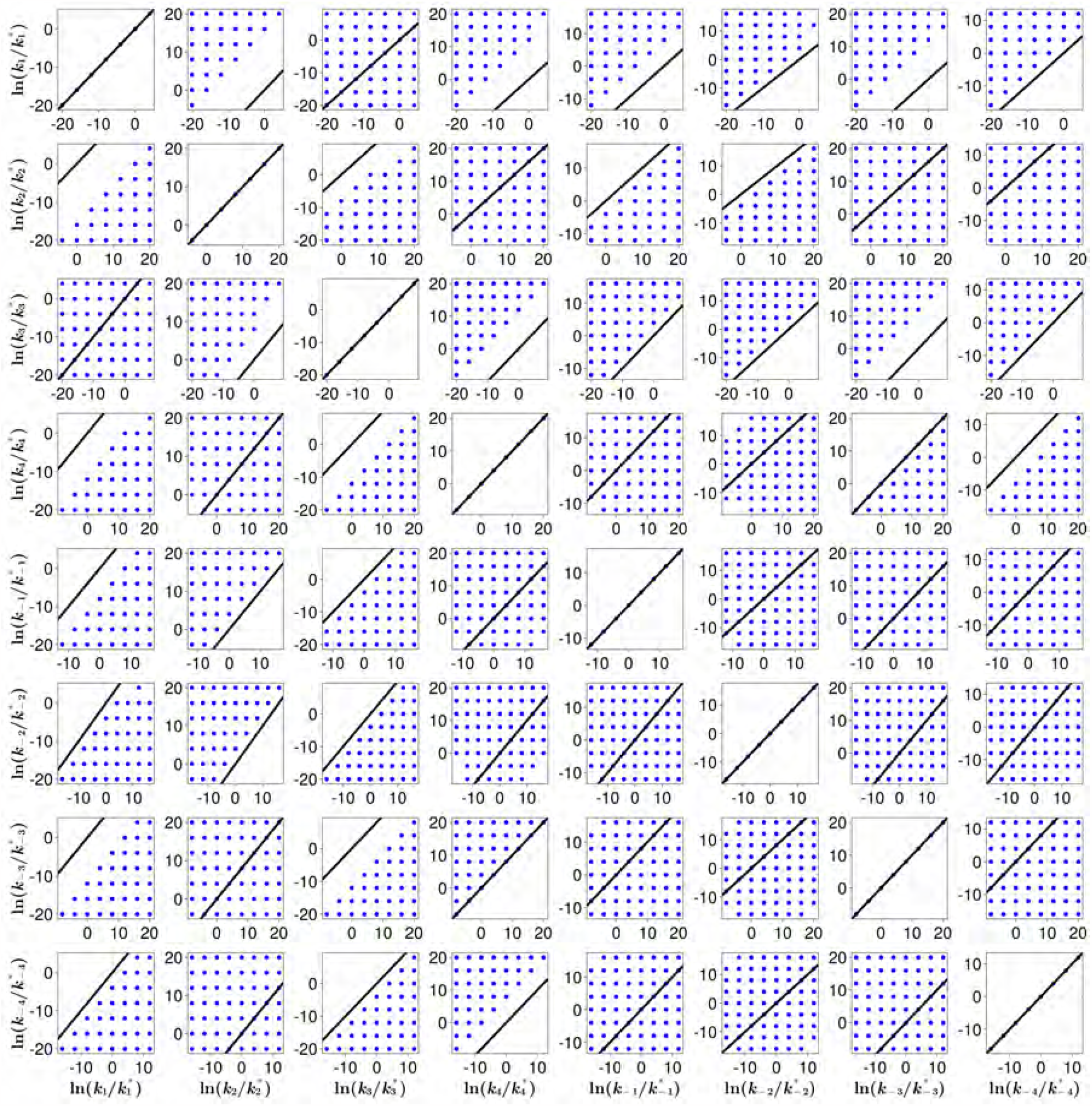
AND: Rate relations between the fold change 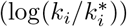 in each pair of the eight promoter switching rates for optimum gates operating anticlockwise. Column labels (bottom) indicate the rate on the x-axis. Row labels (right) indicate the rates on the y-axis. Black line has slope = 1 and intercept = 0. Data was generated using the methods described above.

**Figure 20.**
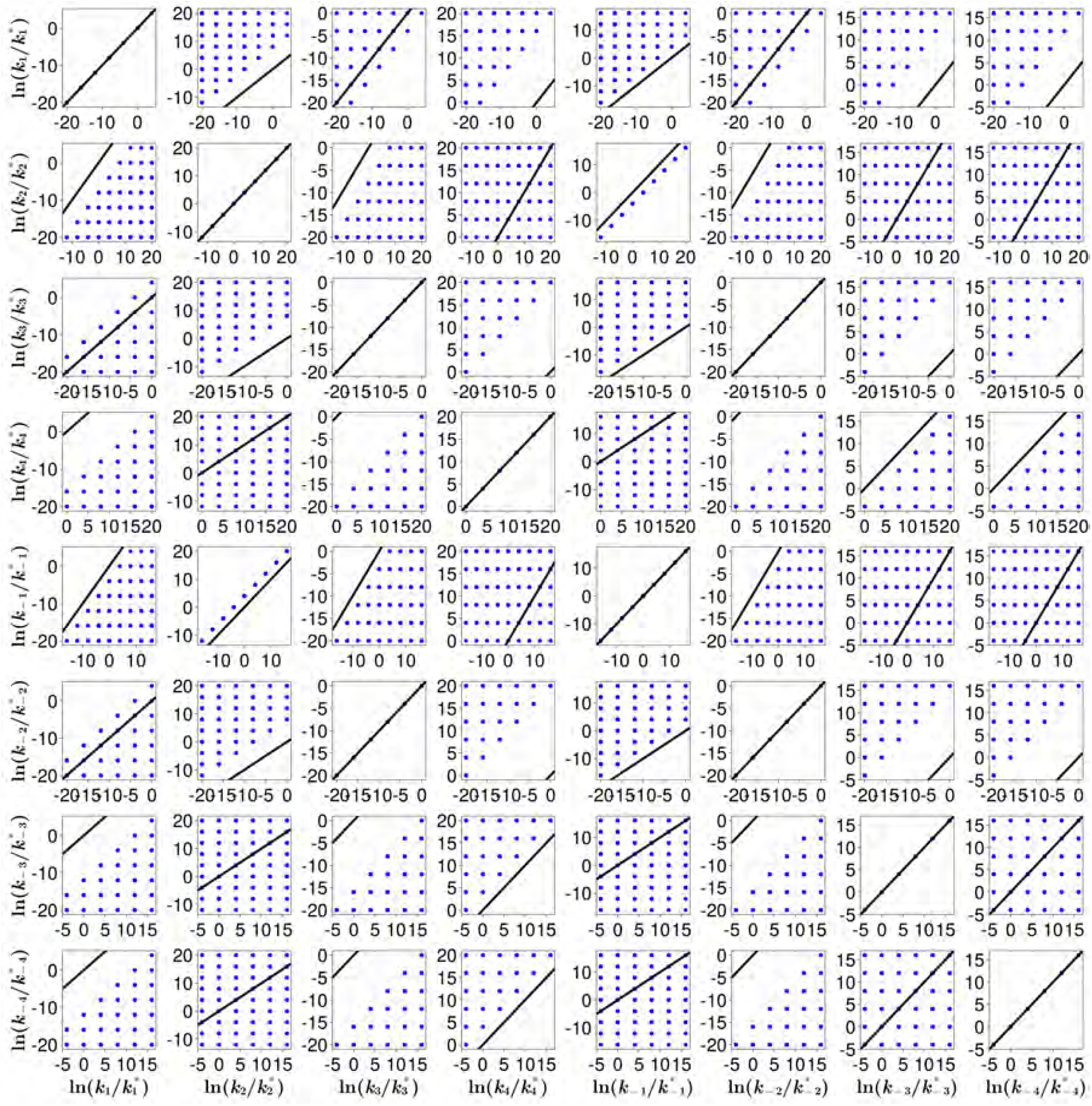
OR cluster 1: Rate relations between the fold change 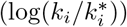 in each pair of the eight promoter switching rates for optimum gates operating anticlockwise for one of the clusters. Column labels (bottom) indicate the rate on the x-axis. Row labels (right) indicate the rates on the y-axis. Black line has slope = 1 and intercept = 0. Data was generated using the methods described above.

**Figure 21.**
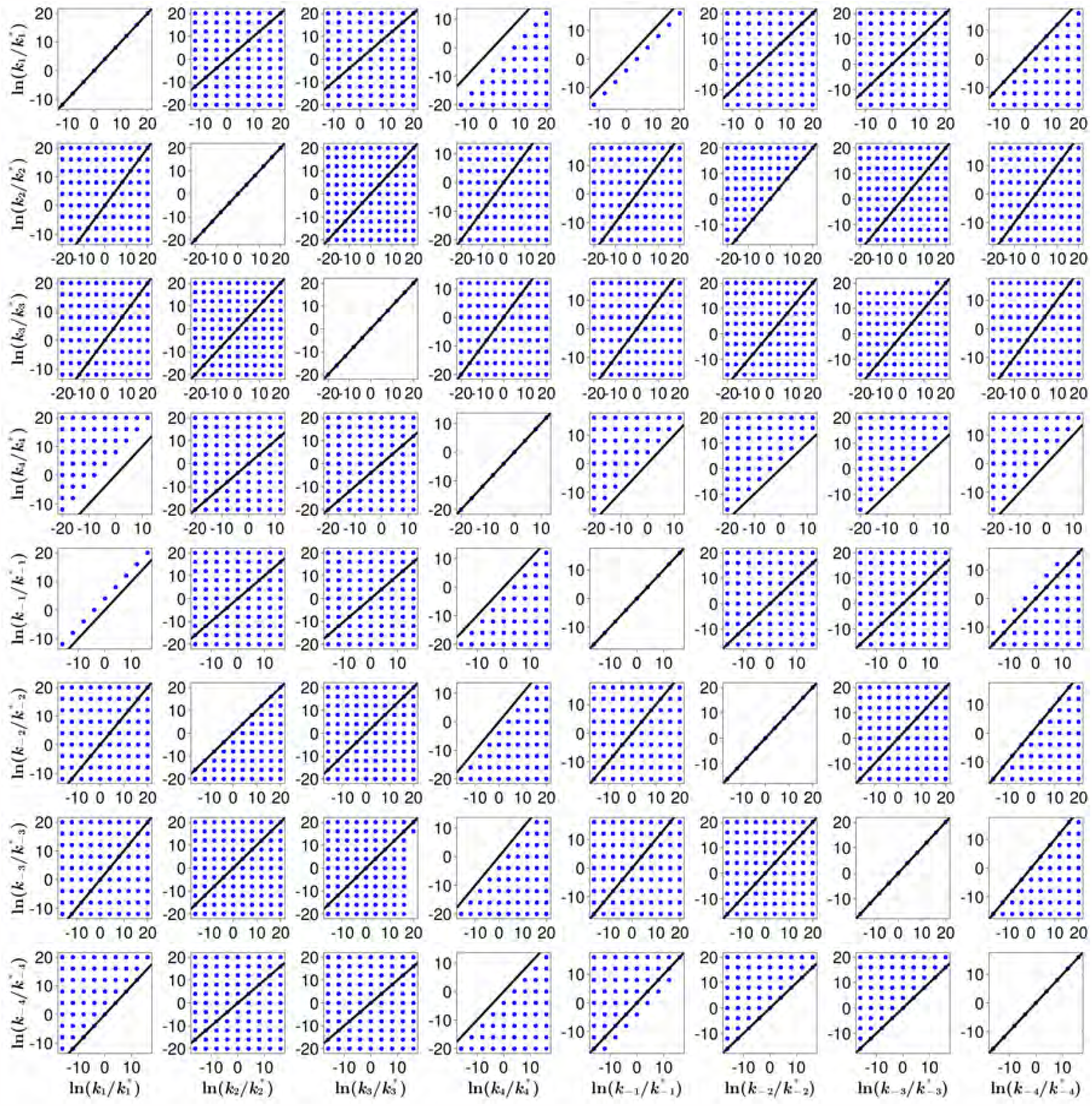
OR cluster 2: Rate relations between the fold change 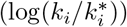 in each pair of the eight promoter switching rates for optimum gates operating anticlockwise for the second cluster. Column labels (bottom) indicate the rate on the x-axis. Row labels (right) indicate the rates on the y-axis. Black line has slope = 1 and intercept = 0. Data was generated using the methods described above.

**Figure 22.**
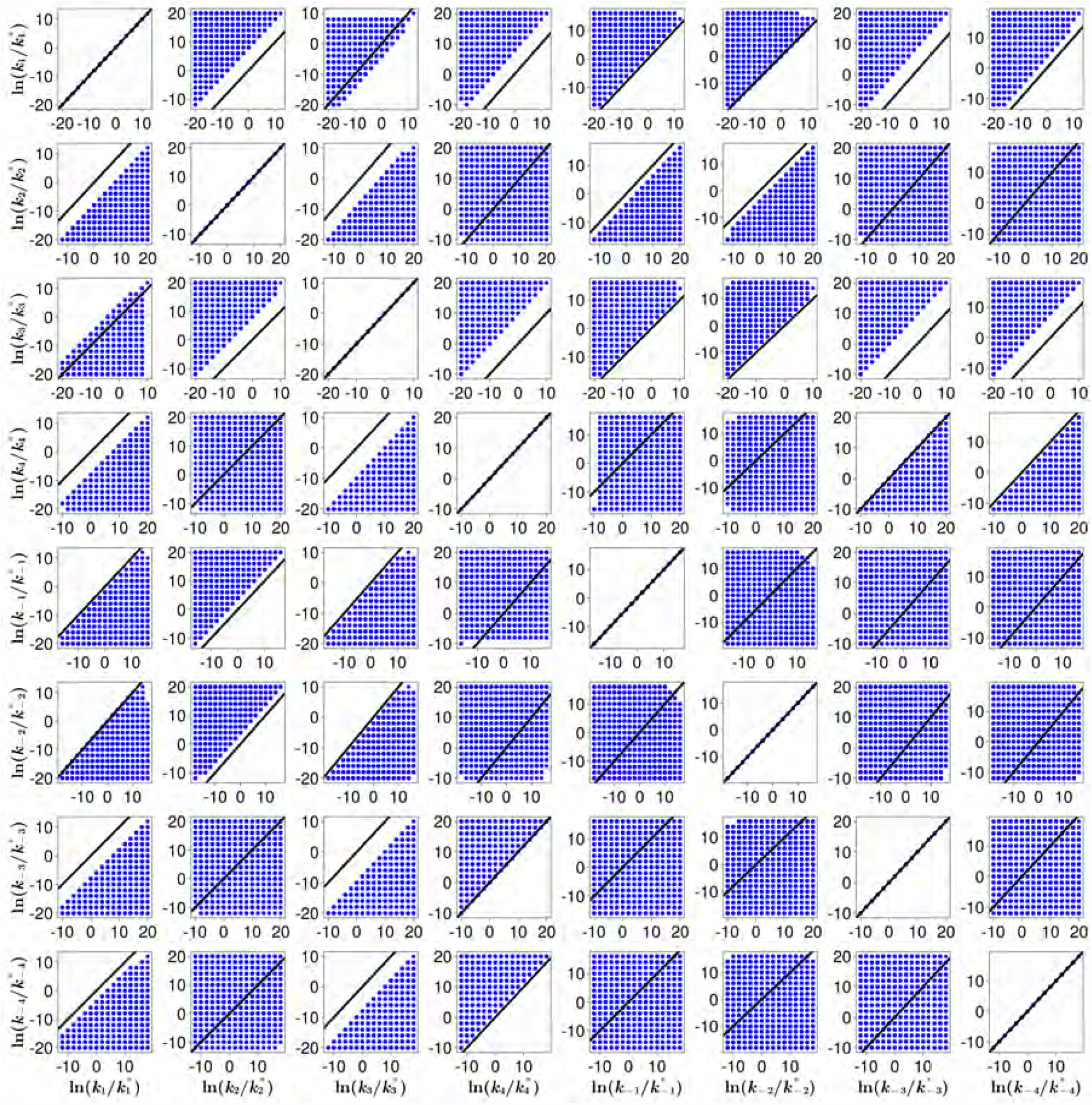
NAND: Rate relations between the fold change 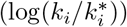 in each pair of the eight promoter switching rates for optimum gates operating anticlockwise. Column labels (bottom) indicate the rate on the x-axis. Row labels (right) indicate the rates on the y-axis. Black line has slope = 1 and intercept = 0. Data was generated using the methods described above.

**Figure 23.**
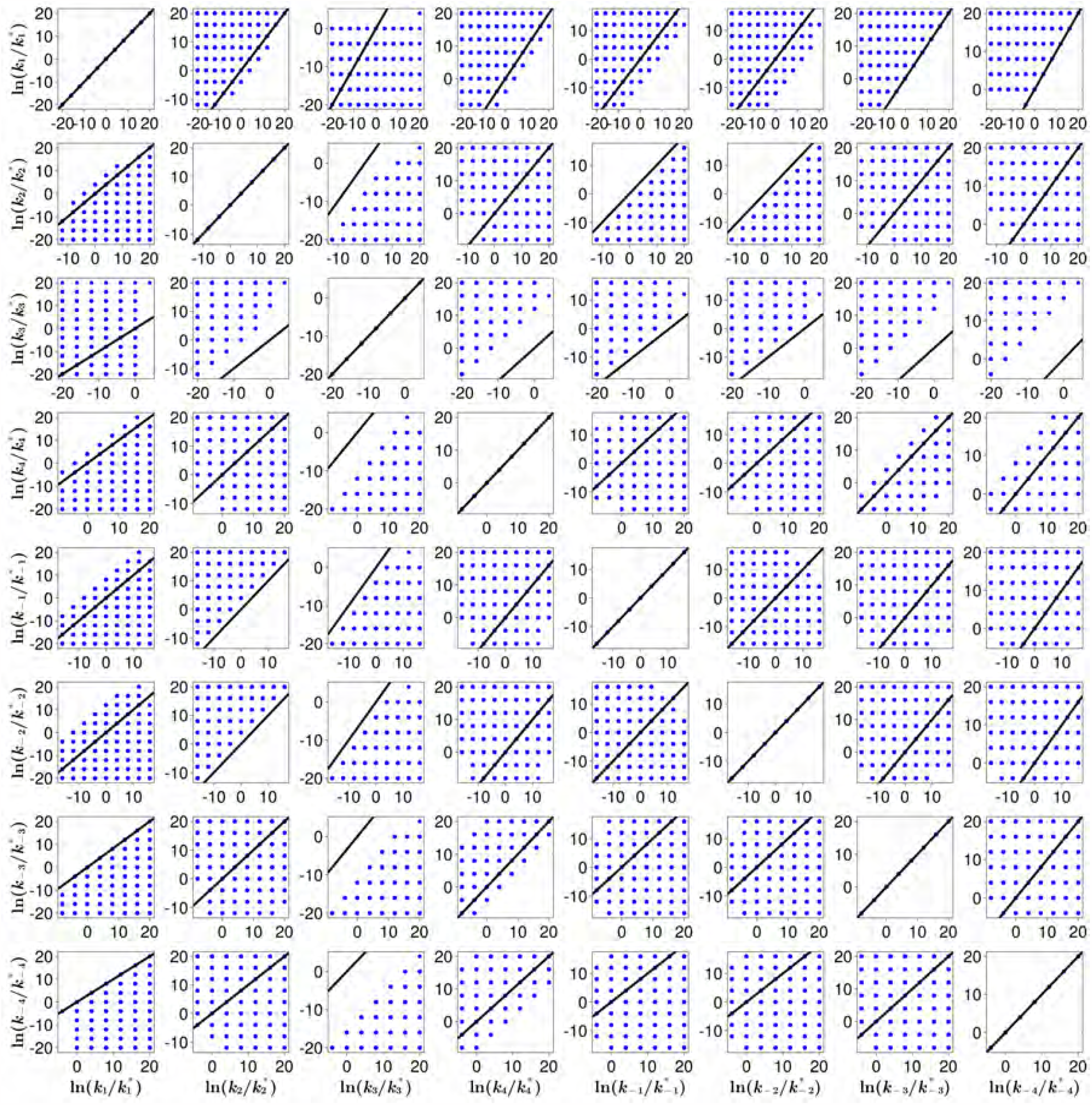
NOR: Rate relations between the fold change 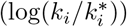 in each pair of the eight promoter switching rates for optimum gates operating anticlockwise. Column labels (bottom) indicate the rate on the x-axis. Row labels (right) indicate the rates on the y-axis. Black line has slope = 1 and intercept = 0. Data was generated using the methods described above.

##### 7.3 Fraction of gates in each regime with mRNA cutoff = 1

**Figure 24.**
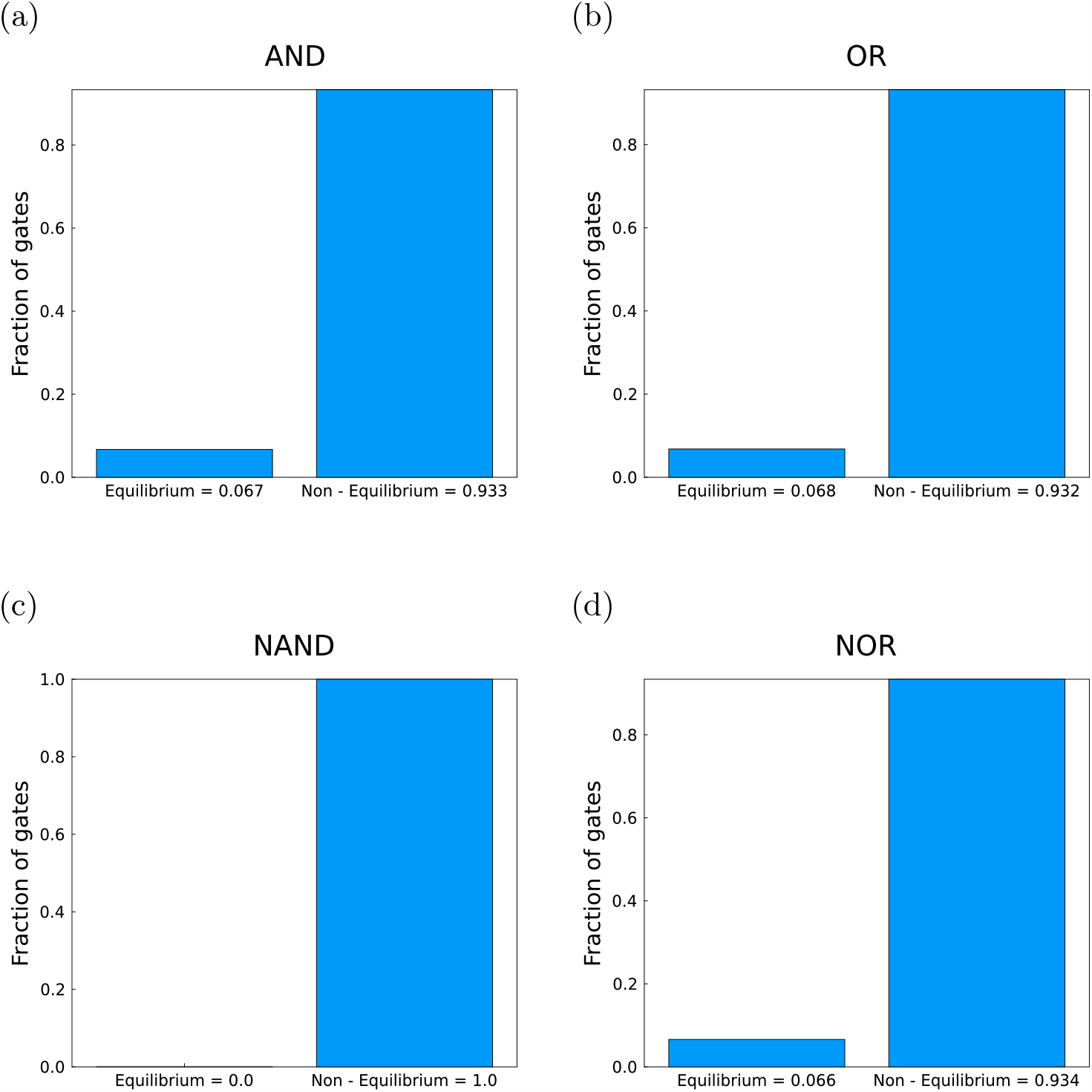
Fraction of the gates encountered in each regime with an mRNA cutoff of 1. Heights of the bars are provided in the x-axis labels. (a) AND, (b) OR, (c) NAND, (d) NOR.

#### 8 LAC and IMP gates can be made with different activation strategies

Data used for the following plots was generated by sweeping the parameter space spanned by 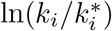 such that 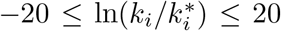 with increments of 2 for each promoter switching rate *k*_*i*_ The *l*_1_ norm cutoff was taken as 0.1. All other parameter values are the same as Table 7 of the Materials and Methods section. The rates of transcription from each promoter state *r*_*i*_ depend on the gene expression strategies and are documented in the captions.

**Figure 25.**
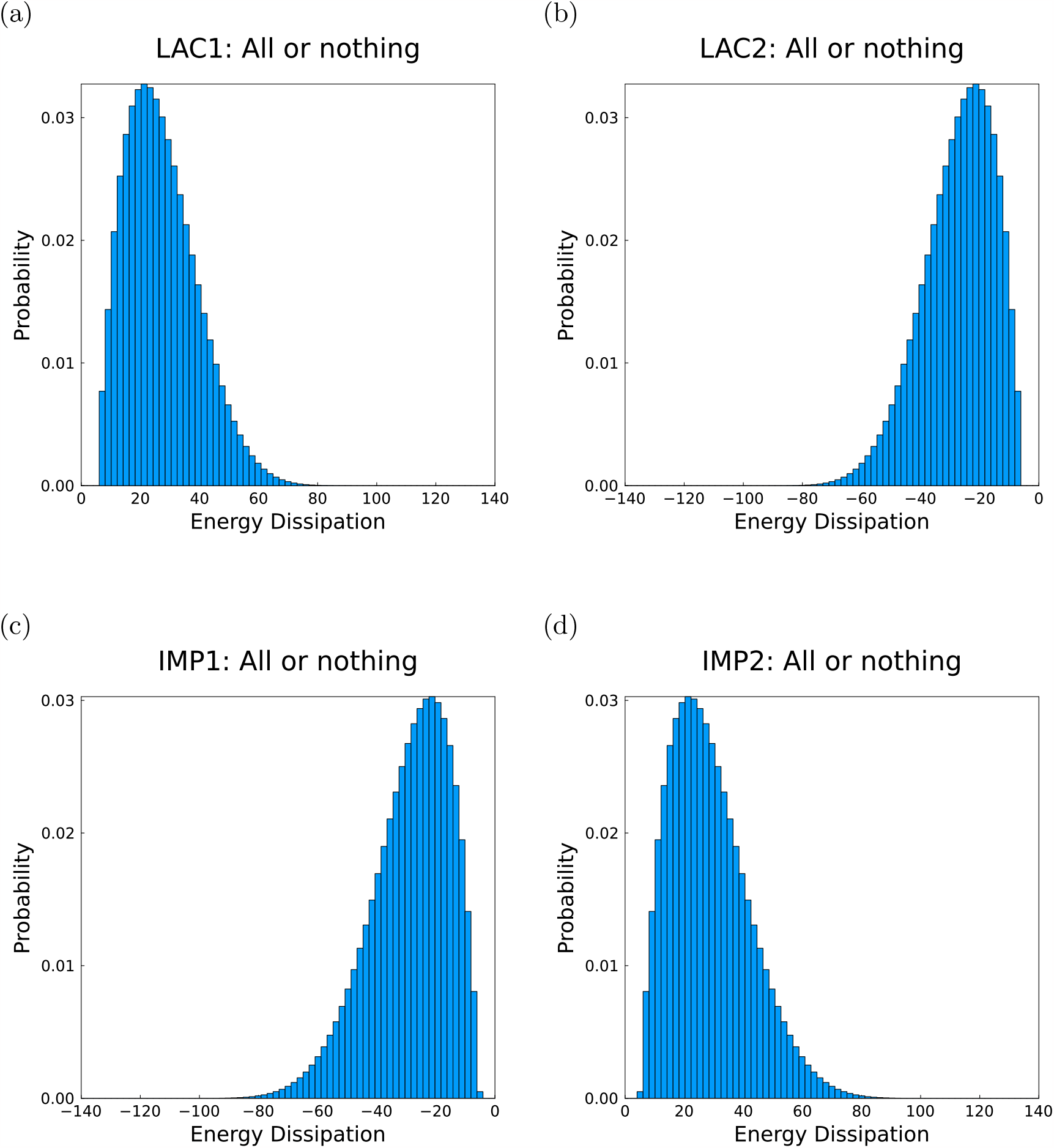
Energy dissipation in *k*_*B*_*T* for gates using an all - or - nothing strategy. *r*_1_ = *r*_2_ = *r*_4_ = 0, *r*_3_ = 0.3*s*^−1^. (a) LAC1, (b) LAC2, (c) IMP1, (d) IMP2.

**Figure 26.**
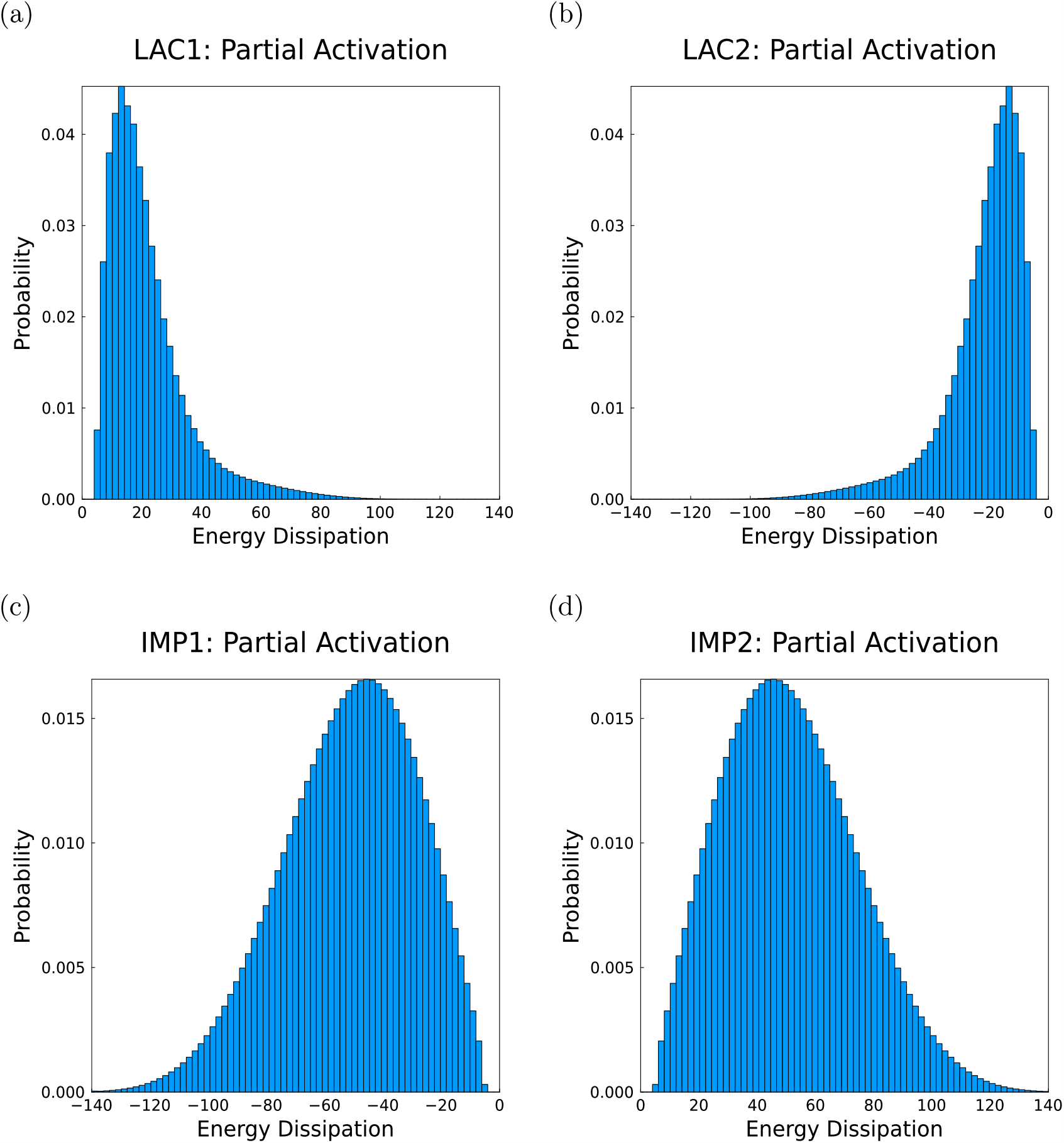
Energy dissipation in *k*_*B*_*T* for gates using a partial activation strategy. *r*_1_ = 0, *r*_2_ = *r*_4_ = 0.15*s*^−1^, *r*_3_ = 0.3*s*^−1^. (a) LAC1, (b) LAC2, (c) IMP1, (d) IMP2.

## Proofs for non-monotonicity of the promoter state occupancies as a function of TF concentrations

First, we need to define the weights and the promoter state occupancies. We set *k*_-1_ = k5, k6 =*k*_-2_, k7 =*k*_-3_, k8 = *k*_-4_ (anticlockwise rates). For the purpose following analysis, A corresponds to the concentration of TF A and B corresponds to the concentration of TF B.

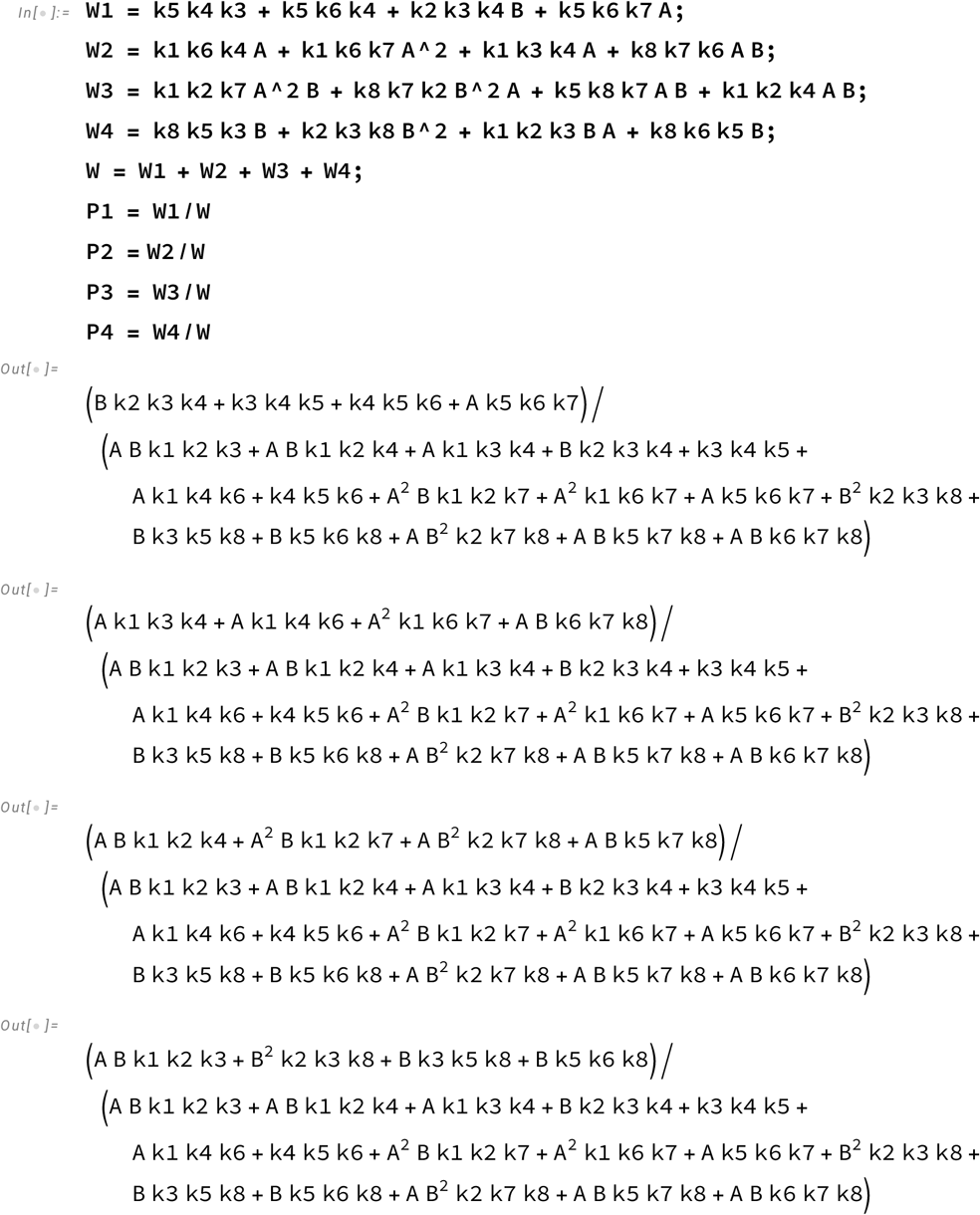

## Promoter state 1 exhibits non-monotonicity with respect to one of the TFs only

We first consider of the occupancy of promoter state 1, *P*_1_. For the rest of the promoter states, the proofs follow in a similar manner. For non-monotonicity with *A*, we need the 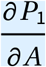 to change sign at least once for*A* > 0.

***In[* . ] : *=*** `DelP1A` **=** `D`[`P1, A`**] //** `FullSimplify`

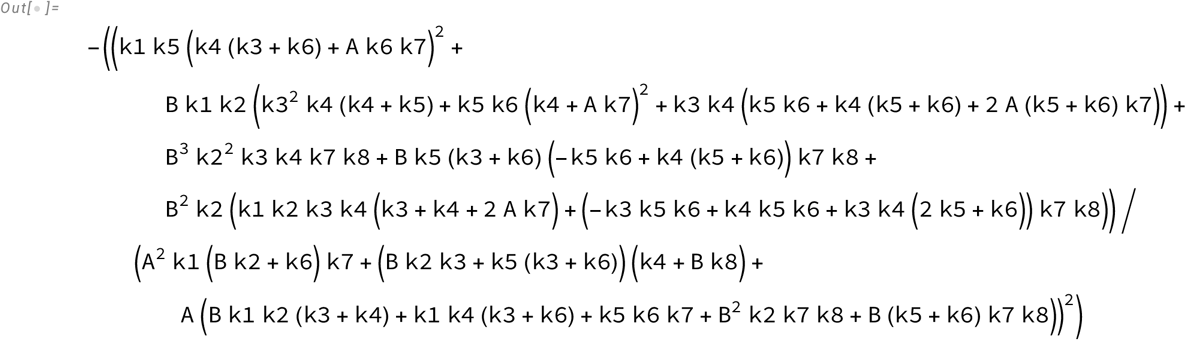

Note that the Denominator:

(A^2^ k1 (B k2 + k6) k7 + (B k2 k3 + k5 (k3 + k6) (k4 + B k8)+ is always positive A (B k1 k2 (k3 + k4) + k1 k4 (k3 + k6) + k5 k6 k7 + B^2^ k2 k7 k8 + B (k5 + k6) k7 k8))^2^ and never changes sign. We consider the numerator,

***In[* . ] : *=*** `NumDelP1A` **=** `Collect`[`Numerator`[`DelP1A`**] //** `Expand, A`]

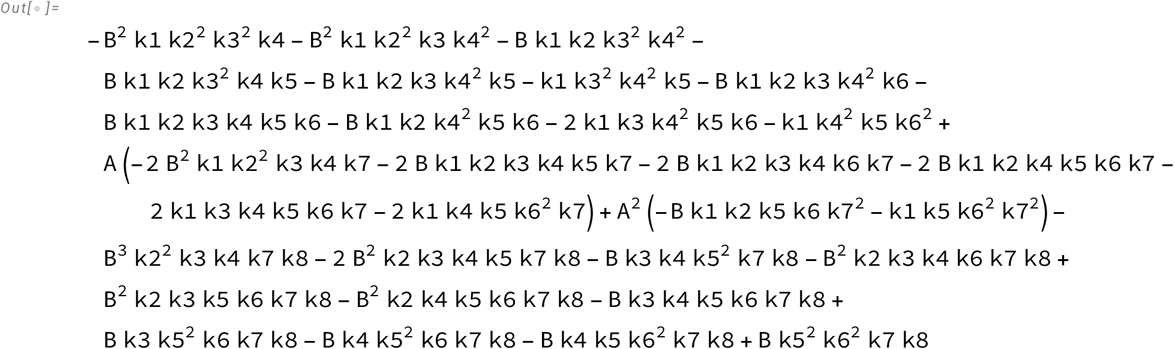

The numerator is a quadratic in *A* of the form *a A*^2^ + *b A* + *c*, where

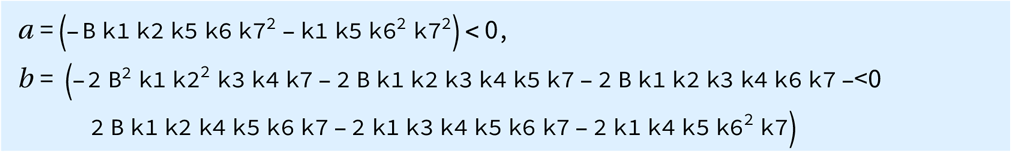

Therefore, for a real root in the positive *A* region (and consequently a sign change in 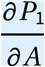, we need *c* > 0 and *b*^2^ - 4 *a c* > 0. The first condition is a strict one, sufficient and necessary, it automatically guarantees *b*^2^ - 4 *a c* > 0. The set of promoter switching rates that satisfy *c* > 0 is a subset of the promoter switching rates that ensure *b*^2^ - 4 *a c* > 0. We first solve for *c* > 0.

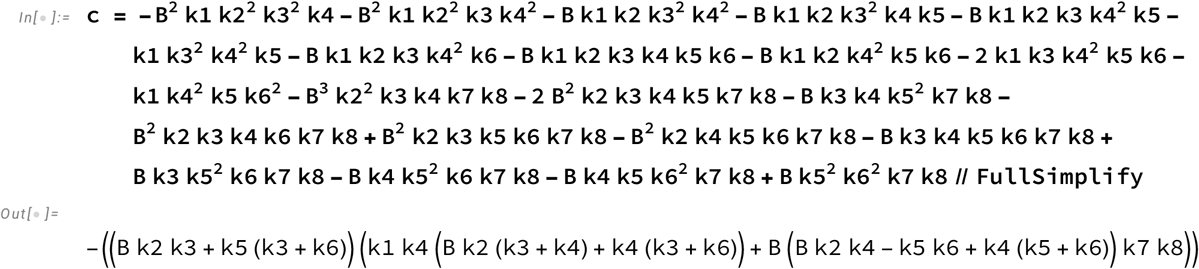

Now, we look at the conditions for which we can find *c* > 0. We do this using Mathematica’s Reduce function that provides regions in our eight-dimensional space where the inequality holds.

***In[* . ] : *=*** `Reduce`[`c` **>** **0**, **{**`k1, k2, k3, k4, k5, k6, k7, k8`**}**, **PositiveReals**]

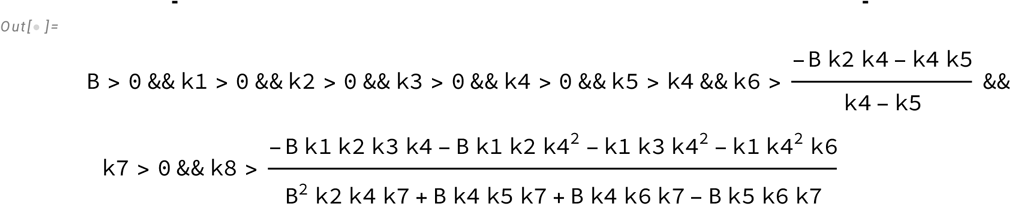

These are strict conditions that are difficult to interpret. We look at the necessary condition, that is *b*^2^ - 4 *a c* > 0.

***In[* . ] : *=*** `DisP1A` **=**

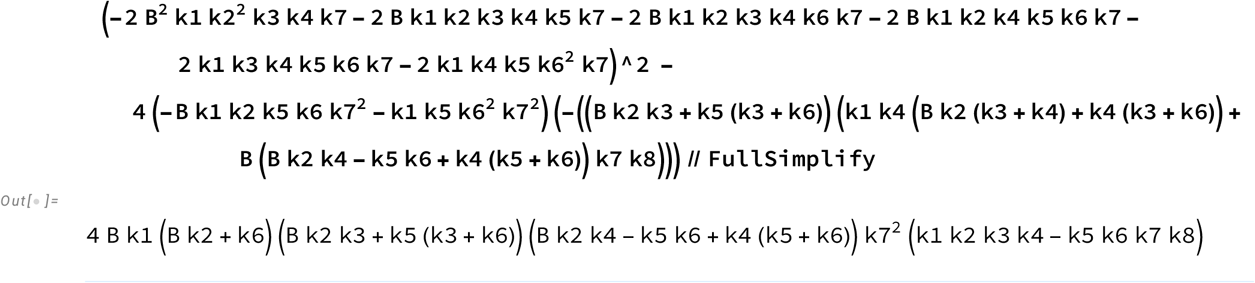

We find the conditions under which *b*^2^ - 4 *a c* > 0

***In[* . ] : *=*** `Reduce`;[`DisP1A` **>** **0**, **{**`k1, k2, k3, k4, k5, k6, k7, k8`**}**, **PositiveReals**]

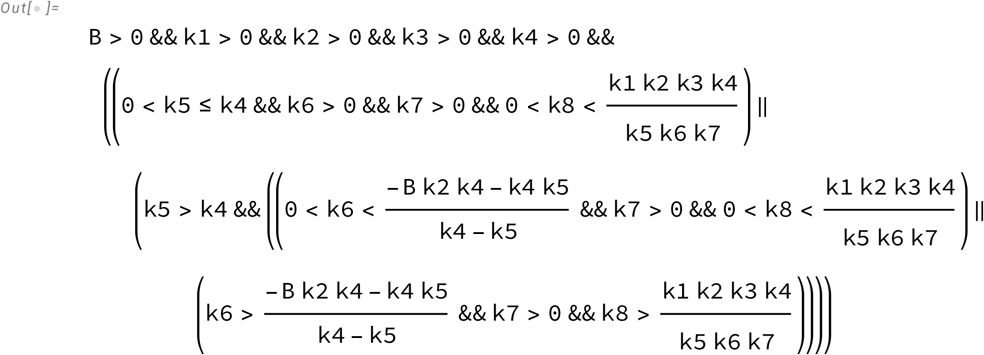

This is a soft condition that is necessary but insufficient to guarantee non-monotonicity. But from our requirement that *c* > 0, we know where the space of parameters satisfying *c* > 0 *and b*^2^ - 4 *a c* > 0 overlap. Namely, we have *k* 5 > *k* 4 and 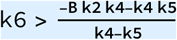. Therefore, the necessary condition we find from solving for 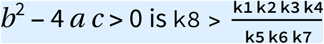.

This implies that for non-monotonicity in A, we will need the cycle to operate **anticlockwise** such that `k8 k5 k6 k7` > `k1 k2 k3 k4`.

Now we look for non-monotonicity of *P*_1_ with respect to B employing the same logic. We find 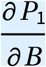 as

***In[* . ] : *=*** `DelP1B` **=** `D`[`P1, B`**] //** `FullSimplify`

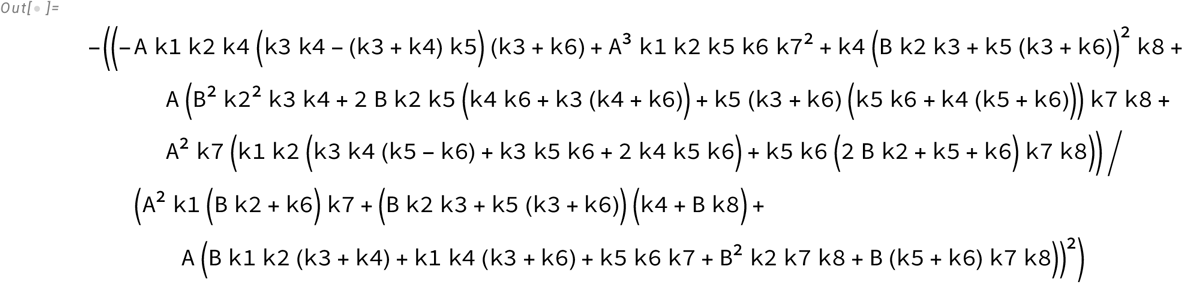

The denominator is always positive:

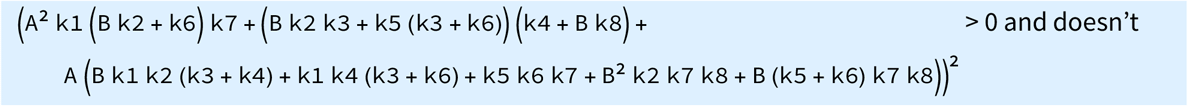

change sign. So we consider the numerator.

***In[* . ] : *=*** `NumDelP1B` **=** `Collect`[`Numerator`[`DelP1B`**] //** `Expand, B`]

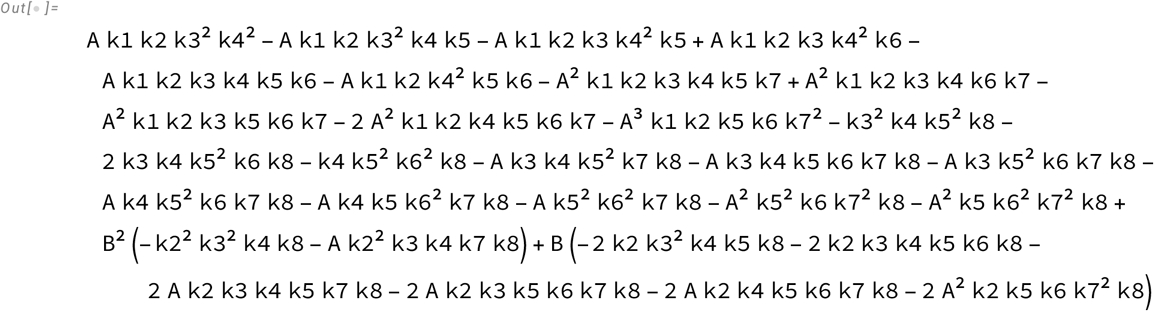

The numerator is a quadratic in B and has the form *a B*^2^ + *b B* + *c*. For a sign change we need a real root in *B* > 0. Note that as before

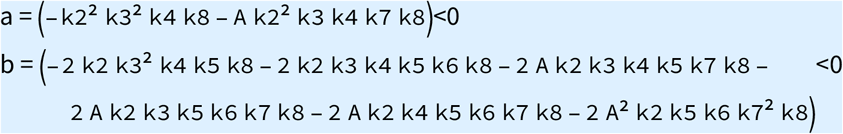

So we need *c* > 0. We solve for the regions in the eight-dimensional space where this condition holds true.

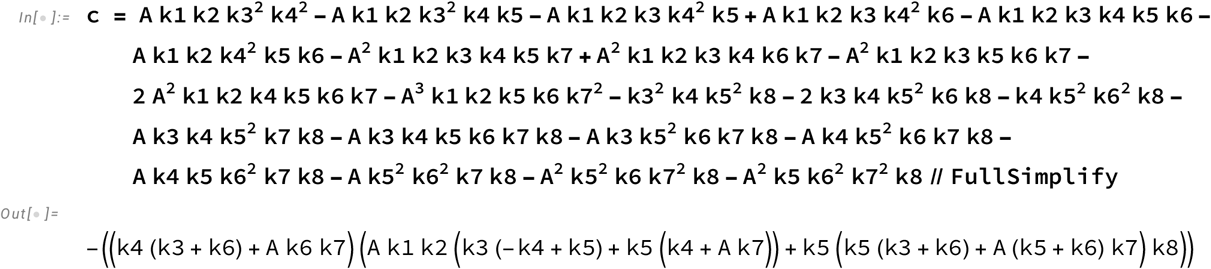

We look for conditions for *c* > 0

*In[* . ] : *=* `Reduce`[`c` **>** **0**, **{**`k1, k2, k3, k4, k5, k6, k7, k8`**}**, **PositiveReals]**

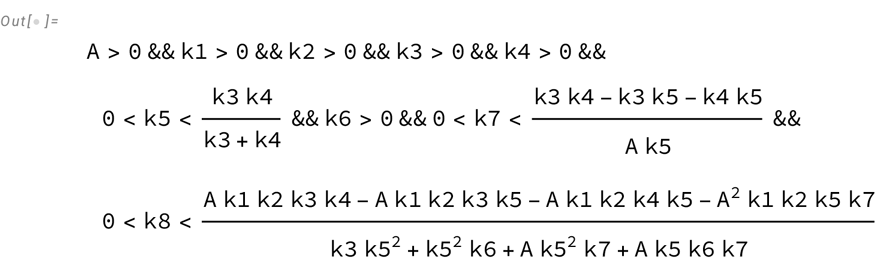

As before, we have the necessary and sufficient conditions that are difficult to interpret. We look for conditions for *b*^2^ - 4 *a c* > 0 to find where the spaces overlap.

***In[* . ] : *=*** `DisP1B` **=**

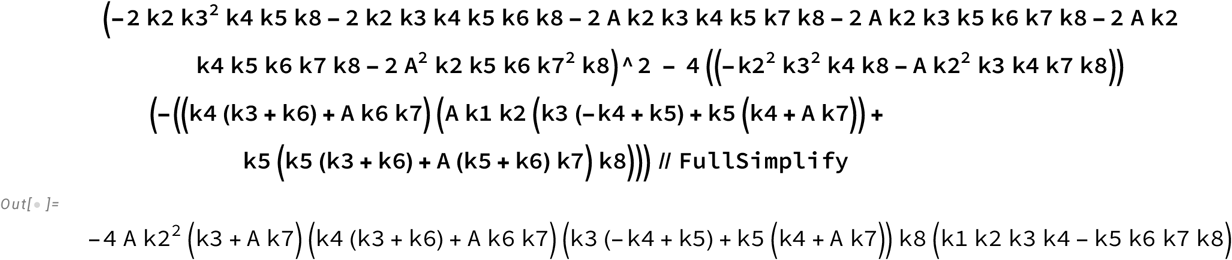

***In[* . ] : *=*** `Reduce`[`DisP1B` **>** **0**, **{**`k1, k2, k3, k4, k5, k6, k7, k8`**}**, **PositiveReals]**

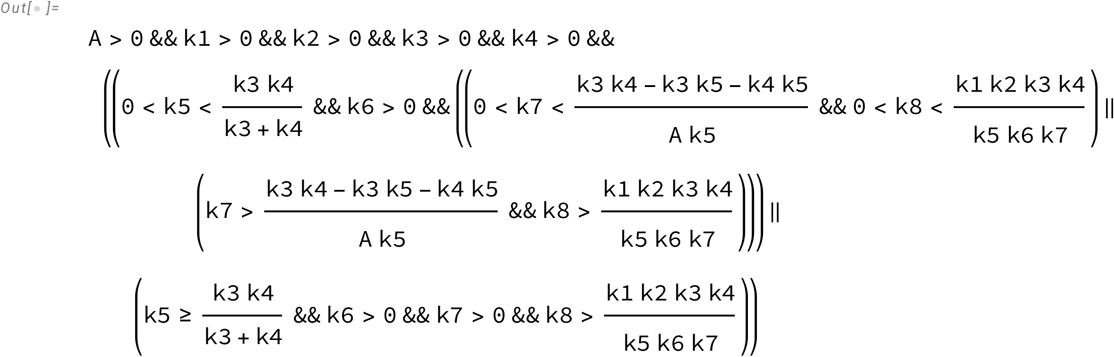

But from our requirement that *c* > 0, we have 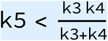 and 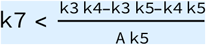. Therefore, the necessary condition is 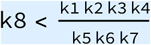

This implies that for non-monotonicity in B we will need the cycle to operate **clockwise** such that k8 k5 k6 k7 < k1 k2 k3 k4.

**<u>Thus P1 can exhibit non-monotonicity only with respect to one of the TFs.</u>**

## Promoter state 2 exhibits non-monotonicity with respect to one of the TFs only

Now, we look at conditions necessary for the non-monotonicity of *P*_2_ with A and B.

We look at 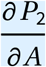 and the conditions for it to have a real root for some *A* > 0.

***In[* . ] : *=*** `DelP2A` **=** `D`[`P2, A`**] //** `FullSimplify`

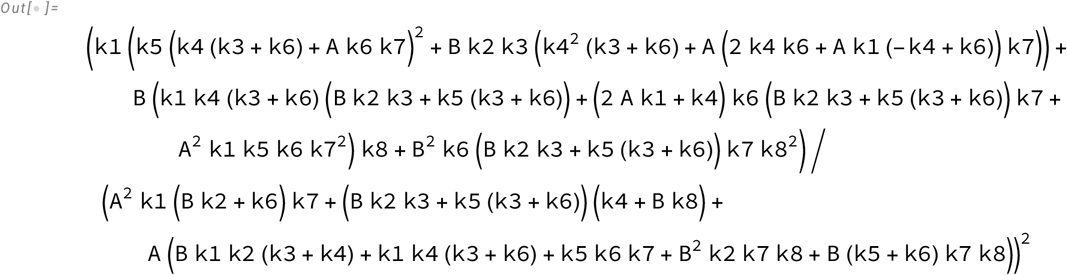

As before, the denominator is always positive. We look at the numerator

***In[* . ] : *=*** `NumDelP2A` **=** `Collect`[`Numerator`[`DelP2A`**] //** `Expand, A`]

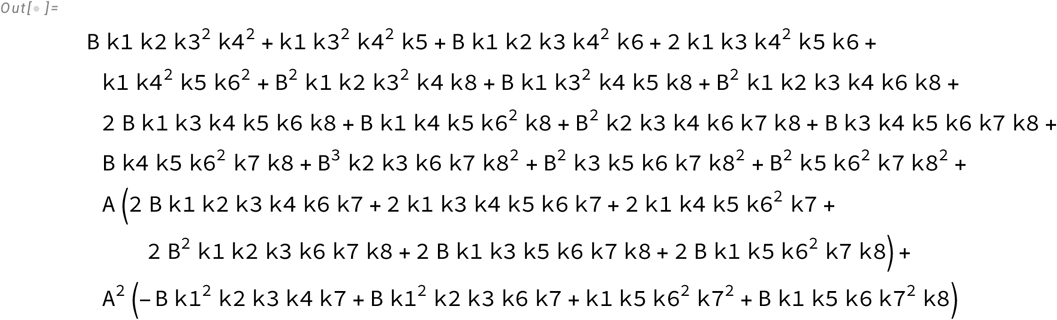

This is a quadratic in A, *a A*^2^ + *b A* + *c* with

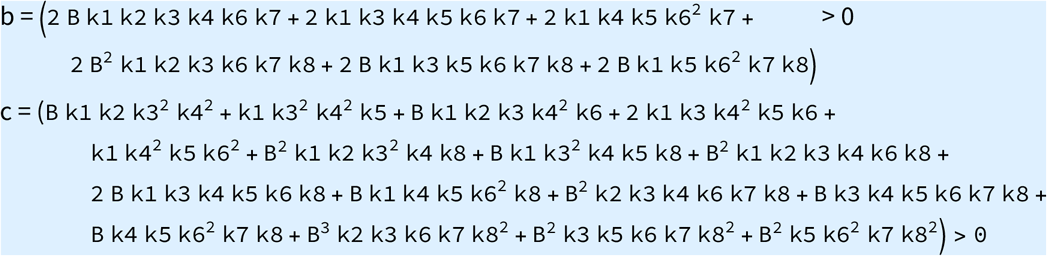

So for the partial derivative to have a real root for some *A* > 0, we need *a < 0* and *b*^2^ - 4 *a c* > 0. We find the conditions for *a* < 0. This is the necessary and sufficient condition that enforces *b*^2^ - 4 *a c* > 0.

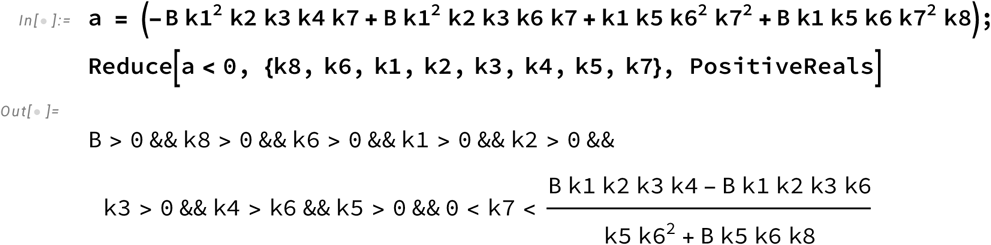

We have *k* 4 > *k* 6 along with some other conditions. Now we find conditions for *b*^2^ - 4 *a c* > 0.

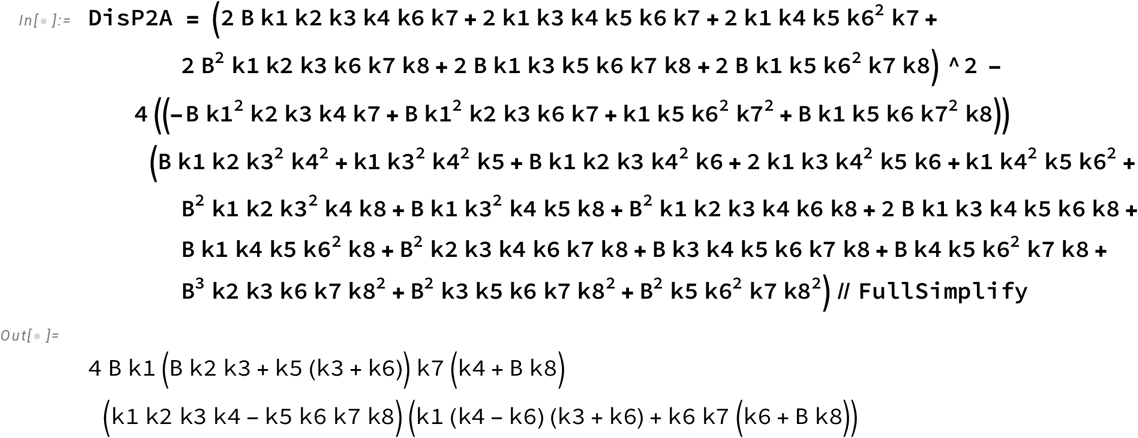

***In[* . ] : *=*** `Reduce`[`DisP2A` **>** **0**, **{**`k8, k6, k1, k2, k3, k4, k5, k7`**}**, `PositiveReals`]

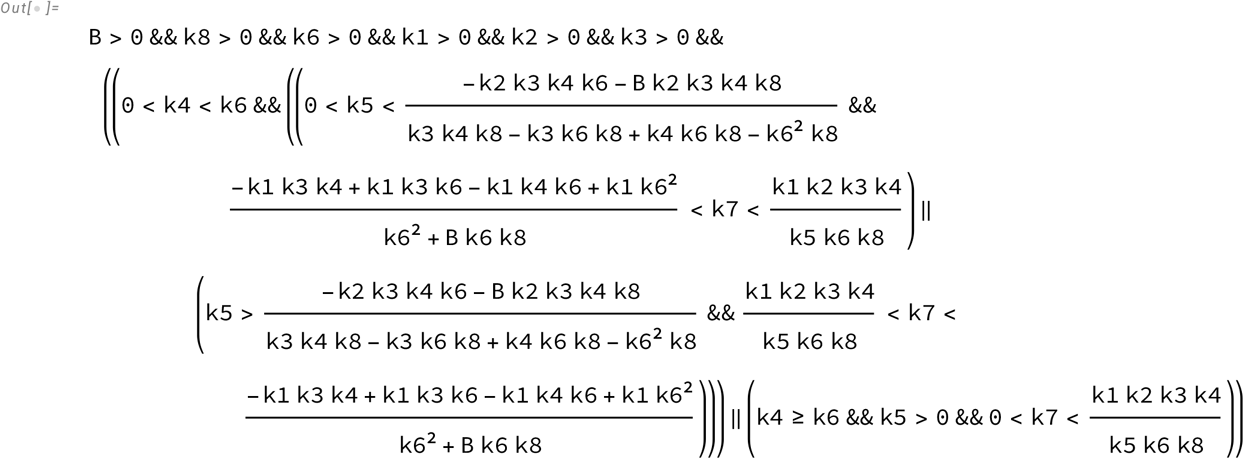

The conditions overlap when 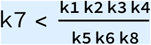. That is, k8 k7 k6 k5 < k1 k2 k3 k4. P2 can be non-monotonic in *A* only if the cycle is **clockwise.**

Next, we consider the case of 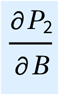. For non-monotonicity with *B*, we need a real root at some *B* > 0.

***In[* . ] : *=*** `DelP2B` **=** `D`[`P2, B`**] //** `FullSimplify`

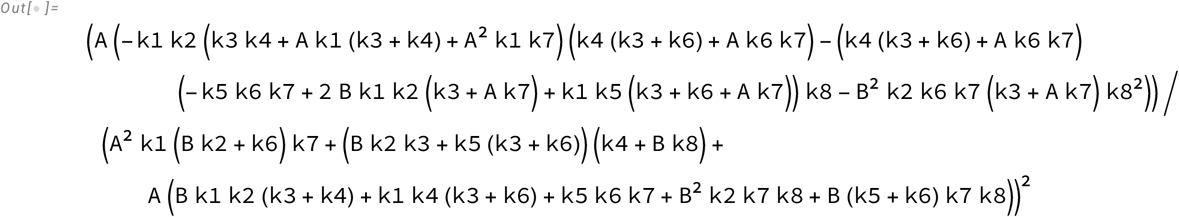

The denominator is always positive, the numerator is a quadratic in B given by *a B*^2^ + *b B* + *c*.

***In[* . ] : *=*** `NumDelP2B` **=** `Collect`[`Numerator`[`DelP2B`**] //** `Expand, B`]

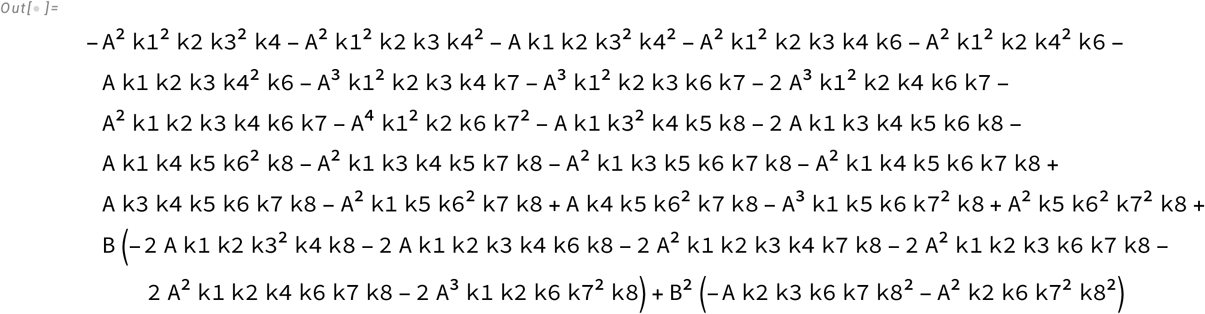

Note that here

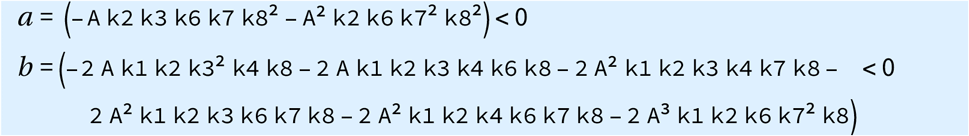

So for a real root at some *B* > 0, we need *c* > 0.

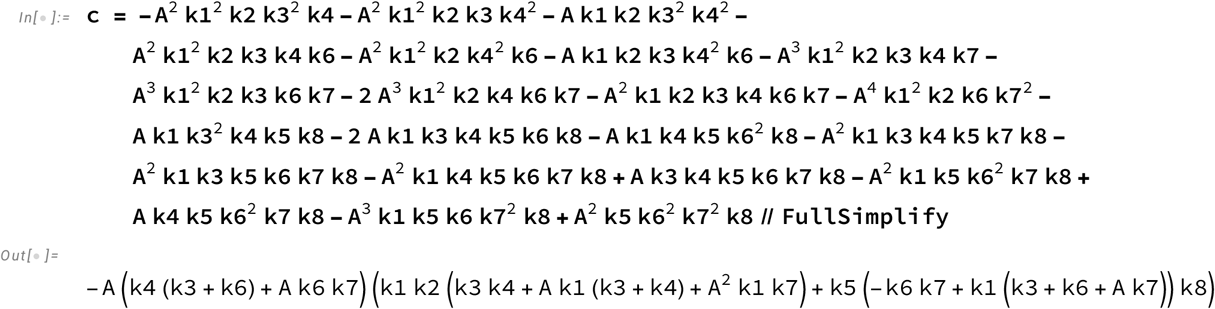

Conditions for *c* > 0 are

***In[* . ] : *=*** `Reduce`[`c` **>** **0**, **{**`k6, k7, k8, k1, k2, k3, k4, k5`**}**, **PositiveReals**]

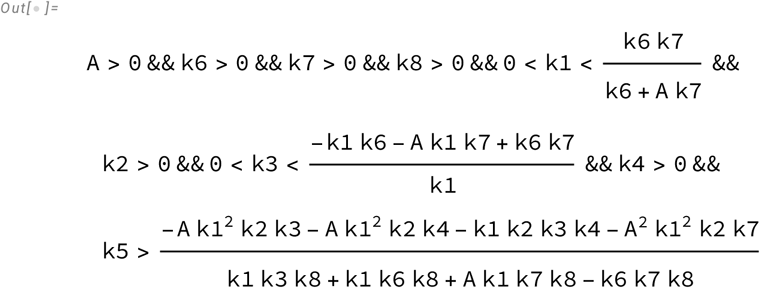

Now we look at *b*^2^ - 4 *a c* > 0

***In[* . ] : *=*** `DisP2B` **=**

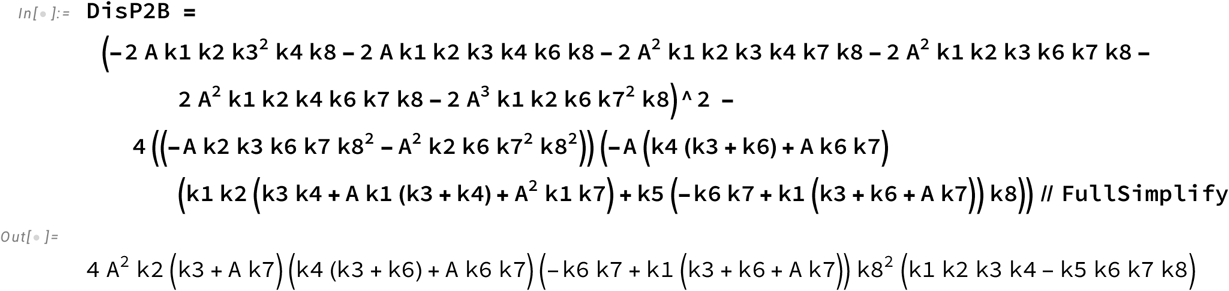

***In[* . ] : *=*** `Reduce`[`DisP2B` **>** **0**, **{**`k6, k7, k8, k1, k2, k3, k4, k5`**}**, **PositiveReals**]

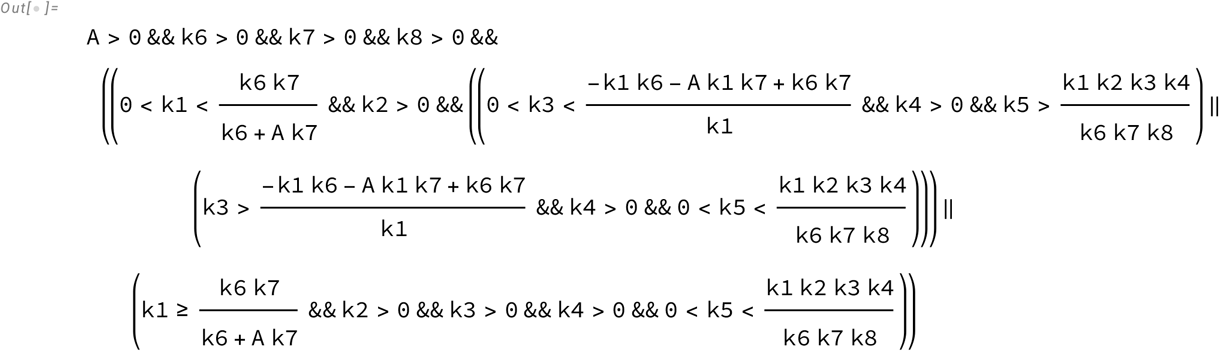

The overlapping conditions are met when 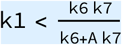 and 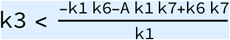 and 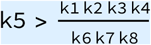.

For P2 to be non-monotonic with B we need k8 k7 k6 k5 > k1 k2 k3 k4

The cycle should be **anticlockwise**

**<u>P2 can exhibit non-monotonicity only with respect to one of the TFs.</u>**

## Promoter state 3 exhibits non-monotonicity with respect to one of the TFs only

The partial derivative 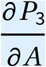*i*s

***In[* . ] : *=*** `DelP3A` **=** `D`[`P3, A`**] //** `FullSimplify`

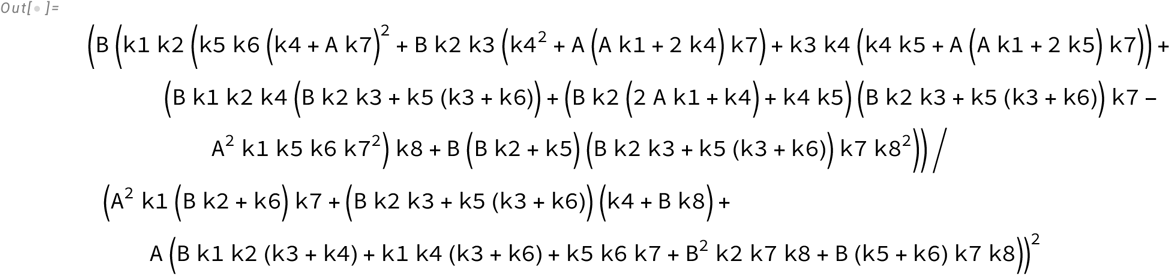

The denominator is always positive, on expanding the numerator as a quadratic in *A* as *a A*^2^ + *b A* + *c*,

***In[* . ] : *=*** `NumP3A` **=** `Collect`[`Numerator`[`DelP3A`**] //** `Expand, A`]

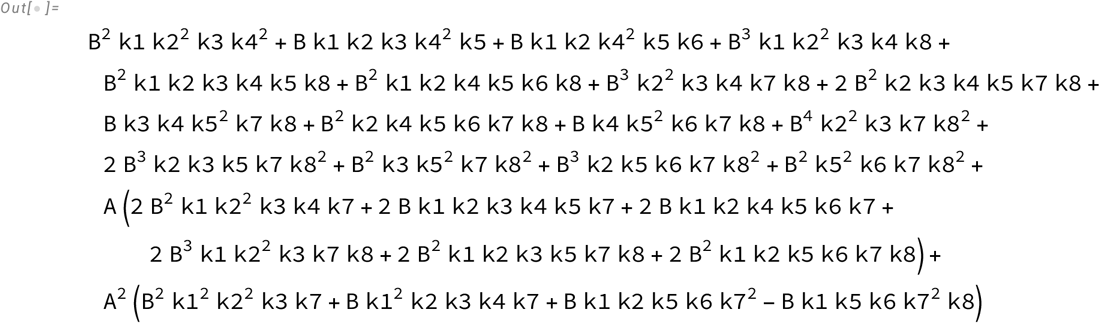

We get:

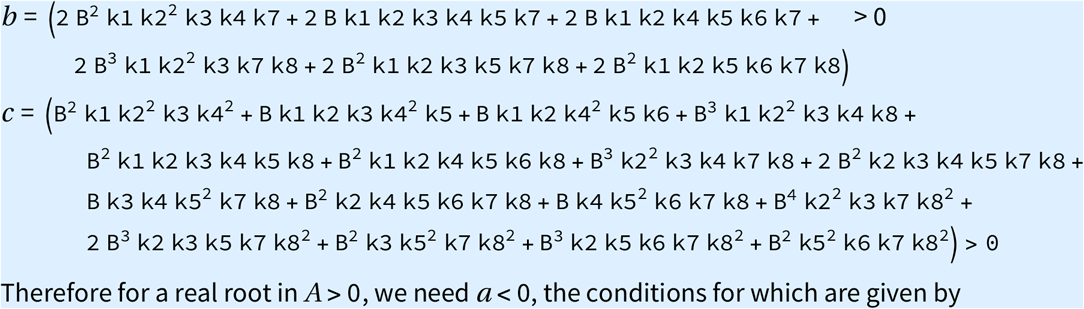

Therefore for a real root in *A* > 0, we need *a* < 0, the conditions for which are given by

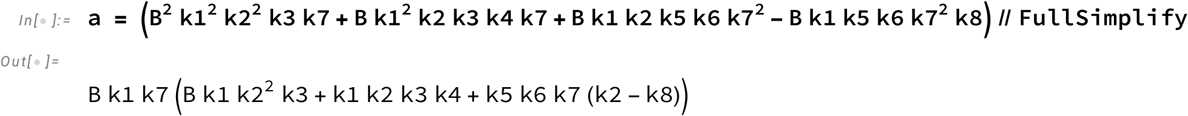

***In[* . ] : *=*** `Reduce`[`a` **<** **0**, **{**`k8, k1, k2, k3, k4, k5, k6, k7`**}**, **PositiveReals**]

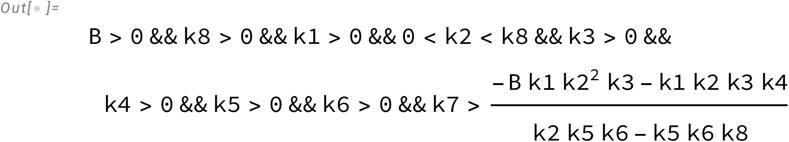

Now we look at *b*^2^ - 4 *a c* > 0 and find the conditions needed for it to be true.

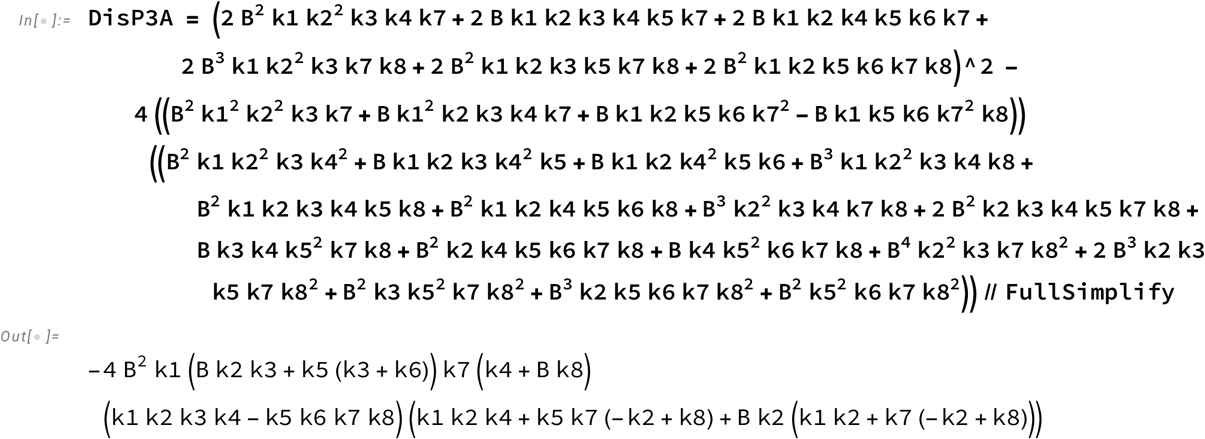

***In[* . ] : *=*** `Reduce`[`DisP3A` **>** **0**, **{**`k8, k1, k2, k3, k4, k5, k6, k7`**}**, **PositiveReals**]

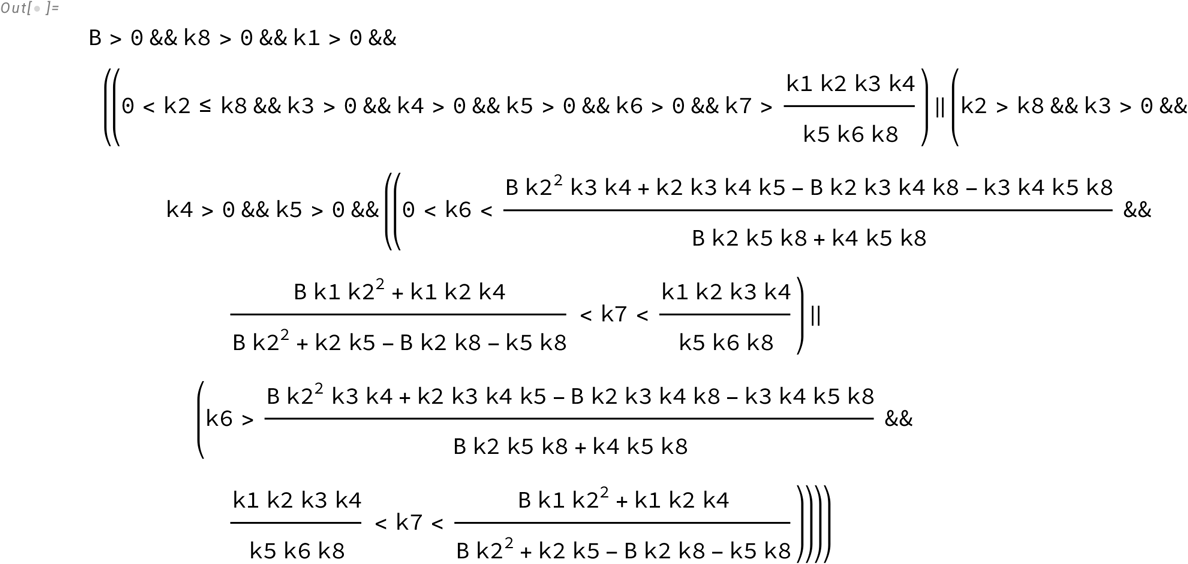

The overlapping conditions given *k* 2 < *k* 8 which is met only when 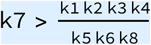. That is k8 k7 k6 k5 > k1 k2 k3 k4. *P*_3_ is non-monotonic with A only if the cycle is **Anticlockwise**.

Now we look at 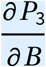. Note that the denominator is always positive.

***In[* . ] : *=*** `DelP3B` **=** `D`[`P3, B`**] //** `FullSimplify`

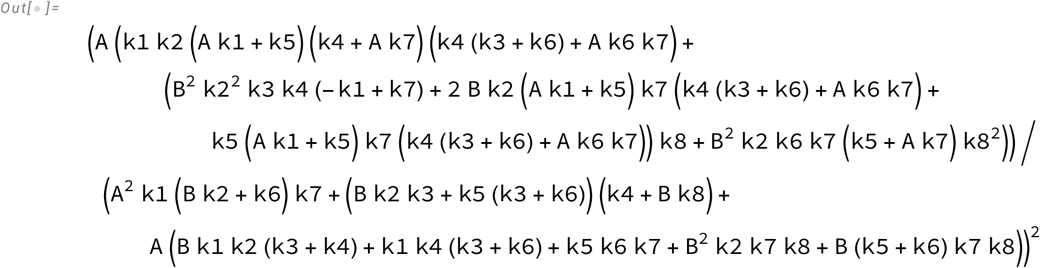

Taking the numerator and expanding it as a quadratic in B,

***In[* . ] : *=*** `NumDelP3B` **=** `Collect`[`Numerator`[`DelP3B`**] //** `Expand, B`]

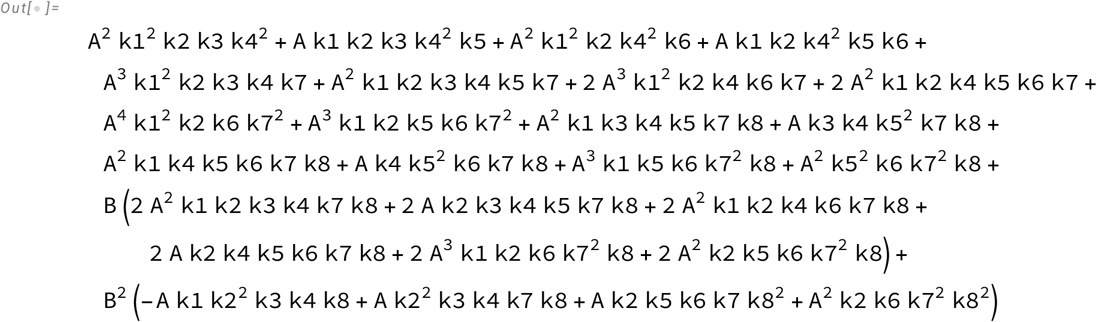

Here we have

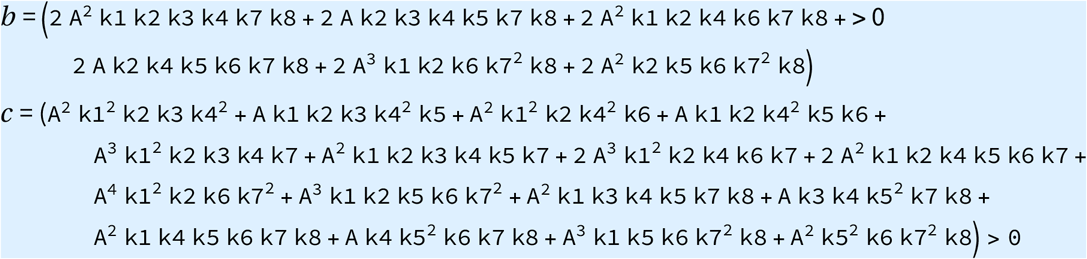

So for real roots in *B* > 0, we need a < 0. We find the conditions for this to hold.

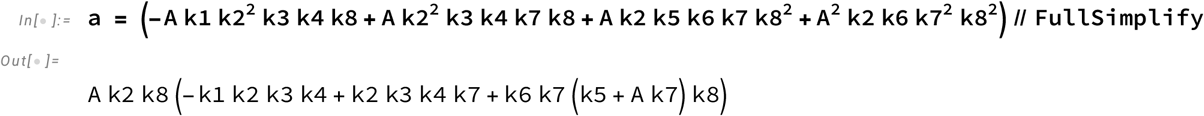

***In[* . ] : *=*** `Reduce`[`a` **<** **0**, **{**`k8, k7, k1, k2, k3, k4, k5, k6`**}**, **PositiveReals**]

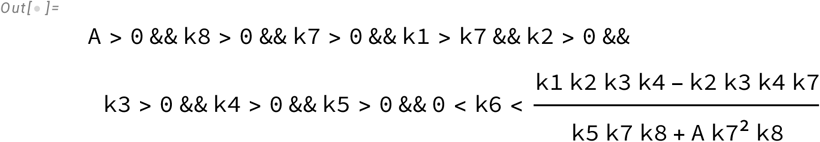

Now we look at the conditions for *b*^2^ - 4 *a c* > 0

***In[* . ] : *=*** `DisP3B` **=**

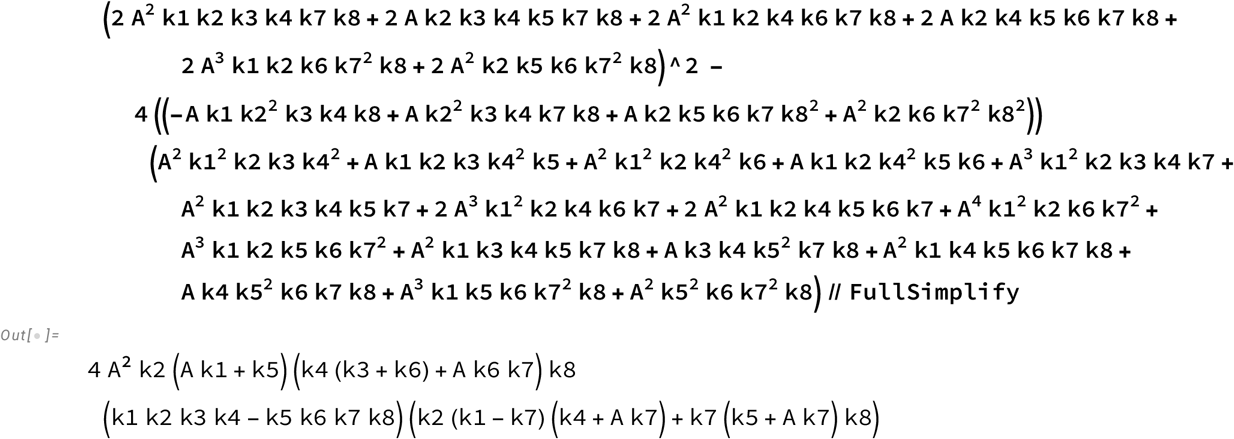

***In[* . ] : *=*** `Reduce`(`DisP3B` **>** **0**, **{**`k8, k7, k1, k2, k3, k4, k5, k6`**}**, **PositiveReals**)

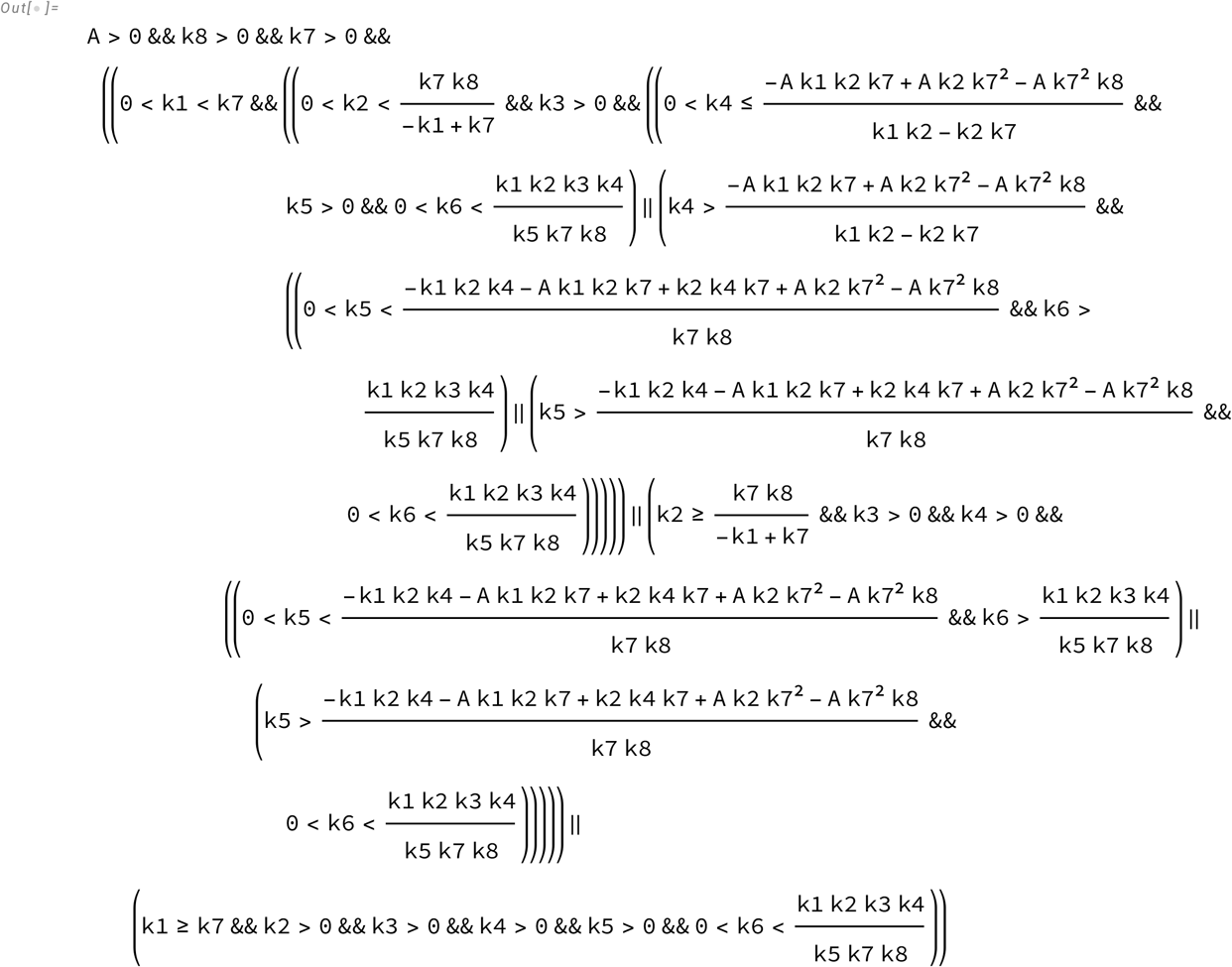

The overlapping conditions given by k1>k7 which is met only when 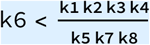. That is k8 k7 k6 k5 < k1 k2 k3 k4. P3 is non-monotonic with B only if the cycle is **Clockwise.**

**<u>P3 can exhibit non-monotonicity only with respect to one of the TFs.</u>**

## Promoter state 4 exhibits non-monotonicity with respect to one of the TFs only

We look at at 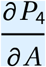.

***In[* . ] : *=*** `DelP4A` **=** `D`[`P4, A`**] //** `FullSimplify`

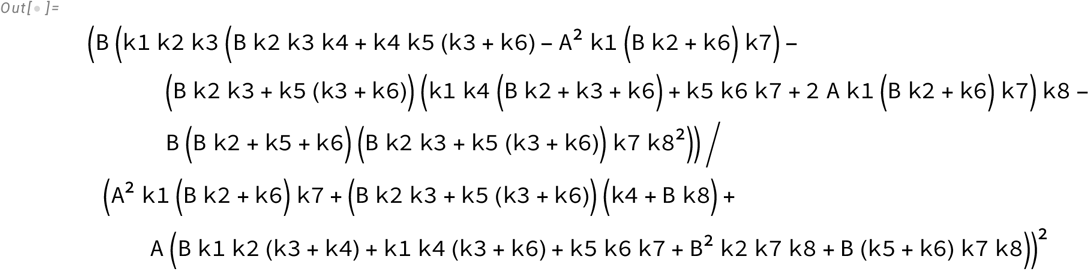

and consider the numerator. On expanding it as a quadratic in *A* in the form *a A*^2^ + *b A* + *c*, we get

***In[* . ] : *=*** `NumDelP4A` **=** `Collect`[`Numerator`[`DelP4A`**] //** `Expand, A`]

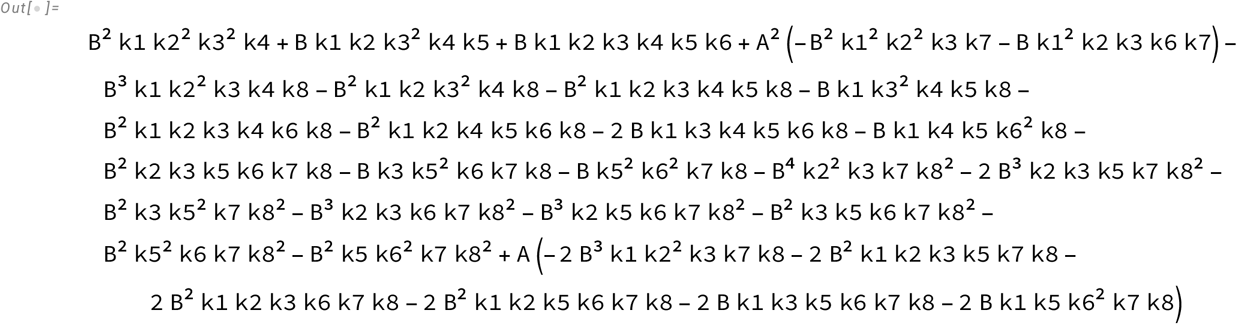

Where we have

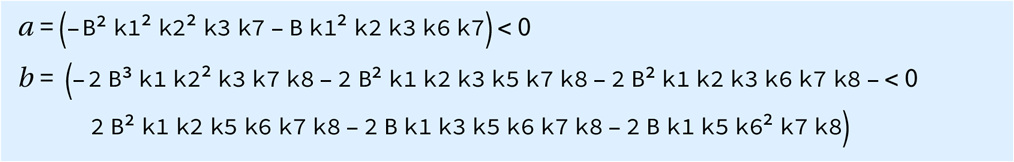

So for real roots in*A* > 0, we need *c* > 0.

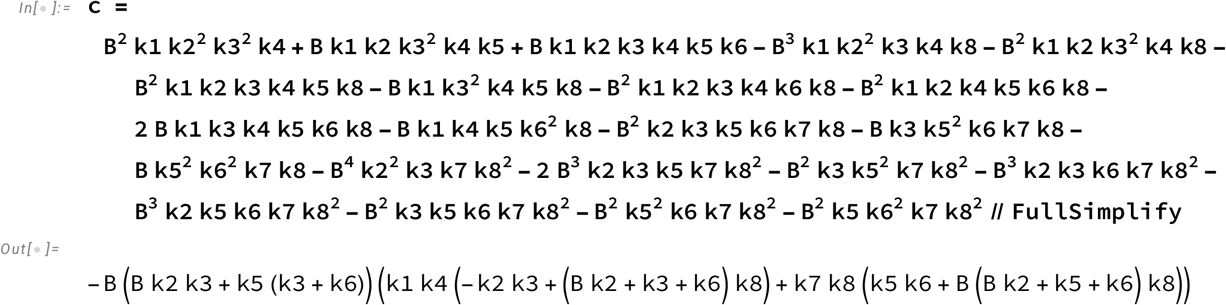

***In[* . ] : *=*** `Reduce`[`c` **>** **0**, **{**`k8, k1, k2, k3, k4, k5, k6, k7`**}**, **PositiveReals**]

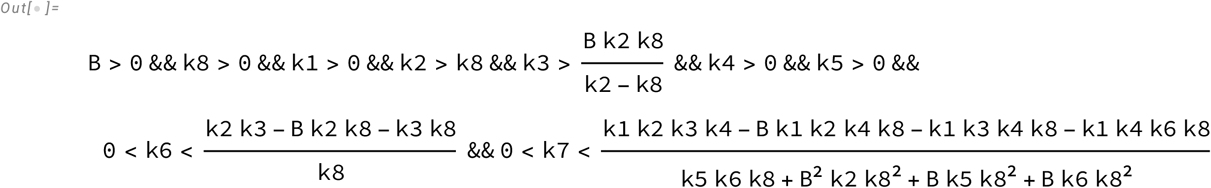

Now we look at *b*^2^ - 4 *a c* > 0

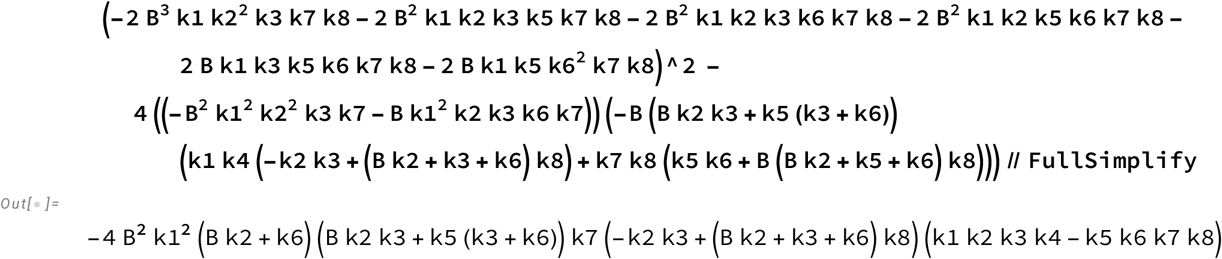

***In[* . ] : *=* `Reduce`[`DisP4A` > `0,` {`k8, k1, k2, k3, k4, k5, k6, k7`}`, PositiveReals`]**

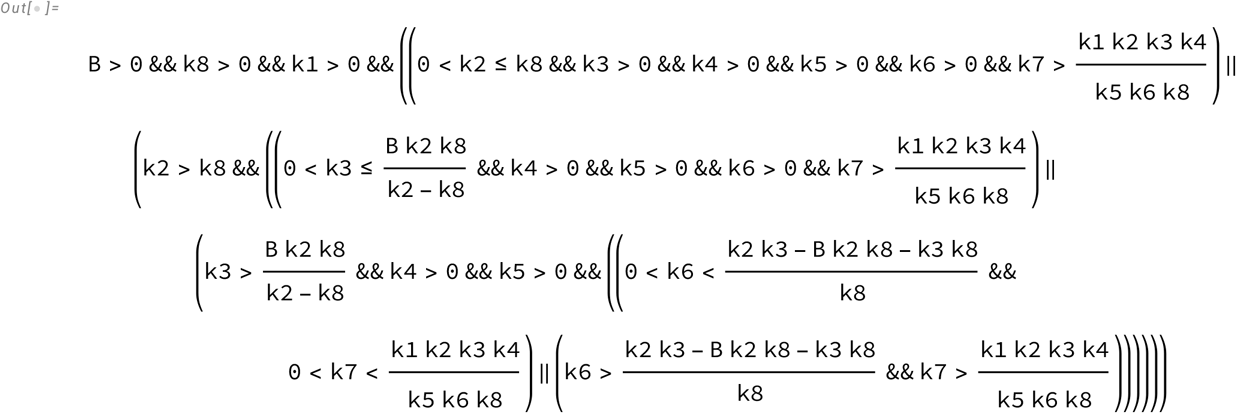

We get the overlap as *k* 2 > *k* 8 and 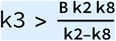 and 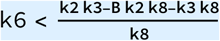 so we have 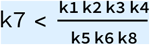.

This means k5 k6 k7 k8 < k1k2 k3 k4.

*P*_4_ can be non-monotonic with A when the cycle operates **clockwise**.

Now we look at 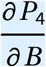.

***In[* . ] : *=*** `DelP4B` **=** `D`[`P4, B`**] //** `FullSimplify`

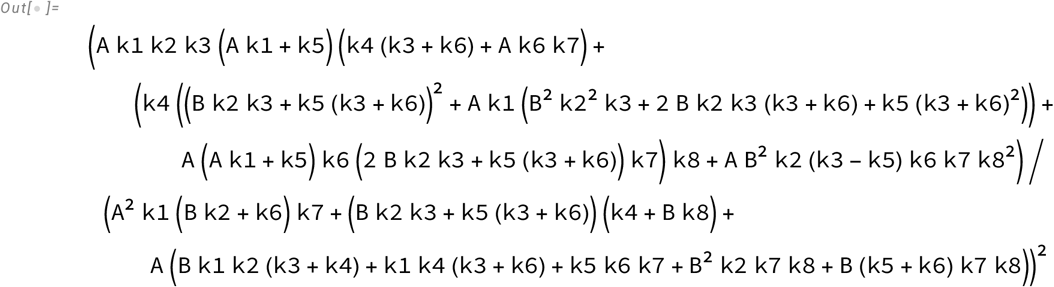

Expanding the numerator as a quadratic in *B* we get the form *a B*^2^ + *b B* + *c*

***In[* . ] : *=*** `NumP4B` **=** `Collect`[`Numerator`[`DelP4B`**] //** `Expand, B`]

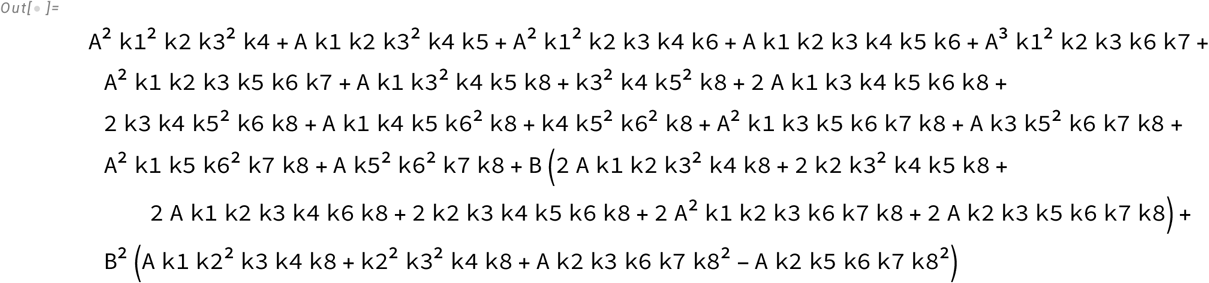

Where we have

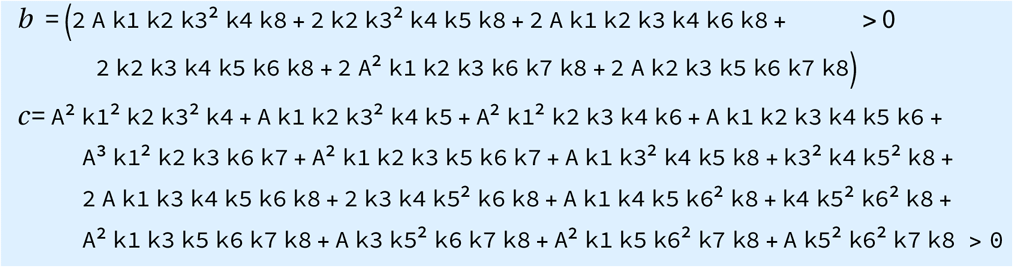

For a real root in *B* > 0, we need *a* < 0.

***In[* . ] : *=*** `a` **= (****A k1 k2**^**2**^ **k3 k4 k8** **+** **k2**^**2**^ **k3**^**2**^ **k4 k8** **+** **A k2 k3 k6 k7 k8**^**2**^ **-** **A k2 k5 k6 k7 k8**^**2**^**) //** `FullSimplify`

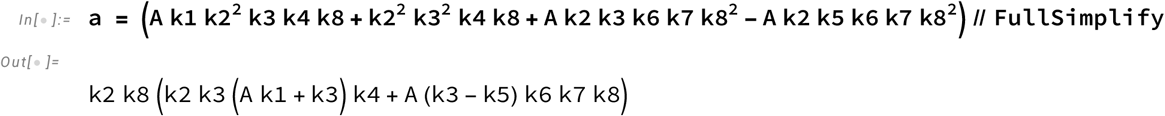

***In[* . ] : *=*** `Reduce`[`a` **<** **0**, **{**`k1, k2, k3, k4, k5, k6, k7, k8`**}**, **PositiveReals**]

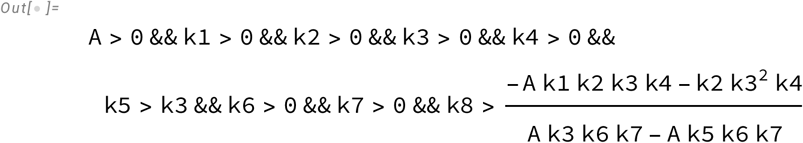

Now we look at *b*^2^ - 4 *a c* > 0

***In[* . ] : *=*** `DisP4A =`

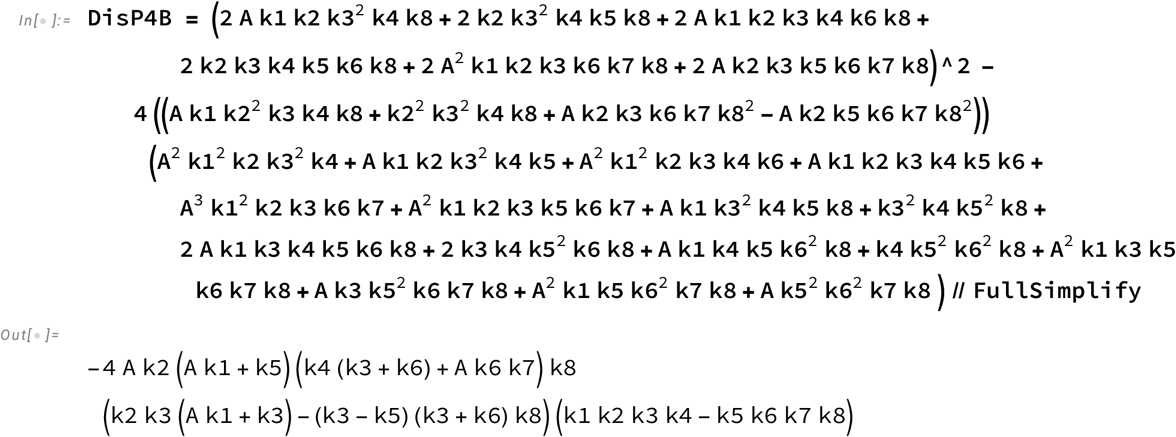

***In[* . ] : *=*** `Reduce[DisP4B > 0, {k1, k2, k3, k4, k5, k6, k7, k8}, PositiveReals]`

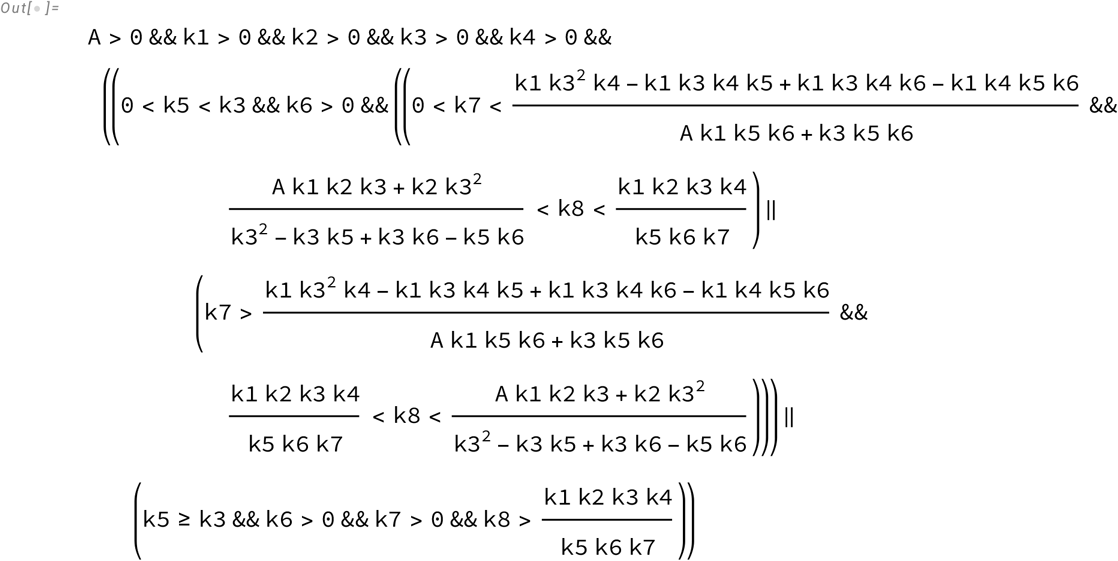

The overlapping space is where k5>k3, so we have 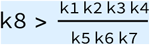. k8 k7 k6 k5 > k1 k2 k3 k4. For P4 to be non-monotonic with B, the cycle needs to be **Anticlockwise**

**<u>P4 can exhibit non-monotonicity only with respect to one of the TFs.</u>**

